# TREM2 agonist antibody rebuilds the resident synovial macrophage lining barrier in rheumatoid arthritis

**DOI:** 10.64898/2026.02.02.703430

**Authors:** Huihui Yuan, Jingyun Wang, Qichao Wang, Le Jiang, Leixin Liu, Weicheng Shen, Peixia Li, Wanting Liu, Ziyu Liu, Fanlei Hu, Xu Cai, Wanli Liu, Qiong Wu, Xiumei Wang, Hao Yan, Xiaodan Sun

## Abstract

Rheumatoid arthritis (RA) can be viewed as a disease of barrier failure, in which CX_3_CR1^+^/TREM2^+^ synovial tissue–resident macrophages that form the lining barrier over cartilage and bone become fragmented and disorganized. However, how to therapeutically rebuild this barrier—and how macrophage states transition during repair—remain unclear. We engineered TR-Ab19, a mouse-selective agonistic antibody against TREM2, as a precision tool to initiate and interrogate barrier repair *in vivo*. TR-Ab19 engages TREM2-linked downstream signaling and redirects synovial macrophages from Clec4d^+^ inflammatory/proliferative programs toward TREM2^+^CX_3_CR1^+^Aqp1^+^ barrier-like states, thereby rebuilding the lining barrier. Across collagen-induced arthritis (CIA) and serum-transfer arthritis (STA) models, TR-Ab19 reduces synovitis, preserves cartilage and bone microarchitecture, limits osteoclastogenesis, and attenuates systemic cytokines and B-cell abnormalities. Single-cell RNA-seq with trajectory and cell–cell communication analyses reveal a TREM2-dependent shift toward a barrier-dominant macrophage ecosystem. Together, these findings establish antibody-mediated reprogramming of resident synovial macrophages as a barrier-centered strategy for RA and provide a framework for instructing macrophage niches in chronic inflammation.

## Introduction

Rheumatoid arthritis (RA) affects ∼1% of the population and remains a major cause of pain, disability, and premature mortality despite biologics and targeted synthetic disease-modifying anti-rheumatic drugs (DMARDs) [1–2]. Genetic, serologic, and longitudinal studies support a multistage model in which mucosal endotypes, defective B-cell checkpoints, and evolving autoimmunity precede persistent synovitis and joint damage [3–7]. Yet many patients experience nonresponse, waning efficacy, residual pain, or ongoing “silent” structural progression [1,6–7], underscoring the need for mechanism-driven interventions that provide durable tissue protection.

Recent work has identified a protective synovial lining “barrier” formed by specialized synovial tissue macrophages (STMs), positioning the synovium as a barrier-like organ. This unit is formed by tissue-resident macrophages (TRMs) whose spatial organization shapes leukocyte flux and disease outcomes in RA [6,8–9]. Single-cell, fate-mapping, imaging, and atlas studies delineate STMs as heterogeneous populations comprising locally renewing TRMs and monocyte-derived inflammatory subsets with distinct roles in joint homeostasis, flares, and remission [9–13]. In healthy joints, CX_3_CR1^+^MerTK^+^TREM2^+^ lining-layer STMs form a continuous layer enriched for junctional components, TAM-family signaling, efferocytosis, and lipid-handling programs that seal the joint cavity and restrain leukocyte ingress [10– 11]. Longitudinal biopsy studies suggest that disruption of the MerTK^+^TREM2^+^ macrophage lining barrier precedes clinical onset, accompanies active RA, and can be partially restored during deep remission [12–13]; by contrast, inflammatory macrophage programs (e.g., HBEGF^+^, SLAMF7^high^, ETS2-driven) promote synovitis, fibroblast invasiveness, and joint destruction [14–16]. Although macrophage-focused approaches—including repolarization, responsive biomaterials, and transcriptional targeting— have shown promise in arthritis [17–28], explicitly reconstruction of the CX_3_CR1^+^TREM2^+^ lining barrier especially plasticity test *in vivo* of barrier-forming macrophage states have not been designed and executed. We hypothesized that selectively reprogramming resident lining STMs to re-establish barrier function is an underexploited therapeutic strategy to suppress synovial inflammation and restore joint homeostasis in RA.

Recent advances in antibody engineering enable precise, instructive modulation of immune cells— particularly macrophages—*in vivo* [29–34]. One paradigm is to target triggering receptor expressed on myeloid cells 2 (TREM2) on tissue-resident macrophages. In neurodegeneration models, TREM2 agonists can reprogram microglial states through SYK–PI3K–linked signaling and improve disease-relevant phenotypes, establishing TREM2 activation as a tractable route to reshape myeloid compartments [35–38]. Notably, synovial lining-layer macrophages that form the CX_3_CR1^+^TREM2^+^ barrier consistently express TREM2 [10–13]. These observations motivate a barrier-repair strategy that directly reprograms TREM2^+^ lining macrophages—an opportunity not systematically addressed by current RA therapies [31–36].

Here, we integrate concepts from RA pathogenesis [1–7], synovial macrophage biology [9–13], and TREM2-targeted immunotherapy [35–38] to develop a barrier reconstruction–centered strategy for macrophage state reprogramming. We hypothesized that TREM2 agonism could selectively rewire synovial tissue–resident macrophages toward barrier-like states and restore the lining barrier as a therapeutic endpoint. We generated TR-Ab19, a mouse-selective TREM2 agonist antibody (Scheme 1A), and tested it across collagen-induced arthritis, CX_3_CR1 lineage tracing, and K/BxN serum-transfer arthritis, combined with multimodal cellular profiling and single-cell transcriptomics. TR-Ab19 reduced synovitis, preserved cartilage and bone microarchitecture, and limited osteoclastogenesis, while dampening systemic cytokines and germinal-center–associated B-cell responses. Mechanistically, TR-Ab19 rebuilt a continuous CX_3_CR1^high^/TREM2^+^ lining barrier and shifted macrophage trajectories from Clec4d^+^ inflammatory/cycling programs toward CX_3_CR1^high^/Aqp1^+^ barrier-like and reparative states, coincident with reduced leukocyte influx and restoration of a barrier-dominant macrophage ecosystem (Scheme 1B). Together, these findings establish antibody-guided reprogramming of resident synovial macrophages as a barrier-centered therapeutic concept for RA.

## Results

### TR-Ab19 is a high-affinity agonistic antibody that selectively binds and activates mouse TREM2

To enable barrier-centered, cell-state reprogramming of synovial lining-layer macrophages in RA, we generated an agonistic monoclonal antibody against mouse TREM2 using a hybridoma screening funnel that integrated binding and function. Mice were immunized with a KLH-conjugated mouse TREM2 peptide epitope, hybridomas were produced, and sequential rounds of ELISA- and activity-guided triage enriched clones with robust recombinant mouse TREM2 (rmTREM2) engagement and agonistic potential (Scheme 1A; Fig. 1A; Supplementary Methods). This multistep workflow narrowed an initial pool to a small set of IgG candidates for head-to-head comparison, from which TR-Ab19 was selected as the lead over an earlier in-house antibody (TR-Ab1) (Scheme 1A; Fig. 1A-D).

**Fig. 1.**
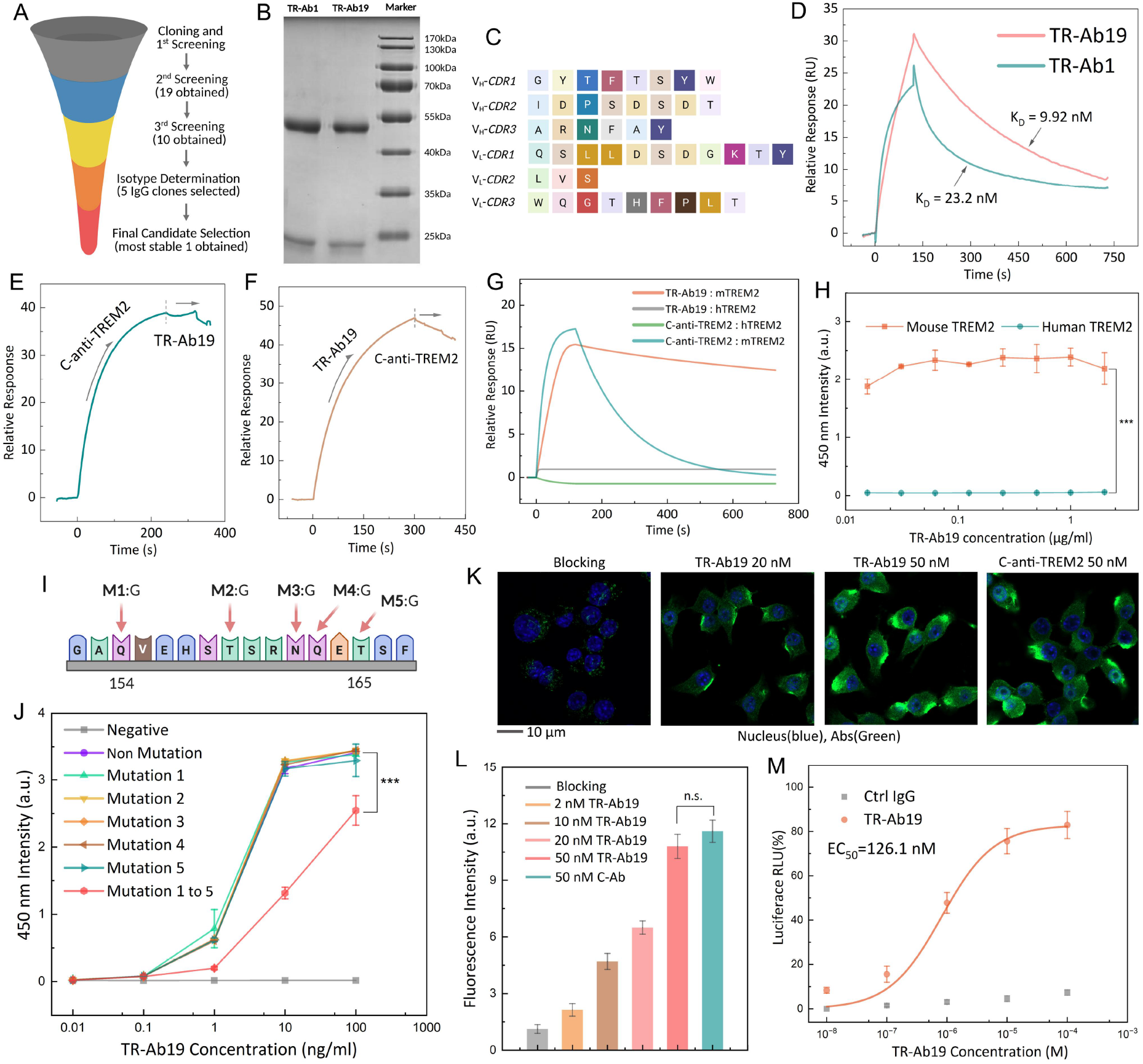
Generation and in vitro characterization of TR-Ab19, a mouse-selective TREM2 agonist antibody. (A) Binding-plus-function screening nominating TR-Ab19 (lead) and TR-Ab1. (B) Reducing SDS–PAGE of purified antibodies. (C) TR-Ab19 heavy- and light-chain CDR sequences. (D) SPR sensorgrams and kinetic fitting for TR-Ab1/TR-Ab19 binding to recombinant mouse TREM2 (rmTREM2). (E–F) SPR competition with a commercial anti-TREM2 antibody, consistent with partially overlapping/sterically coupled epitopes. (G–H) Species selectivity by SPR/ELISA, showing rmTREM2 binding but minimal binding to recombinant human TREM2 (rhTREM2). (I–J) Epitope localization by mutational mapping. (K–L) Confocal imaging of RAW264.7 macrophages showing dose-dependent binding to endogenous TREM2. (M) NFAT–luciferase assay demonstrating dose-dependent agonistic activity (n = 3; mean ± SD). Abbreviations: CDR, complementarity-determining region; ELISA, enzyme-linked immunosorbent assay; NFAT, nuclear factor of activated T cells; SPR, surface plasmon resonance.

To establish a genetically defined reagent, we cloned the TR-Ab19 VH and VL sequences from the parental hybridoma. RNA quality control, RT–PCR amplification, colony screening, and sequencing confirmed single, clean VH/VL products with consistent inserts across clones, defining the TR-Ab19 variable regions and CDR composition (Fig. S1). Recombinant TR-Ab19 was then expressed and purified, yielding the expected heavy- and light-chain bands and high purity by SDS–PAGE and immunoblotting (Fig. 1B). The CDR sequences of TR-Ab19 are shown in Fig. 1C. Surface plasmon resonance (SPR) showed that TR-Ab19 and TR-Ab1 both recognized immobilized rmTREM2, whereas TR-Ab19 displayed slower dissociation and higher apparent affinity (K_D_ = 9.92 nM) (Fig. 1D). A commercial anti-TREM2 antibody (C-anti-TREM2) was selected for subsequent analysis and comparison. Sequential-injection competition SPR showed reciprocal interference between TR-Ab19 and the commercial anti-TREM2 antibody, consistent with steric competition and partially overlapping epitopes rather than identical binding sites (Fig. 1E–F and Fig. S1F). ELISA and SPR showed that TR-Ab19 binds mouse TREM2 with saturable kinetics but shows minimal binding to human TREM2 (Fig. 1G–H and Fig. S1F). Sequence-guided mutational mapping localized the TR-Ab19 epitope on mouse TREM2 (Fig. 1I and Fig. S1F). We combined structure-guided prioritization with alanine-scanning mutagenesis of a surface-exposed ectodomain segment (aa 154–165). While single substitutions had minimal effects, simultaneous alanine substitution of the targeted residues nearly abolished binding by ELISA, supporting a conformational epitope centered on this loop (Fig. 1I-J). Epitope mapping places the TR-Ab19 footprint near the reported ADAM10/17 cleavage region, suggesting a potential steric contribution in addition to agonism; this hypothesis requires direct experimental validation (Fig. 1I and Fig. S1F). Furthermore, we tested whether TR-Ab19 functions as a bona fide agonist. TR-Ab19 recognized endogenous TREM2 on macrophages. In RAW264.7 cells, confocal microscopy of RAW264.7 cells revealed clear, dose-dependent membrane staining with TR-Ab19 (absent for isotype control), consistent across the concentration range tested in the main and supplementary datasets (Fig. 1K – L; Fig. S2). TR-Ab19 induced a strong, concentration-dependent increase in fluorescence intensity, whereas control IgG was inactive; dose–response fitting yielded an EC_50_ of ∼126.1 nM (Fig. 1M).

Together, these data establish TR-Ab19 as a sequence-defined, mouse-selective, high-affinity TREM2 agonistic antibody that engages native macrophage TREM2 and activates downstream signaling— providing the molecular foundation for subsequent *in vivo* studies testing whether TREM2-driven state reprogramming can rebuild the STMs barrier and restrain RA pathogenesis.

### TR-Ab19 ameliorates collagen-induced arthritis (CIA) and preserves joint structure

We next asked whether pharmacologic TREM2 activation could protect joints *in vivo*. We used collagen-induced arthritis (CIA) in C57BL/6 mice as a chronic mouse model of RA. CIA was induced by type II collagen immunization on days 0 and 21, and TR-Ab19 was delivered intra-articularly at early disease (day 18) followed by an intraperitoneal dose one week after the booster (day 28). Healthy, CIA, CIA + TR-Ab19, and Trem2−/− (TREM2 KO) CIA mice were followed longitudinally and analyzed on day 56 (Fig. 2A).

**Fig. 2.**
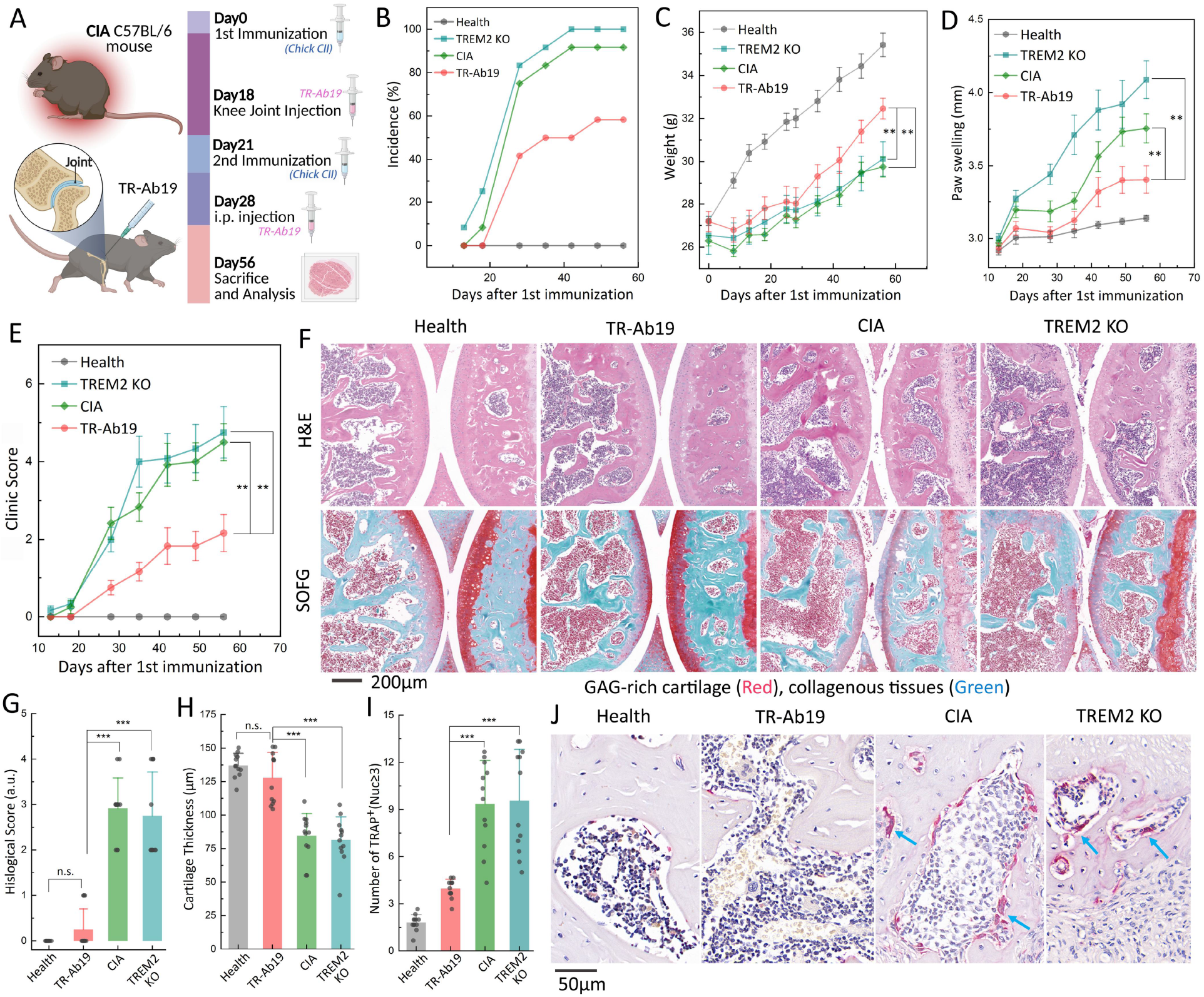
TR-Ab19 ameliorates collagen-induced arthritis and protects joint structure in a TREM2-dependent manner. (A) CIA study design and dosing schedule (Supplementary Methods). (B–E) Longitudinal arthritis incidence, body weight, paw swelling, and clinical score (scoring criteria in Supplementary Methods). (F) Representative ankle-joint histology (H&E and Safranin O–Fast Green; scale bar, 200 μm; related analyses in fig. S4–S5). (G) Histopathology scores (criteria in Supplementary Methods; related data in fig. S6). (H) Cartilage thickness quantified from Safranin O–Fast Green sections (μm). (I–J) Osteoclastogenesis assessed by TRAP staining: TRAP^+^ osteoclasts per field (defined as TRAP^+^ cells with ≥3 nuclei) and representative images (scale bar, 50 μm; related data in Fig. S7–S9). Data are shown as mean ± SEM unless otherwise stated. n values are indicated in the plots (each dot represents one mouse unless specified). Statistical tests follow the Statistical analysis section in Supplementary Methods; *P < 0.05, **P < 0.01, ***P < 0.001; n.s., not significant. Abbreviations: CIA, collagen-induced arthritis; TRAP, tartrate-resistant acid phosphatase.

TR-Ab19 markedly attenuated arthritis onset and progression. Compared with untreated CIA mice, TR-Ab19–treated animals showed a lower cumulative incidence of arthritis, improved body-weight trajectories, and significantly decreased paw swelling and clinical scores throughout the observation period (Fig. 2B–E). In contrast, TREM2 KO mice developed more severe disease, with higher clinical scores and greater paw swelling than wild-type CIA mice (Fig. 2B–E), indicating that endogenous TREM2 signaling restrains chronic joint inflammation and that TR-Ab19 amplifies this protective pathway.

Histologic analysis corroborated the clinical benefit and revealed robust structural protection. H&E staining of ankle joints from CIA and TREM2 KO mice showed marked synovial hyperplasia, dense inflammatory infiltrates, pannus formation, and extensive cartilage erosion, in sharp contrast to the smooth articular surfaces and preserved architecture in healthy controls (Fig. 2F; Figs. S3–S4). Safranin O–Fast Green staining demonstrated substantial loss of glycosaminoglycan content and thinning/discontinuity of articular cartilage in CIA and TREM2 KO joints, whereas TR-Ab19–treated joints retained intense Safranin O staining and relatively thick, continuous cartilage layers approaching those of healthy mice (Fig. 2F; Figs. S4–S5). Quantitative histomorphometry confirmed that TR-Ab19 significantly reduced composite histologic arthritis scores and partially restored cartilage thickness compared with untreated CIA and TREM2 KO groups (Fig. 2G–H; Fig. S6A–B).

Because osteoclasts are principal mediators of bone erosion in RA, we next examined TRAP-positive cells along subchondral and trabecular bone surfaces. CIA and especially TREM2 KO joints displayed abundant multinucleated TRAP^+^ osteoclasts concentrated at erosion fronts, consistent with aggressive osteoclastogenesis. In contrast, TR-Ab19–treated joints showed a marked reduction in TRAP^+^ multinucleated osteoclasts, approaching the sparse distribution seen in healthy controls (Fig. 2I–J; Figs. S6C, S7–S9). Quantification confirmed a significant decrease in osteoclast burden after TR-Ab19, whereas CIA and TREM2 KO groups remained elevated (Fig. 2I; Fig. S6C). Collectively, these data demonstrate sustained clinical and structural benefit from agonistic TREM2 activation in CIA, while the exacerbated phenotype in TREM2-deficient mice underscores the importance of TREM2 signaling in joint homeostasis.

### TR-Ab19 preserves bone microarchitecture, dampens systemic inflammation, and is superior to a commercial TREM2 antibody

Three-dimensional micro–computed tomography (micro-CT) reconstructions confirmed bone destruction in CIA and TREM2-KO CIA, whereas TR-Ab19 preserved surface architecture and trabecular integrity (Fig. 3A, Fig. S10; acquisition/segmentation parameters in Supplementary Methods). Quantitative micro-CT confirmed that multiple indices of cortical and trabecular integrity—including Cr.BV, Tb.BV, Tb.N, and Tb.Th—were significantly reduced in CIA and especially deteriorated in TREM2 KO CIA mice, whereas TR-Ab19 partially or largely rescued each parameter (Fig. 3B). Analysis of distal tibial metaphyses similarly revealed decreased Tb.BS and increased Tb.Sp in CIA and TREM2 KO mice; TR-Ab19 tended to normalize these parameters (Fig. S11). Together with reduced TRAP^+^ osteoclast burden and improved histologic scores, these data indicate that TREM2 activation by TR-Ab19 protects the bony scaffold of arthritic joints.

**Fig. 3.**
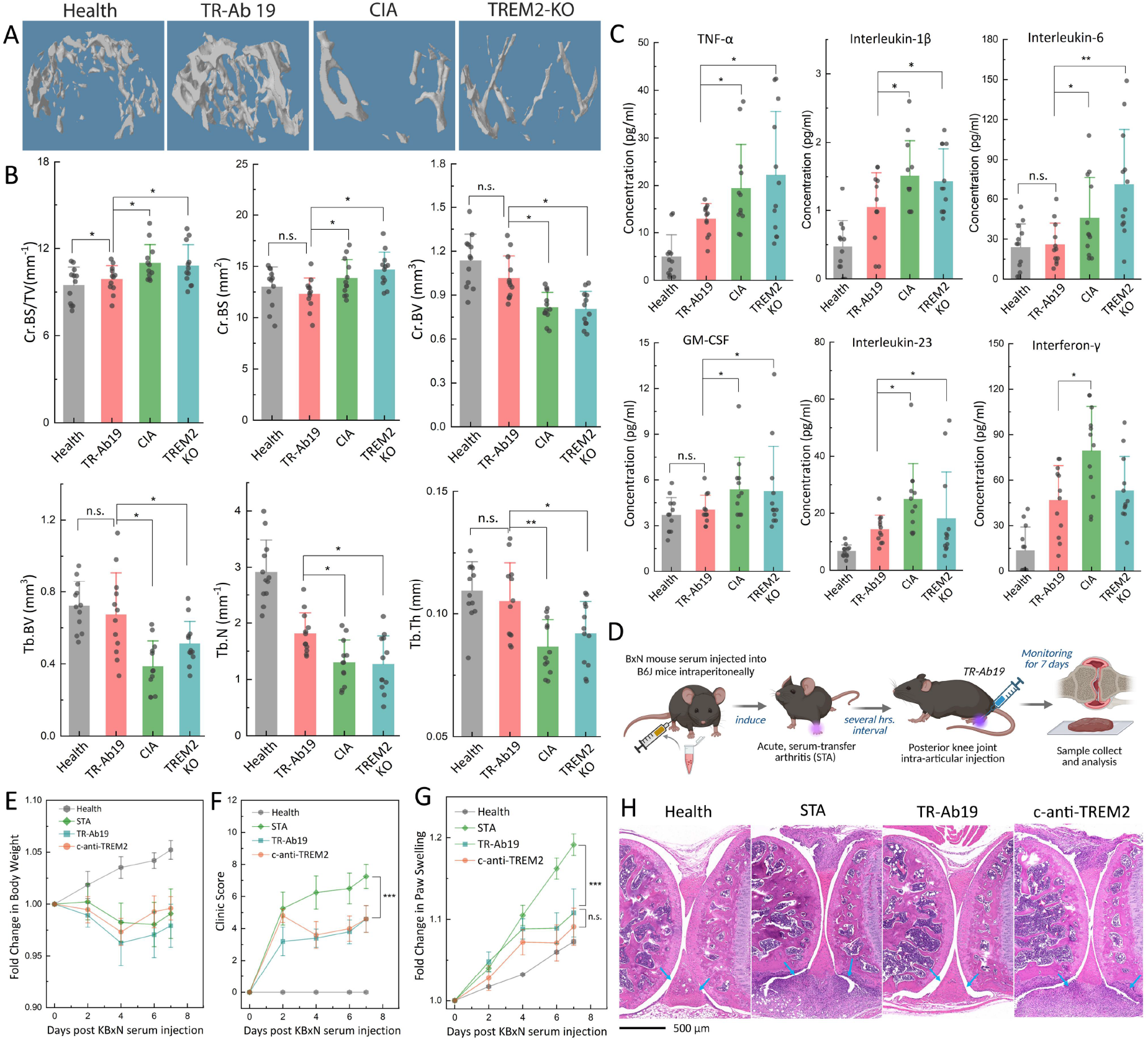
TR-Ab19 protects bone microarchitecture in CIA and improves disease in serum-transfer arthritis. (A) Representative 3D micro-CT reconstructions of knee joints from indicated groups (acquisition/segmentation parameters in Supplementary Methods; related data in Fig. S10). (B) Quantification of micro-CT structural parameters (related data in Fig. S11). (C) Serum cytokine profiling showing attenuation of inflammatory cytokines with TR-Ab19 (units as reported; extended panel in fig. S12). (D) Serum-transfer arthritis (STA) design with intra-articular TR-Ab19 or C-anti-TREM2. (E–G) Longitudinal body weight, clinical score, and paw swelling in STA (scoring criteria in Supplementary Methods). (H) Representative H&E staining of STA ankle joints (scale bars as indicated; n values and scoring criteria in Supplementary Methods). Data are shown as mean ± SEM unless otherwise stated; n values are indicated in the plots. Statistical tests follow Supplementary Methods; *P < 0.05, **P < 0.01, ***P < 0.001; n.s., not significant. Abbreviations: micro-CT, micro–computed tomography; STA, serum-transfer arthritis.

Because structural damage reflects both local and systemic inflammation, we next profiled circulating cytokines in the CIA cohorts. Multiplex serum analysis showed marked elevation of pro-inflammatory mediators, including TNF-α, IL-1β, IL-6, GM-CSF, IL-23, and IFN-γ, in CIA and TREM2 KO CIA mice compared with healthy controls. TR-Ab19 treatment significantly reduced several of these cytokines toward baseline, indicating that TREM2 agonism attenuates systemic inflammatory signaling (Fig. 3C). An extended panel revealed increases in additional pro-inflammatory (IL-2, IL-17, IL-12p70) and dysregulated Th2/regulatory cytokines (IL-4, IL-5, IL-10, IL-13, IL-18, IL-22, IL-27) in CIA and TREM2 KO mice, whereas TR-Ab19 generally shifted these broader perturbations toward healthy levels (Fig. S12), consistent with reduced osteoclastogenesis and preserved bone mass.

Based on the above, we further used serum-transfer arthritis (STA), a short-term arthritic mice model, to compare the therapeutic potential of TR-Ab19 to commercial TREM2 antibodies. B6 mice received arthritogenic K/BxN serum followed by intra-articular injection of TR-Ab19 or a commercial anti-TREM2 antibody (C-anti-TREM2) and were monitored for 7 days (Fig. 3D). Vehicle-treated STA mice developed rapid weight loss, high clinical scores, and prominent paw swelling, whereas TR-Ab19–treated mice showed attenuated weight loss, significantly lower arthritis scores, and reduced paw edema (Fig. 3E–G). Histopathology of STA ankle joints corroborated these findings: vehicle-treated joints exhibited synovial hyperplasia, dense inflammatory infiltrates, pannus, and early cartilage erosion, whereas TR-Ab19– treated joints showed reduced inflammatory pathology and better-preserved cartilage; the commercial antibody showed a smaller effect under the same conditions (Fig. 3H). Together with the CIA data, these results indicate that TR-Ab19 confers structural and anti-inflammatory protection in both chronic and acute arthritis settings and, in this study, outperformed the commercial anti-TREM2 antibody under matched dosing and readouts.

### TR-Ab19 restores the CX_3_CR1^+^ synovial macrophage barrier in a TREM2-dependent manner

Building on its structural and anti-inflammatory protection, we next asked whether TR-Ab19 acts by rebuilding the synovial CX_3_CR1^+^TRMs barrier. CIA was induced in *CX*_*3*_*CR1*^*Cre*^ R26-tdTomato (CIA-tdTomato) mice, which then received intra-articular and intraperitoneal TR-Ab19 on the same schedule as wild-type CIA (Fig. 4A). As in earlier cohorts, TR-Ab19 substantially ameliorated disease, lowering clinical scores and arthritis incidence, limiting paw swelling, and preserving body weight compared with untreated CIA-tdTomato controls (Fig. 4B–E).

**Fig. 4.**
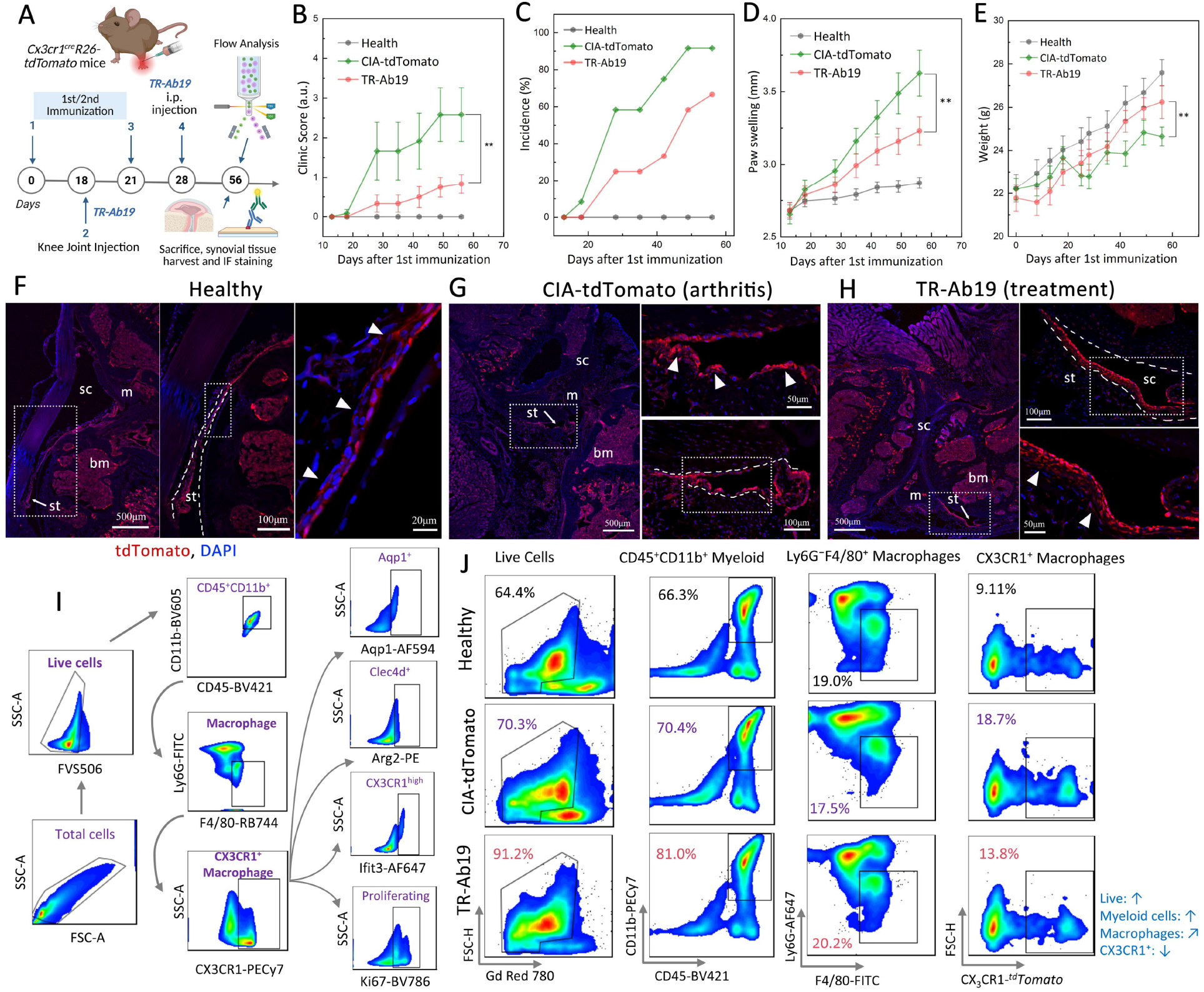
TR-Ab19 restores the CX_3_CR1^+^ synovial macrophage lining barrier in lineage-tracing CIA mice. (A) Study design for CIA in *CX*_*3*_*CR1*^*cre*^R26-tdTomato (CIA-tdTomato) mice with TR-Ab19 dosing and endpoint analyses. (B–E) Time courses of clinical score, arthritis incidence, paw swelling, and body weight in Healthy, CIA-tdTomato, and TR-Ab19–treated CIA-tdTomato cohorts. (F–H) Confocal imaging of knee joints showing tdTomato^+^ CX_3_CR1-lineage macrophages (red) and nuclei (DAPI, blue): continuous lining organization in Healthy joints, disruption in CIA, and partial restoration after TR-Ab19. Dashed boxes indicate higher magnification. Labels: sc, synovial cavity; bm, bone marrow; st, synovial tissue; m, meniscus. (I) Flow-cytometry gating strategy for synovial cells. (J) Representative flow plots showing recovery of viable synovial CD45^+^ cells and enrichment of CD11b^+^F4/80^+^ macrophages, including CX_3_CR1^+^ subsets, after TR-Ab19. Related barrier-disruption controls and gating are in Fig. S13– S14. Data are shown as mean ± SEM unless otherwise stated; n values are indicated in the plots. Statistical tests follow Supplementary Methods; *P < 0.05, **P < 0.01, ***P < 0.001; n.s., not significant.

High-resolution confocal imaging of knee joints showed that healthy tdTomato mice harbor a continuous CX_3_CR1 lineage–traced macrophage lining (tdTomato^+^, red) along the synovial cavity, adjacent to bone marrow and meniscus (Fig. 4F). CIA disrupted this organization, producing fragmented, disordered tdTomato^+^ clusters (Fig. 4G). Notably, TR-Ab19 largely restored a continuous tdTomato^+^ layer along the synovial surface, with barrier-like architecture resembling healthy joints (Fig. 4H). Higher-magnification views of the boxed regions in Fig. 4F–H highlight the contrast between intact, broken, and rebuilt barriers.

To quantify these changes, we performed multicolor flow cytometry on synovial cells from CIA-tdTomato mice. A defined gating strategy (Fig. 4I) was used to identify live CD45^+^CD11b^+^ cells, F4/80^+^ macrophages, and CX_3_CR1^+^ subsets. Representative plots showed that TR-Ab19 increased the proportion of live synovial cells, expanded CD45^+^CD11b^+^ myeloid cells, and enriched F4/80^+^CX_3_CR1^+^ macrophages compared with untreated CIA-tdTomato joints (Fig. 4J), in line with the anatomic barrier restoration seen by imaging.

We next tested whether this effect requires TREM2. In TREM2 KO joints, F4/80 and CX_3_CR1 immunofluorescence showed a less continuous F4/80^+^CX_3_CR1^+^ lining at baseline, and CIA further disrupted CX_3_CR1^+^ macrophage organization. Importantly, TR-Ab19 did not restore a barrier-like layer in TREM2 KO CIA joints (Fig. S13), indicating that both barrier formation and its therapeutic reconstruction are TREM2-dependent. Complementary multicolor flow cytometry in conventional CIA cohorts (Healthy, CIA, CIA + TR-Ab19, and TREM2 KO CIA) using a parallel gating scheme (Fig. S14) showed that TR-Ab19 maintained F4/80^+^ and CX_3_CR1^+^ macrophages within live CD45^+^CD11b^+^ cells, whereas TREM2 KO CIA joints displayed an altered myeloid composition and expanded but disorganized CX_3_CR1^+^ cells.

### TR-Ab19 restrains germinal-center B cell responses and links TREM2 signaling to human RA

Having shown that TR-Ab19 rebuilds a TREM2-dependent CX_3_CR1^+^ synovial macrophage barrier, we next asked whether this local reprogramming is coupled to systemic B cell modulation. Using multicolor flow cytometry, we defined the key subsets of splenic CD19^+^ B cells, including naïve, germinal center (GC), plasmablast (Pb), memory, plasma (Pla/ASC) B cells, and T follicular helper (Tfh; CXCR5^+^PD-1^+^CD4^+^ T) cells (Fig. 5A). In CIA and TREM2 KO CIA mice, GC B cells, plasmablasts, Tfh cells, and histologic GC counts were all markedly increased relative to healthy controls, and spleens displayed enlarged, hyperplastic, and often disorganized GCs (Fig. 5B–F). TR-Ab19 substantially attenuated this GC axis, reducing plasmablast, GC B-cell, and Tfh frequencies and lowering GC numbers toward healthy levels. Across CIA and CIA + TR-Ab19 mice, the proportion of B cells among splenic lymphocytes inversely correlated with clinical arthritis scores (Fig. 5G), linking control of B-cell activation—and thus humoral immunity—to disease amelioration and to the barrier restoration described above.

**Fig. 5.**
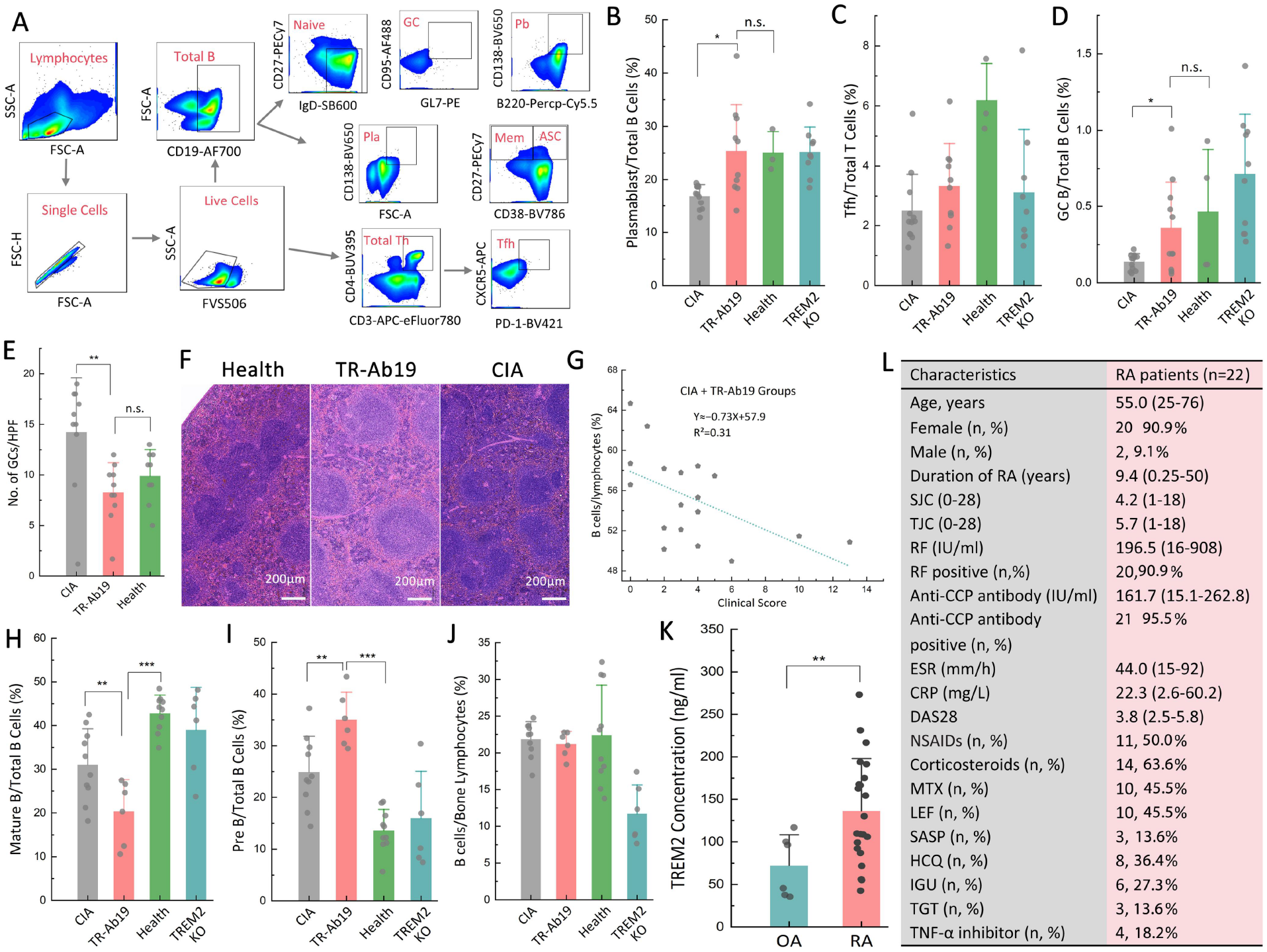
TR-Ab19 restrains germinal-center–associated B-cell activation and links TREM2 to human RA synovial-fluid sTREM2. (A) Flow-cytometry gating of splenic lymphocytes to define CD19^+^ B-cell subsets (naïve, GC, plasmablast, memory, plasma) and CD4^+^ Tfh cells (CXCR5^+^PD-1^+^). (B–D) Frequencies of plasmablasts, Tfh cells, and GC B cells across indicated mouse cohorts. (E–F) GC counts per high-power field and representative splenic H&E images (scale bar, 200 μm). (G) Spearman correlation between arthritis score and splenic B-cell proportion. (H–J) Bone marrow B-cell compartments (mature, pre-B, total; gating in fig. S15; extended analyses in fig. S16). (K) Synovial-fluid soluble TREM2 (sTREM2) in OA and RA patients. (L) RA cohort characteristics (n = 22). Data are mean ± SEM; n values are indicated in plots; statistics per Supplementary Methods (*P < 0.05, **P < 0.01, ***P < 0.001; n.s.). Abbreviations: GC, germinal center; Tfh, follicular helper T cell; OA, osteoarthritis; RA, rheumatoid arthritis.

We then examined whether TR-Ab19 also normalizes B-cell development in bone marrow. CD19^+^B220^+^ marrow cells were subdivided into pro-B, pre-B, immature, and mature B cells, with pre-BI, pre-BII, and pre-BIII stages further resolved (Fig. S15). CIA, particularly in TREM2 KO mice, disrupted this hierarchy, altering the balance of mature, pre-B, and total B cells (Fig. 5H–J). TR-Ab19 partially corrected these defects, decreasing mature B cells, increasing pre-B cells, and rebalancing total B-cell proportions toward healthy profiles (Fig. 5H–J). Extended analyses showed that CIA expanded pro-B and immature compartments and that TR-Ab19 partially normalized these shifts (Fig. S16A), while also moving splenic B-cell states toward a more physiological distribution with dampened GC/plasmablast responses (Fig. 5B,D and Fig. S16B). In contrast, TREM2 KO CIA mice retained or exacerbated these abnormalities, wherase the balance of TR-Ab in maintaining B-cell developmental and effector further reinforcing the TREM2 role in recovery arthritis.

To relate these mechanistic insights to human disease, we measured TREM2 level of synovial fluid in patients with osteoarthritis (OA) and RA. Soluble TREM2 concentrations in synovial fluid were higher in RA than OA (Fig. 5K), consistent with altered TREM2 biology in inflammatory joint disease; the cellular source and mechanistic interpretation (e.g., shedding versus increased expression/cell number) require dedicated validation. Clinical and laboratory characteristics of the RA cohort (n = 22), including demographics, serologic status, and inflammatory markers, are summarized in Fig. 5L and provide a translational backdrop for the mouse data.

Together with the structural preservation, cytokine modulation, and CX_3_CR1^+^ barrier reconstruction, these results show that TR-Ab19–mediated TREM2 activation coordinates innate and adaptive immunity: it rebuilds a synovial macrophage barrier, restrains splenic GC/Tfh–plasmablast responses, restores bone marrow B-cell homeostasis, and aligns with elevated free TREM2 level observed in synovial fluid of human RA.

### TR-Ab19 reprograms synovial macrophage states and TREM2 signaling to rebalance barrier and inflammatory programs

To connect TR-Ab19–mediated barrier reconstruction with underlying macrophage programs, we performed single-cell RNA sequencing of synovial macrophages collected from healthy, CIA, TR-Ab19, and TREM2-KO joints. Unsupervised clustering resolved four major macrophage states—CX_3_CR1^high^ “barrier-like”, Aqp1^+^ reparative, Clec4d^+^ inflammatory, and proliferating macrophages—visualized by UMAP and annotated using canonical markers and pathway signatures (Top2a/Mki67/Stmn1 for proliferating; Clec4d/Adam8/Arg2 for inflammatory; and complement/barrier/phagocytic programs for CX_3_CR1^high^ and Aqp1^+^ subsets) (Fig. 6A). Projection by experimental condition showed condition-dependent remodeling of the macrophage landscape (Fig. 6B, Figs. S17–S18). Composition analysis revealed that healthy joints were enriched for CX_3_CR1^high^ barrier-like macrophages, whereas CIA and TREM2-KO CIA joints were dominated by Clec4d^+^ inflammatory (and increased proliferative) states; notably, TR-Ab19 selectively expanded the CX_3_CR1^high^ compartment while contracting Clec4d^+^ and proliferating subsets, shifting macrophages toward barrier-like programs (Fig. 6C). Consistent with this, the Clec4d^+^ inflammatory fraction decreased with TR-Ab19 and remained elevated in disease contexts (Fig. 6D), whereas CX_3_CR1^+^ barrier-like macrophages increased with TR-Ab19 and were reduced in CIA/TREM2-KO CIA.

**Fig. 6.**
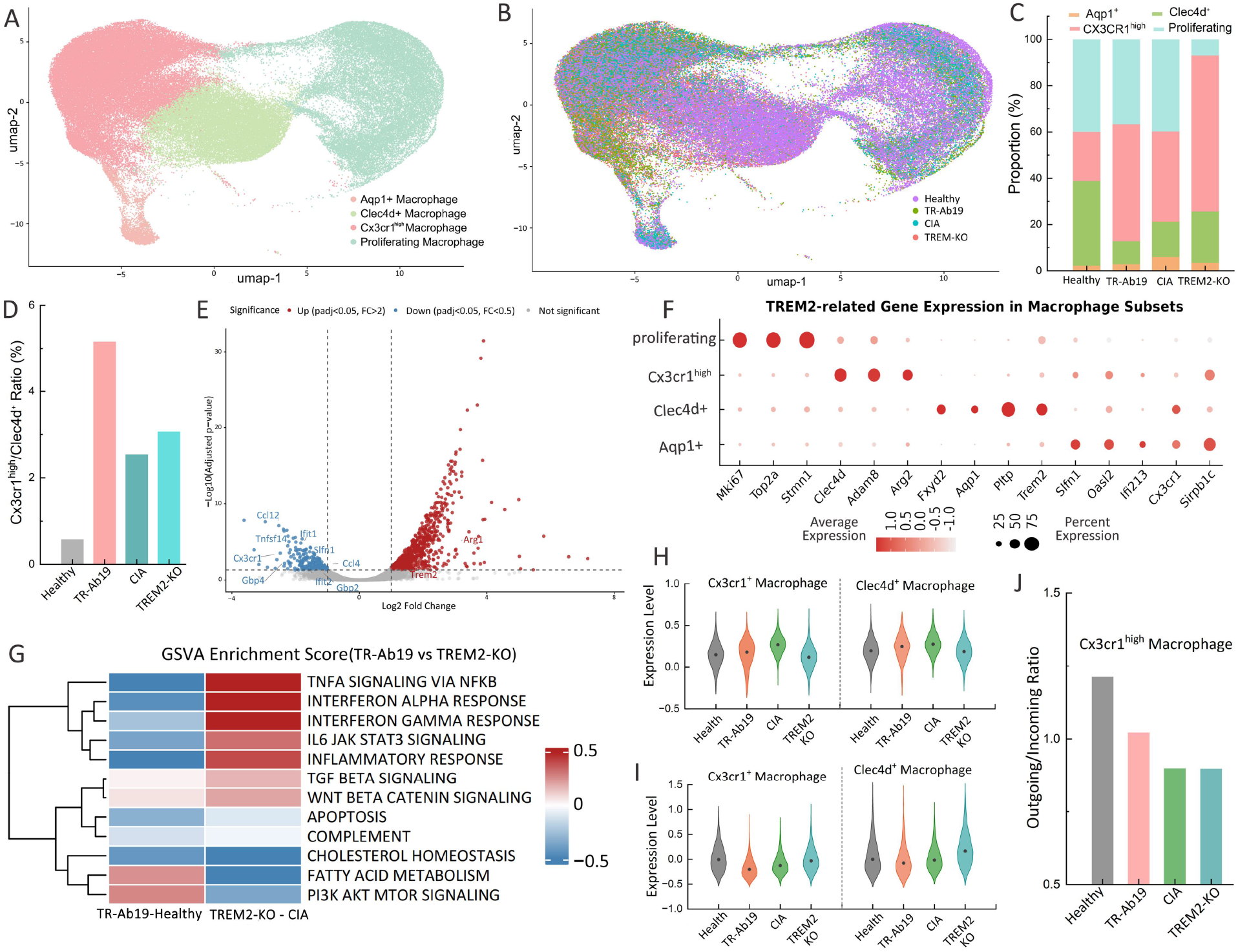
Single-cell transcriptomics reveals TR-Ab19–driven rebalancing toward barrier-like macrophage states. (A) UMAP of synovial macrophages annotated into four states: CX3CR1^high^ barrier-like, Aqp1^+^ reparative, Clec4d^+^ inflammatory, and proliferating macrophages (markers indicated). (B) UMAP split by condition (Healthy, CIA, CIA + TR-Ab19, and TREM2-KO CIA; Fig. S17–S18). (C–D) State composition and quantification of Clec4d^+^ inflammatory and CX3CR1^high^ barrier-like fractions across conditions. (E) Representative DEGs for indicated comparisons (multiple-testing correction in Supplementary Methods). (F) Feature plots of Trem2 and selected pathway genes across states. (G) GSVA pathway enrichment across conditions/states (Fig. S19–S22). (H–I) Tissue-repair and inflammation module scores computed per cell from curated gene sets (Fig. S23). (J) CellChat-derived outgoing/incoming signaling ratios indicating disease-associated network dominance and partial normalization with TR-Ab19 (Fig. S24). Data are mean ± SEM unless stated; n values are indicated in plots; statistics per Supplementary Methods (*P < 0.05, **P < 0.01, ***P < 0.001; n.s.). Abbreviations: DEG, differentially expressed gene; GSVA, gene set variation analysis; UMAP, uniform manifold approximation and projection.

Expression mapping of the TREM2 pathway showed that Trem2 and related genes were concentrated in the CX_3_CR1^high^ and Aqp1^+^ states but were reduced in Clec4d^+^ inflammatory macrophages and largely absent from the proliferating subset, indicating that inflammatory/cycling states lie outside the TREM2-governed barrier network (Fig. 6F). Differential expression analysis further supported global reprogramming with TR-Ab19, with reduced inflammatory signatures and a shift toward tissue-repair/homeostatic programs (representative differentially expressed genes shown) (Fig. 6E). At the pathway level, GSVA in synovial macrophages revealed enrichment of TNFα/NF-κB, IL-6–JAK–STAT3, interferon response, apoptosis, and complement programs in CIA and TREM2-KO CIA, whereas TR-Ab19 reverted these patterns toward a more metabolic/homeostatic profile (including lipid/cholesterol and fatty-acid metabolism–related signatures) (Fig. 6G). Across all subsets, GSVA highlighted PI3K-AKT and repairment signatures in Aqp1^+^ and CX_3_CR1^+^ cells treated using TR-Ab19, inflammatory cytokine pathways in Clec4d^+^ cells and proliferating macrophages (Figs. S19–S22). Module-score/violin-plot analyses corroborated that TR-Ab19 increased a tissue-repair module preferentially in CX_3_CR1^+^ macrophages (with comparatively weaker induction in Clec4d^+^ cells), while dampening an inflammation module—most prominently within the Clec4d^+^ inflammatory subset (Fig. 6H-I, Fig. S23). Finally, CellChat analysis of communication directionality showed that the outgoing/incoming signaling ratio was elevated in inflammatory networks (notably Clec4d^+^ macrophages) in disease settings and was partially normalized by TR-Ab19; corresponding ratios for Clec4d^+^ and CX_3_CR1^+^ subsets are shown (Fig. 6J, Fig. S24). Monocle pseudotime arranged macrophages along a trajectory from early barrier-like Aqp1^+^/CX_3_CR1^+^ states to late inflammatory/proliferative Clec4d^+^ and cycling states. Healthy samples occupied early pseudotime, whereas CIA and TREM2-KO were skewed toward late pseudotime, and TR-Ab19 shifted distributions back toward early, barrier-like positions (Fig. S25–S26), consistent with the restoration of lining-layer macrophages (Fig. 4F-H).

We next validated these shifts functionally. Multicolor flow cytometry of synovial barrier macrophages showed that CIA and TREM2-KO joints disorders the proportions of Clec4d^+^/proliferating and CX_3_CR1^high^/Aqp1^+^ cells, whereas TR-Ab19 recovered CX_3_CR1^+^/CX_3_CR1^high^ frequencies (Fig. 7A). In bulk synovium, qRT–PCR confirmed that TR-Ab19 upregulated Trem2 and co-receptors Sirpb1a/Sirpb1c, reshaped downstream mediators (Itpr3, Stat2, Socs3), and modulated Sirpb1b and Fosb compared with CIA and TREM2-KO joints (Fig. 7B-C; Fig. S27). Confocal imaging showed a dose-dependent increase in PI3K fluorescence in TR-Ab19–treated macrophages (0–10 μg ml^−1^), and immunoblotting demonstrated enhanced SYK activation in wild-type but not TREM2-KO cells (Fig. 7D–G), establishing a TREM2–SYK– PI3K axis directly engaged by TR-Ab19.

**Fig. 7.**
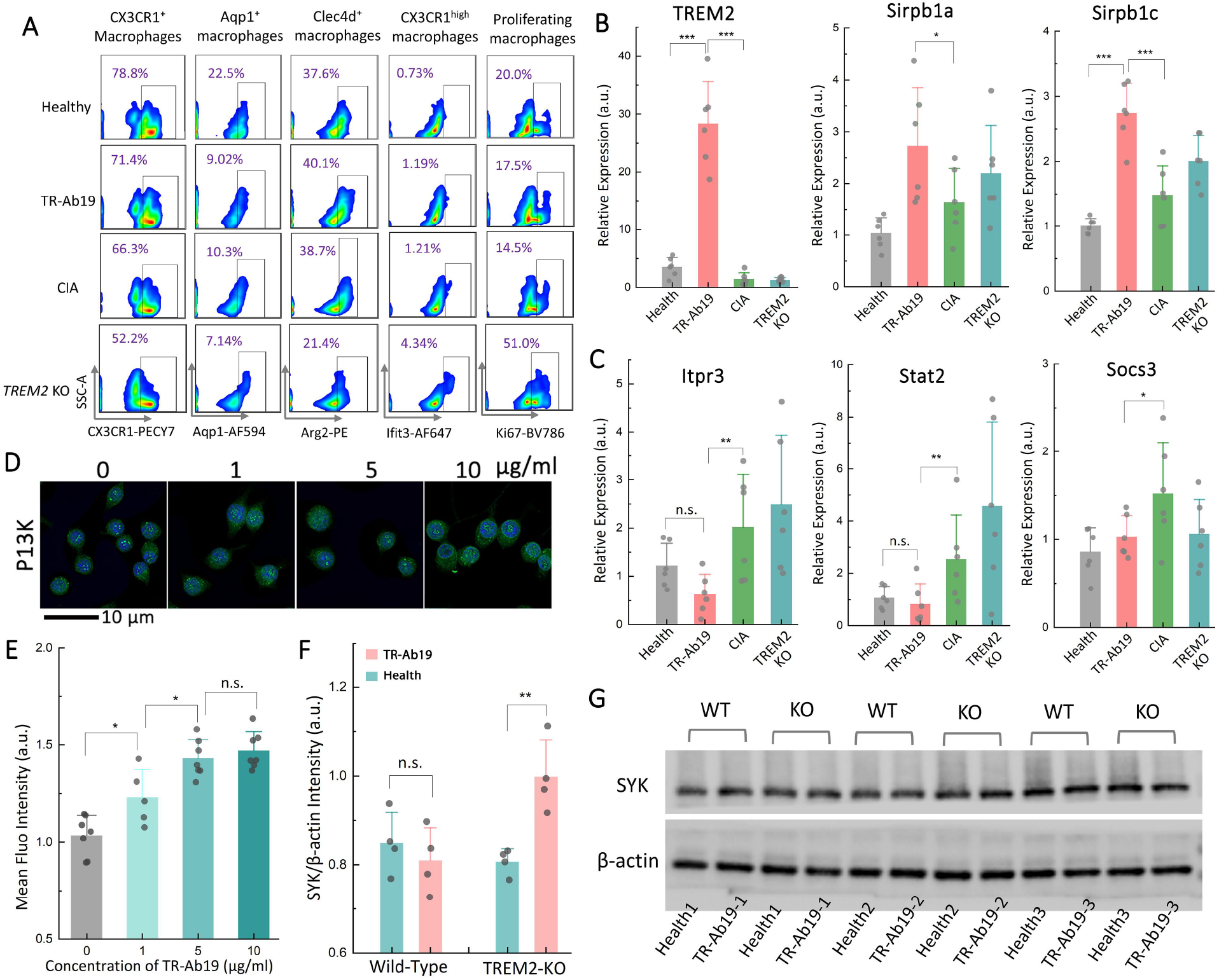
Validation of TR-Ab19–induced TREM2 signaling programs in synovial macrophages. (A) Flow cytometry of synovial F4/80^+^ macrophages from Healthy, CIA, CIA + TR-Ab19, and TREM2 KO CIA mice, quantifying CX_3_CR1^+^/CX_3_CR1^high^, Aqp1^+^, Clec4d^+^, and Ki67^+^ subsets (marker panel as indicated). (B–C) qRT–PCR of synovial TREM2-pathway genes: Trem2/Sirpb1a/Sirpb1c (B) and Itpr3/Stat2/Socs3 (C). (D–E) Macrophage stimulation with TR-Ab19 (0–10 μg ml^−1^) followed by PI3K immunofluorescence imaging (D; scale bar, 10 μm) and quantification (E). (F–G) SYK immunoblotting in WT and TREM2 KO macrophages treated with control IgG or TR-Ab19: representative blots (F) and densitometry (G). Data are mean ± SEM unless stated; n values are indicated in plots. Statistics per Supplementary Methods; *P < 0.05, **P < 0.01, ***P < 0.001; n.s., not significant. Abbreviations: CIA, collagen-induced arthritis; qRT– PCR, quantitative reverse transcription PCR; WT, wild type.

Together with the structural protection, cytokine modulation, barrier reconstruction, and B-cell rebalancing described above, these data show that TR-Ab19 reprograms synovial macrophages at single-cell and signaling levels—restoring a TREM2-dependent CX_3_CR1^+^/Aqp1^+^ barrier compartment, contracting Clec4d^+^ inflammatory and proliferating states, and normalizing their communication networks and trajectories.

## Discussion

Our study identifies TR-Ab19 as an on-target TREM2 agonist that enables barrier-centered myeloid state reprogramming in rheumatoid arthritis. From a binding-plus-function screen, TR-Ab19 binds mouse TREM2, robustly engages endogenous macrophage TREM2, and activates a myeloid reporter in a dose-dependent manner (Fig. 1A–M). Mechanistically, TR-Ab19 links TREM2 engagement to downstream signaling readouts and redirects macrophage states toward tissue-protective programs beyond blocking inflammatory ligand–receptor axes. Signaling and transcriptomic analyses support a coordinated shift toward protective states (Fig. 6A–G and Fig. 7A–E). At single-cell resolution, TR-Ab19 shifts synovial macrophage composition away from inflammatory/proliferative states and toward barrier-like and repair-associated phenotypes, supporting an instructive role of TREM2 agonism in reinforcing resident barrier identity (Fig. 6A–G).

This macrophage state shift is consistent with a tissue-scale sequence in which reinforcement of the STM lining barrier limits inflammatory influx, dampens destructive effector loops, and improves joint outcomes. Across arthritis models, TR-Ab19 improves disease and limits structural damage, with loss-of-function genetics supporting on-target dependence (Fig. 2A–J and Fig. 3A–H). Imaging and cytometry converge on a barrier-centered endpoint—reconstitution of continuous CX_3_CR1^high^ lining-layer organization accompanied by reduced inflammatory macrophage programs and leukocyte accumulation (Fig. 4A–J). Benefits extend beyond innate inflammation to adaptive remodeling, as TR-Ab19 is associated with reduced germinal-center activity and normalization of B-cell compartments (Fig. 5A–J). Together, these results support a feed-forward repair program initiated by TREM2 activation in resident STMs that limits influx-driven synovitis and downstream osteo-destructive circuitry (Fig. 2A–J; Fig. 3A–H; Fig. 4A–J).

Conceptually, this work advances a disease-modifying principle of “barrier repair through state control,” in which engaging an endogenous protective receptor stabilizes tissue-resident macrophage identity and restores the lining barrier as an explicit therapeutic endpoint (Fig. 4A–J). The convergence of *in vivo* efficacy, barrier reconstruction, and single-cell state shifts positions TREM2 agonism as a tractable strategy to reprogram inflammatory niches, analogous to microglial reprogramming in neurodegeneration, but deployed here to rebuild the joint lining barrier (Fig. 6A–G and Fig. 7A–E). Looking forward, patient-linked observations (Fig. 5K–L) motivate development of translational TREM2 agonists with optimized epitope/Fc geometry and delivery, alongside deeper dissection of barrier modules that couple receptor triggering to durable synovial homeostasis.

Our study aligns with a rapidly evolving RA landscape that is increasingly shaped by mechanism-anchored, cell-state–guided interventions. Single-cell stratification of synovium into inflammatory endotypes enables precision targeting of myeloid–stromal–lymphoid ecosystems [39], and emerging modalities—including lymphocyte “reset” strategies, programmable immune devices, and TAM-family– linked interventions—are redefining how immune circuits can be interrogated and therapeutically rewritten [40–46]. In parallel, multifunctional biomaterials that integrate immunomodulation with tissue repair further expand the interventional toolbox [47–51]. Against this backdrop, TR-Ab19 adds a complementary axis: reinstating a macrophage lining barrier to regulate inflammatory flux, thereby shifting the field from episodic “damage control” toward sustained barrier reconstruction as a disease-modifying strategy.

Several limitations and opportunities remain. Future studies should quantify whether TR-Ab19 affects TREM2 ectodomain shedding and define how epitope/Fc geometry couples agonism to barrier-state programming. Because TR-Ab19 is mouse-selective, translation will require human-reactive agonists with tuned valency/geometry and careful safety evaluation given broader TREM2 expression. Dynamic imaging and longitudinal single-cell profiling across diverse human pathotypes will be essential to define the timing, durability, and reversibility of barrier repair. Finally, combining barrier-rebuilding TREM2 agonists with DMARDs, biologics, or B-cell–directed therapies may enable dose reduction and more durable remission while preserving host defense. More broadly, analogous barrier-focused designs could be extended to other TREM2^+^/TAM^+^ macrophage niches (e.g., gut, lung, CNS), suggesting a general strategy for rebuilding myeloid barriers in chronic inflammatory and degenerative diseases.

## Materials and Methods

### Study design

This study aimed to (i) generate and characterize a mouse TREM2-agonistic monoclonal antibody (TR-Ab19), (ii) evaluate its therapeutic efficacy and immunomodulatory effects in murine inflammatory arthritis models, and (iii) profile synovial macrophage states and their remodeling by TR-Ab19 using flow cytometry, qRT-PCR and single-cell RNA sequencing (scRNA-seq). Animals were allocated to groups with comparable age, sex and, when applicable, disease stage. Group sizes, numbers of independent experiments and statistical tests are reported in the figure legends and Statistical analysis section. Investigators were blinded to treatment during clinical scoring and histological evaluation when feasible.

### Mice and arthritis models

All animal procedures were approved by the Ethics Committee of Biology and Medicine, Tsinghua University (24-SXD2) and performed under specific-pathogen–free conditions. K/BxN mice were generated by crossing KRN TCR-transgenic C57BL/6 mice with NOD mice [52]; Trem2^−/−^ (TRME2 KO) and *Cx3cr1*^*Cre*^R26-tdTomato mice were obtained from The Jackson Laboratory and maintained on matched backgrounds. K/BxN serum-transfer arthritis (K/BxN STA; hereafter STA) was induced in C57BL/6J recipients by a single intraperitoneal injection of pooled arthritogenic serum collected from clinically arthritic K/BxN donors (150–200 μl per mouse), and mice were analyzed 7 days after serum transfer. Collagen-induced arthritis (CIA) was induced in C57BL/6J, TRME2 KO and *Cx3cr1*^*Cre*^R26-tdTomato mice by intradermal immunization with chicken type II collagen emulsified in adjuvant (day 0) followed by a booster immunization (day 21), and mice were monitored for up to 35 days (day 56 after primary immunization).

### Clinical assessment of arthritis and TR-Ab19 treatment

Arthritis severity was evaluated by body weight, incidence, paw swelling, and a clinical index ranging from 0 to 16 (cumulative score of 0–4 per paw: 0, no signs of inflammation; 4, maximal swelling and erythema), scored in a blinded manner. Paw thickness was measured using a Vernier caliper. In STA, TR-Ab19 was administered intra-articularly (0.1 mg/ml, 20 μl per knee) on day 1 after serum transfer. In CIA, TR-Ab19 was injected intra-articularly on day 18 (before booster immunization) and, where indicated, followed by a single intraperitoneal dose (5 mg/kg) 1 week after the booster.

### Human specimens and sTREM2 measurement

Human studies were approved by the Institutional Medical Ethics Review Board of Peking University People’s Hospital (2016PHB163-01). RA patients fulfilled 1987 ACR or 2010 ACR/EULAR criteria; OA patients met ACR criteria for knee OA. Clinical data (age, sex, disease duration, DAS28, ESR, CRP, RF, anti-CCP) were recorded. Synovial fluid was obtained by sterile knee aspiration, centrifuged to remove cells and debris, and supernatants were stored at −80 °C. Soluble TREM2 concentrations were measured using a commercial ELISA.

### Generation and purification of TR-Ab19

A mouse TREM2-derived peptide epitope was synthesized and conjugated to KLH for immunization of BALB/c mice. High-titer clones were selected by indirect ELISA, and splenocytes were fused with Sp2/0 myeloma cells to generate hybridomas. Supernatants were screened by ELISA, and positive clones were subcloned and sequenced to confirm heavy- and light-chain variable regions. Monoclonal antibodies were purified from culture supernatants by protein A/G chromatography, buffer-exchanged into PBS and, when required, subjected to endotoxin removal for in vivo administration.

### Biochemical and functional characterization of TR-Ab19

Binding of TR-Ab19 to recombinant mouse and human TREM2 ectodomains was quantified by indirect ELISA and surface plasmon resonance (SPR; Biacore). Single-cycle kinetics with a 1:1 Langmuir model were used to derive association/dissociation rate constants and K_D_ values. Epitope specificity was assessed by peptide competition and alanine-scanning/point-mutant ELISAs. Cell-surface binding and TREM2 agonism were evaluated in RAW264.7 macrophages line, allowing determination of EC_50_ values from dose–response curves.

### Histology, immunofluorescence and micro-CT of joints

Ankle and knee joints were fixed in 4% paraformaldehyde, decalcified in 10% EDTA, embedded in paraffin and sectioned. Sagittal sections were stained with H&E (synovitis, pannus), Safranin O–Fast Green (cartilage proteoglycan) and TRAP (osteoclasts). Histological scores for inflammation, cartilage damage and bone erosion, as well as TRAP^+^ multinucleated osteoclast counts, were obtained using established criteria. For immunofluorescence, sections underwent antigen retrieval, blocking and incubation with primary and fluorophore-conjugated secondary antibodies; nuclei were counterstained with DAPI and imaged by confocal microscopy. Micro-CT of ankle/knee joints (SkyScan) was used to reconstruct 3D bone and quantify trabecular and cortical parameters using standardized ROIs and thresholds.

### Flow cytometry of synovial, splenic and bone marrow cells

Single-cell suspensions from synovium, spleen and bone marrow were prepared by enzymatic or mechanical dissociation. After Fc blocking, cells were stained with multicolor antibody panels. Synovial macrophages were defined as live CD45^+^CD11b^+^Ly6G^-^F4/80^+^ cells and further analyzed for markers including CX_3_CR1, Aqp1, Arg2, Ifit3 and Ki67. Splenic and bone marrow B-cell subsets (naïve, GC, plasmablast/plasma, and memory fractions) and Tfh cells were identified using established gating strategies. Data were acquired on a BD FACSDiscover S8/FACSymphony A5 and analyzed with FlowJo.

### Single-cell RNA sequencing and computational analysis

Synovial single-cell suspensions were processed on a BD single-cell platform and sequenced on an Illumina system. FASTQ files were converted to UMI count matrices aligned to mm10 using a platform-compatible pipeline (details and versions in Supplementary Methods), followed by standard QC and downstream analysis in Seurat/Harmony. After quality control (gene counts, mitochondrial fraction, doublet removal), data were normalized and integrated in Seurat with Harmony batch correction. UMAP was used for visualization and SNN-based clustering. Macrophage subsets were annotated using CX_3_CR1, Aqp1, Clec4d, Arg2, Ifit3, Mki67 and published references [11]. Differential expression, GSVA pathway analysis, CellChat ligand–receptor inference and Monocle-based trajectory analysis were used to define transcriptional programs, intercellular communication and pseudotime dynamics.

### Serological assays and statistical analysis

Serum cytokines were measured using a 17-plex bead-based immunoassay. Data are presented as mean ± s.e.m. unless stated otherwise. Normality was assessed before choosing parametric (t test, ANOVA) or non-parametric (Mann–Whitney, Kruskal–Wallis) tests, with appropriate multiple-comparisons corrections. For scRNA-seq differential expression, p values were adjusted using Benjamini–Hochberg FDR. P < 0.05 was considered statistically significant.

### Detailed materials and methods are provided in the Supplementary Information

## Supporting information

Supporting Infromation

## Acknowledgements

The authors thank the National Natural Science Foundation of China (No. 52272278, 92159305, 52402347) and National Postdoctoral Program for Innovative Talents (No. BX20230190) for their support of funding. We thank the Tsinghua University Flow Cytometry Platform and Xiuyu Zhou, as well as Xiaoxue Xu of the Capital Medical University Flow Cytometry Facility and Jianfeng Lei of the Animal CT Platform for their technical support. Thanks prof. Christophe Benoist and his lab from Harvard Medical School for sharing KRN model mice.

## Author Contributions

H.H.Y., L.J., and X.D.S. conceived the study. H.H.Y. and L.J. designed and generated the TR-Ab19 antibody. H.H.Y. and J.Y.W. performed most of the in vivo experiments, with assistance from L.X.L. Q.C.W. led the single-cell RNA-seq analyses, and W.C.S. contributed to early single-cell data analysis and interpretation. P.X.L. assisted with data analysis. Z.Y.L. and X.M.W. provided key experimental infrastructure, contributed to project discussions, and offered critical suggestions. F.L.H. and X.C. provided human samples and related clinical support. H.Y. joined the project mid-course, coordinated data integration and interpretation, provided experimental support for single-cell sequencing, and drafted the initial version of the manuscript. H.H.Y., J.Y.W., L.J., H.Y., and X.D.S. together with other co-authors, revised and approved the final manuscript.

## Data and materials availability

The data that support the figures within this paper are available upon reasonable request.

**Scheme 1.**
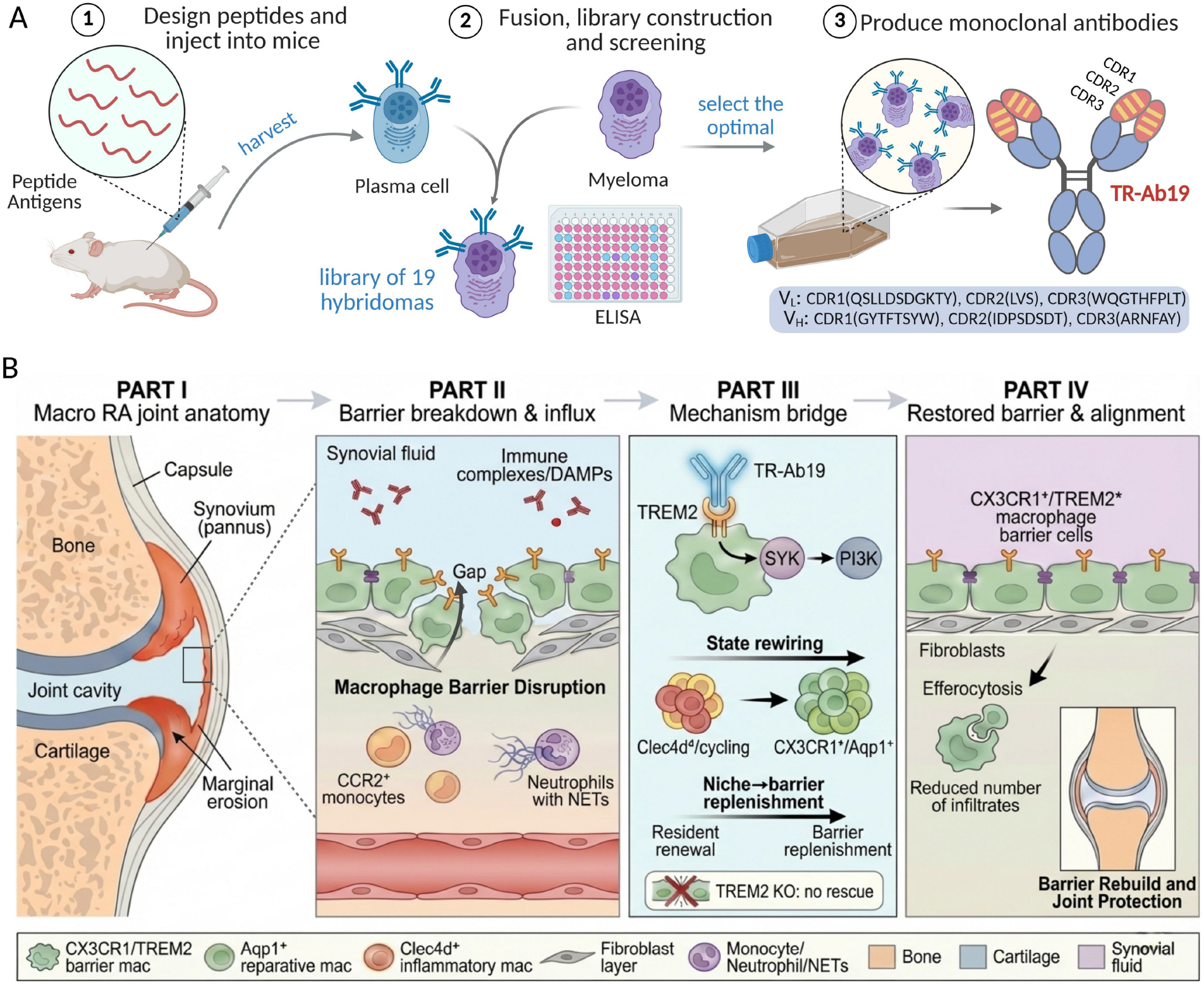
TR-Ab19 reprograms synovial macrophages to rebuild a TREM2^+^ lining barrier and protect arthritic joints. (A) Antibody-generation workflow: mice are immunized with TREM2 peptide epitopes, hybridomas are produced and screened by ELISA and functional assays, and the lead clone TR-Ab19 is selected (CDR sequences shown). (B) Working model (Parts I–IV): (I) Joint anatomy and synovial lining. (II) In active arthritis, the CX_3_CR1^+^/TREM2^+^ lining macrophage barrier is disrupted, permitting influx/accumulation of inflammatory cues and infiltrating myeloid cells. (III) TR-Ab19 engages TREM2 and promotes downstream signaling to shift macrophage states from Clec4d^+^ inflammatory/cycling programs toward CX_3_CR1^high^/Aqp1^+^ reparative/barrier-like states, enabling barrier replenishment. (IV) Barrier restoration is associated with improved lining organization, reduced inflammatory infiltrates, and protection of joint structure across CIA and STA models. Abbreviations: CIA, collagen-induced arthritis; STA, serum-transfer arthritis; STMs, synovial tissue macrophages; NETs, neutrophil extracellular traps.

