## Supporting Infromation for "TREM2 agonist antibody rebuilds the resident synovial macrophage lining barrier in rheumatoid arthritis"

### Materials and Methods

### KEY RESOURCES TABLE

| REAGENT or RESOURCE | SOURCE | IDENTIFIER |
|---|---|---|
| **Antibodies** | | |
| Goat anti-Mouse TREM2 Antibody | Abcam | Cat#ab95470 |
| Goat anti-Mouse IgG (H+L) Cross-Adsorbed Secondary Antibody, Alexa Fluor 488 | Thermo Fisher Scientific | Cat#A-11001 |
| Syk (D3Z1E) XP Rabbit mAb | Cell Signaling Technology | Cat#13198 |
| Anti-PI 3 Kinase p85 alpha Antibody | Abcam | Cat#ab191606 |
| Goat anti-Rabbit IgG (H+L) Cross-Adsorbed Secondary Antibody, Alexa Fluor 488 | Thermo Fisher Scientific | Cat#A-11008 |
| Goat anti-Mouse IgG (H+L) Secondary Antibody, HRP | Thermo Fisher Scientific | Cat#31430 |
| Goat anti-Rabbit IgG (H+L) Secondary Antibody, HRP | Thermo Fisher Scientific | Cat#31460 |
| Anti-beta Actin Antibody | Abcam | Cat#ab8226 |
| Brilliant Violet 421 anti-Mouse CD45 (30-F11) | BioLegend | Cat#103134; 1 µL |
| Anti-human/mouse CD11b PE-Cyanine 7 (M1/70) | BioLegend | Cat#101215; 1 µL |
| FITC anti-Mouse F4/80 (BM8) | BioLegend | Cat#123108; 2 µL |
| Alexa Fluor 647 anti-Mouse Ly-6G (1A8) | BioLegend | Cat#127609; 0.5 µL |
| PE anti-Mouse $CX_3CR1$ (SA011F11) | BioLegend | Cat#149005; 1 µL |
| Purified Rat anti-Mouse CD16/CD32 (2.4G2) | BD Biosciences (BD Pharmingen) | Cat#553142; 1:500 |
| Brilliant Violet 605 anti-mouse/human CD11b Antibody (M1/70) | BioLegend | Cat#101237; 1 µL |
| RB744 rat anti-Mouse F4/80 (T45-2342) | BD Biosciences (BD Horizon) | Cat#570508; 1 µL |
| FITC anti-Mouse Ly-6G Antibody (1A8) | BioLegend | Cat#127605; 1 µL |
| PE/Cyanine7 anti-Mouse $CX_3CR1$ Antibody (SA011F11) | BioLegend | Cat#149015; 1 µL |
| Aquaporin 1/AQP1 Antibody Alexa Fluor 594 (1/22) | Santa Cruz Biotechnology | Cat#sc-32737; 1 µL |
| Arginase 2/ARG2 Antibody PE (A-10) | Santa Cruz Biotechnology | Cat#sc-393496; 1 µL |
| IFIT3 Antibody Alexa Fluor 647 (E-10) | Santa Cruz Biotechnology | Cat#sc-393396; 1 µL |
| Ki-67 Monoclonal Antibody (SolA15), Brilliant Violet 786 | Thermo Fisher Scientific | Cat#417-5698-80; 2 µL |
| Goat Anti-Mouse IgA alpha Chain (HRP) | Abcam | Cat#ab97235 |
| Goat Anti-Mouse IgM mu Chain (HRP) | Abcam | Cat#ab97230 |
| Goat Anti-Mouse IgG1 (HRP) | Abcam | Cat#ab97240 |
| Goat Anti-Mouse IgG2a Heavy Chain (HRP) | Abcam | Cat#ab97245 |
| Goat Anti-Mouse IgG2b Heavy Chain (HRP) | Abcam | Cat#ab97250 |
| Goat Anti-Mouse IgG3 Heavy Chain (HRP) | Abcam | Cat#ab97260 |

| | | |
|---|---|---|
| CD45R (B220) Monoclonal Antibody (RA3-6B2), eFluor™ 450 | Thermo Fisher Scientific | Cat#48-0452-82; 2 µL |
| CD19 Monoclonal Antibody (eBio1D3), Super Bright 645 | Thermo Fisher Scientific | Cat#64-0193-82; 2 µL |
| IgM Monoclonal Antibody (II/41), PerCP-eFluor 710 | Thermo Fisher Scientific | Cat#46-5790-82; 2 µL |
| CD93 (AA4.1) Monoclonal Antibody (AA4.1), PE | Thermo Fisher Scientific | Cat#12-5892-82; 1 µL |
| CD25 Monoclonal Antibody (PC61.5), APC | Thermo Fisher Scientific | Cat#17-0251-82; 0.5 µL |
| CD43 Monoclonal Antibody (eBioR2/60), FITC | Thermo Fisher Scientific | Cat#11-0431-82; 0.25 µL |
| CD19 Monoclonal Antibody (eBio1D3), Alexa Fluor™ 700 | Thermo Fisher Scientific | Cat#56-0193-82; 2 µL |
| IgD Monoclonal Antibody (11-26c (11-26)), Super Bright™ 600 | Thermo Fisher Scientific | Cat#63-5993-82; 2 µL |
| CD27 Monoclonal Antibody (LG.7F9), PE-Cyanine7 | Thermo Fisher Scientific | Cat#25-0271-82; 2 µL |
| BV650 Rat Anti-Mouse CD138 | BD Biosciences (BD Horizon) | Cat#564068; 1 µL |
| CD95 (APO-1/Fas) Monoclonal Antibody (15A7), Alexa Fluor™ 488 | Thermo Fisher Scientific | Cat#53-0951-82; 4 µL |
| GL7 Monoclonal Antibody (GL-7 (GL7)), PE | Thermo Fisher Scientific | Cat#12-5902-82; 1 µL |
| PerCP/Cyanine5.5 anti-Mouse/Human CD45R/B220 Antibody (RA3-6B2) | BioLegend | Cat#103236; 1 µL |
| BV786 Rat Anti-Mouse CD38 (90/CD38) | BD Biosciences (OptiBuild) | Cat#740887; 1 µL |
| Brilliant Violet 421™ anti-Mouse CD279 (PD-1) Antibody (29F.1A12) | BioLegend | Cat#135217; 1 µL |
| CD185 (CXCR5) Monoclonal Antibody (SPRCL5), APC | Thermo Fisher Scientific | Cat#17-7185-82; 2 µL |
| BUV395 Rat Anti-Mouse CD4 | BD Biosciences (BD Horizon) | Cat#568375; 1 µL |
| CD3 Monoclonal Antibody (17A2), APC-eFluor™ 780 | Thermo Fisher Scientific | Cat#47-0032-82; 0.5 µL |
| CD117 (c-Kit) Monoclonal Antibody (2B8), APC-eFluor™ 780 | Thermo Fisher Scientific | Cat#47-1171-82; 1 µL |
| **Chemicals, Peptides, and Recombinant Proteins** | | |
| Recombinant Human TREM-2 Protein (ECD, His Tag) | Sino Biological | Cat#11084-H08H |
| Recombinant Mouse TREM-2 Protein (hFc Tag) | Sino Biological | Cat#50149-M02H |
| Chick Type II Collagen | Chondrex, Inc. | Cat#20012; 100 µg/ml |
| Complete Freund’s Adjuvant (5mg/ml M. Tuberculosis) | Chondrex, Inc. | Cat#7023 |
| Incomplete Freund’s Adjuvant | Chondrex, Inc. | Cat#7002 |
| Tamoxifen | Sigma-Aldrich (Merck) | Cat#T5648 |
| Corn oil | Sigma-Aldrich (Merck) | Cat#C8267 |
| Ghost dye red 780 viability dye | TONBO Biosciences | Cat#13-0865; 1:500 |

| | | |
|---|---|---|
| Brilliant Stain Buffer | BD Biosciences (BD Horizon) | Cat#563794 |
| Chloroform | Merck | CAS: 67-66-3 |
| Isopropyl alcohol | Merck | CAS: 67-63-0 |
| Fixable Viability Dye eFluor 506 | Thermo Fisher Scientific | Cat#65-0866-18; 1:500 |
| **Critical Commercial Assays** | | |
| Human TREM2 ELISA Kit | Abcam | Cat#ab224881 |
| Mouse TREM2 (Extracellular) SimpleStep ELISA | Abcam | Cat#ab309115 |
| ProcartaPlex Mo Th1/Th2/Th9/ Th17/Th22/Treg 17plex | Thermo Fisher Scientific | Cat#EPX170-26087-901 |
| EasySep mouse F4/80 positive selection kit | STEMCELL Technologies | Cat#100-0617 |
| Zombie Aqua fixable viability kit (FVS506) | BioLegend | Cat#423101; 1:500 |
| Transcription Factor Buffer Set | BD Biosciences (BD Pharmingen) | Cat#562574 |
| 5X All-In-One RT MasterMix | ABM (Applied Biological Materials) | Cat#G490 |
| Blastaq 2xPCR MasterMix | ABM (Applied Biological Materials) | Cat#G891 |
| **Experimental Models: Cell Lines** | | |
| RAW264.7 | ATCC | Cat#TIB-71 |
| **Experimental Models: Organisms/Strains** | | |
| C57BL/6J mice | Laboratory Animals Resources Center, Tsinghua University. | N/A |
| Non-obese diabetes (NOD) mice | Laboratory Animals Resources Center, Tsinghua University. | N/A |
| KRN, Tcra$^{-/-}$ mice | Gift from Harvard University | N/A |
| K/BxN STA mice | Generated in-house | N/A |
| C57BL/6J-*Trem2*$^{em2Adiuj}$/J | The Jackson Laboratory | Strain #:027197 |
| B6;129S6-*Gt(ROSA)26Sor*$^{tm9(CAG\text{-}tdTomato)Hze}$/J (*tdTomato*) | The Jackson Laboratory | Strain #:007905 |
| B6.129P2(C)-Cx3cr1$^{tm2.1(cre/ERT2)Jung}$/J (*Cx3cr1*$^{ERcre}$) | The Jackson Laboratory | Strain #:020940 |
| **Software and Algorithms** | | |
| DataViewer (version 1.5.6.2) | Bruker | SKYSCAN 1276; v1.5.6.2 |
| CT Analyser (version 1.17.7.2) | Bruker | SKYSCAN 1276; v1.17.7.2 |
| Realistic 3D-Visualization (version 2.3.2.0) | Bruker | SKYSCAN 1276; |

| | | |
|---|---|---|
| | | v2.3.2.0 |
| Rstudio | Posit | RStudio (version N/A) |
| Seurat (v5.2.1) | R package | v5.2.1 |
| DoubletFinder (v2.0.4) | R package | v2.0.4 |
| Harmony (v1.2.3) | R package | v1.2.3 |
| DESeq2 (v1.44.0) | Bioconductor | v1.44.0 |
| clusterProfiler (v4.12.6) | Bioconductor | v4.12.6 |
| STRING database | STRING | https://string-db.org/ |
| Cytoscape software (v3.10.3) | Cytoscape Consortium | v3.10.3 |
| ggplot2 (v3.5.1) | R package | v3.5.1 |
| pheatmap (v1.0.12) | R package | v1.0.12 |
| ggrepel (v0.9.6) | R package | v0.9.6 |
| ggVennDiagram (v1.5.2) | R package | v1.5.2 |
| SPSS | IBM | IBM SPSS Statistics (version N/A) |
| Graphpad | GraphPad Software | GraphPad Prism (version N/A) |
| Origin | Origin Software | Origin 2024 |

**Experimental Model and Subject Details**

**Cell lines**

Sp2/0 murine myeloma cells were maintained in Dulbecco's Modified Eagle Medium (DMEM; Sigma-Aldrich, Cat. No. D0822) supplemented with 10% fetal bovine serum (FBS; Gibco), 1% penicillin-streptomycin (Gibco), and 2 mM L-glutamine (Gibco) at 37°C in a humidified incubator with 5% $CO_2$. RAW264.7 macrophages (ATCC) were cultured in DMEM containing 10% FBS and 1% penicillin-streptomycin under the same conditions. Cell lines were routinely tested and confirmed negative for mycoplasma contamination using a PCR-based assay.

**Animals**

All animal procedures were approved by the Ethics Committee of Biology and Medicine, Tsinghua University (24-SXD2) and performed under specific-pathogen–free conditions. Trem2$^{-/-}$ (TREM2 KO) mice (C57BL/6J-*Trem2*$^{em2Adiuj}$/J; The Jackson Laboratory, stock 027197) and Cx3cr1$^{ERCre}$ mice (stock 020940) were obtained from The Jackson Laboratory. For lineage tracing, Cx3cr1$^{ERCre}$ mice were crossed with B6;129S6-*Gt(ROSA)26Sor*$^{tm9(CAG\text{-}tdTomato)Hze}$/J (Ai9; stock 007905) to generate Cx3cr1$^{ERCre}$;Rosa26LSL-tdTomato mice (hereafter *CX$_3$CR1*$^{cre}$R26-tdTomato, or CIA-tdTomato). Detailed experimental procedures are provided below (In Vivo Arthritis Models and Interventions).

**Human specimens**

Human specimen research was approved by the Institutional Medical Ethics Review Board of Peking University People's Hospital (approval number: 2016PHB163-01). Written informed consent was obtained from all participants. Rheumatoid arthritis (RA; n = 22) patients were enrolled by the

Department of Rheumatology, Peking University People's Hospital. RA patients fulfilled the 1987 American College of Rheumatology (ACR) revised criteria or the 2010 ACR/EULAR classification criteria, and OA patients met ACR criteria for knee osteoarthritis. Demographic and clinical information including age, sex, disease duration, swollen joint count (SJC), tender joint count (TJC), DAS28, erythrocyte sedimentation rate (ESR), C-reactive protein (CRP), rheumatoid factor (RF), and anti–cyclic citrullinated peptide antibody (anti-CCP) were recorded. Synovial fluid was obtained by sterile knee aspiration, centrifuged at 1500 rpm for 10 min to remove cells and debris, and supernatants were stored at −80°C until analysis.

**Generation and Characterization of TR-Ab19**

**Peptide antigen design and mouse immunization**

A mouse TREM2–derived peptide epitope (CDAGDLWVPEESSSFEGAQVEHSTSRNQET) was synthesized and conjugated to keyhole limpet hemocyanin (KLH). The peptide-KLH conjugate was diluted in 1× PBS to 1.2 mg/mL and emulsified 1:1 (v/v) with adjuvant immediately before immunization. Female BALB/c mice (6–8 weeks old; n = 4) received a primary subcutaneous immunization containing 60 µg peptide in complete Freund's adjuvant (CFA). Booster immunizations were administered on days 14, 28, and 42 using 30 µg peptide emulsified in incomplete Freund's adjuvant (IFA). Blood was collected from the inner canthus for serum isolation, and antibody titers were assessed by indirect ELISA using serial serum dilutions starting at 1:200. For terminal boosting, mice received an intraperitoneal injection of 50 µg peptide on day 56. Splenocytes from the mouse with the highest serum titer were harvested on day 59 for hybridoma generation.

**Hybridoma generation and screening**

Hybridomas were generated following the Köhler–Milstein fusion principle with minor modifications. Sp2/0 myeloma cells were expanded for 3 days before fusion. Splenocytes were combined with Sp2/0 cells at a myeloma:splenocyte ratio of 1:5 in DMEM (final volume, 1 mL). Fusion was initiated by gradual addition of 1 mL pre-warmed 50% polyethylene glycol (PEG 1500; Roche, Cat. No. 10783641001) at 37°C with gentle agitation. After 1 min incubation at room temperature, the mixture was slowly diluted in 10 mL DMEM to terminate fusion, and cells were collected by centrifugation. Fused cells were resuspended in HAT selection medium (Sigma-Aldrich, Cat. No. H0262) and plated into microplates. After 5 days of selection, medium was replaced with DMEM supplemented with HT (Sigma-Aldrich, Cat. No. H0137), and cultures were maintained for an additional 10 days to allow clone outgrowth.

Hybridoma supernatants were screened for anti-peptide reactivity by indirect ELISA. High-binding microplates were coated with 2 µg/mL BSA-coupled peptide, washed with PBS containing 0.05% Tween-20 (PBST), and blocked with 2% FBS for 1 h at room temperature. Hybridoma supernatants (100 µL per well) were incubated for 1 h; immunized mouse serum (1:1000) served as a positive control. After washing, HRP-conjugated goat anti-mouse IgG secondary antibody (1:5000) was added for 1 h. Plates were developed using TMB substrate at 37°C for 10 min, the reaction was stopped with 2 N $H_2SO_4$, and absorbance was measured at 450 nm. Clones with robust, specific signals were expanded for confirmatory assays and downstream characterization.

### Variable region amplification and sequencing

To support clone tracking and sequence verification, variable regions of candidate hybridomas were amplified by PCR using standard murine immunoglobulin primer sets. PCR products were analyzed by agarose gel electrophoresis and, where needed, purified and submitted for Sanger sequencing. Sequences were used for internal clone identification and to confirm consistency across passages.

### Antibody purification

For downstream biochemical assays and in vivo studies, monoclonal antibodies were purified from hybridoma culture supernatants by protein A/G affinity chromatography and buffer exchanged into sterile PBS. When required for in vivo administration, purified antibodies were passed through endotoxin-removal columns and tested to ensure endotoxin levels were below commonly accepted thresholds for mouse dosing.

### Recombinant TREM2 proteins

Recombinant mouse TREM2 extracellular domain protein was obtained from Sino Biological (Cat. No. 50110-M08H) and used for ELISA and surface plasmon resonance (SPR) analyses. Where indicated, recombinant human TREM2 and/or engineered TREM2 mutants were used for cross-species binding assessment and epitope mapping (see ELISA and epitope mapping sections).

### ELISA-based binding assays and epitope mapping

Indirect ELISA was used to quantify TR-Ab19 binding to recombinant TREM2 proteins and to evaluate epitope specificity. High-binding 96-well plates were coated overnight at 4°C with recombinant mouse or human TREM2 extracellular domain (typically 1–2 µg/mL in carbonate coating buffer). Plates were washed with PBST and blocked with 2% FBS (or 3% BSA) for 1 h at room temperature. TR-Ab19, commercial anti-TREM2 antibodies, or isotype controls were added in serial dilutions and incubated for 1 h. After washing, HRP-conjugated secondary antibodies were applied for 1 h, developed with TMB, and read at 450 nm. Binding curves were fitted by nonlinear regression (four-parameter logistic) to estimate apparent $EC_{50}$ values when appropriate.

For peptide competition assays, TR-Ab19 was pre-incubated with synthetic peptide corresponding to the mapped mouse TREM2 epitope (amino acids 154–165) at the indicated concentrations before being added to TREM2-coated plates. Competitive inhibition was quantified as the reduction in ELISA signal relative to antibody alone. For alanine-scanning or point-mutation mapping, BSA-coupled mutant peptides were used as coating antigens in the indirect ELISA format, enabling residue-level assessment of TR-Ab19 recognition.

### Surface plasmon resonance (SPR)

Binding kinetics between TR-Ab19 and recombinant mouse TREM2 were measured by SPR using a Biacore T200 instrument (Cytiva). TR-Ab19 was captured on a protein A sensor chip to a target immobilization level of ~300 response units (RU), and recombinant mouse TREM2 was injected as a five-point concentration series (3.12–50 nM) in single-cycle kinetics mode at 25°C. Each injection consisted of a 180-s association phase followed by a 600-s dissociation phase. Sensorgrams were reference-subtracted and fitted to a 1:1 Langmuir binding model using Biacore evaluation software to estimate association (Ka), dissociation (k_d), and equilibrium dissociation constants ($K_D$).

### Macrophage binding and functional activation assays

For cell-surface binding assays, RAW264.7 macrophages were seeded on glass coverslips and allowed to adhere overnight. Cells were incubated with TR-Ab19, a commercial anti-TREM2 antibody, or isotype control at the indicated concentrations (e.g., 0.1, 1, and 10 μg/mL) at 4°C to minimize internalization. After washing with cold PBS, bound antibodies were detected using fluorophore-conjugated secondary antibodies, nuclei were counterstained with DAPI, and images were acquired using a confocal microscope. Fluorescence intensities were quantified in ImageJ using identical acquisition settings across conditions.

To quantify TREM2 signaling activity, an NFAT-luciferase reporter RAW264.7 macrophage cell line engineered to report TREM2/DAP12 activation was used. Reporter cells were treated with serial dilutions of TR-Ab19, commercial anti-TREM2 antibody, or isotype control for the indicated duration, and luciferase activity was measured using a luminescence-based assay. Dose–response curves were fitted by nonlinear regression to determine $EC_{50}$ values.

### NFAT–luciferase reporter assay for TREM2 signaling activation

To quantify agonist-driven activation of canonical TREM2 signaling, we used RAW264.7 macrophages harboring a stable NFAT–luciferase reporter. Cells were maintained in Minimum Essential Medium supplemented with 10% fetal bovine serum, 100 U/mL penicillin, 100 μg/mL streptomycin, and 0.25 μg/mL amphotericin B at 37°C in 5% $CO_2$, consistent with our general RAW264.7 culture conditions. Cells were seeded into white, opaque 96-well plates and allowed to adhere overnight. The next day, cells were treated with serial dilutions of TR-Ab19 or isotype-matched control IgG (added as soluble antibodies directly to the culture medium, without plate pre-coating) for 24 h.

After stimulation, NFAT reporter activity was quantified by measuring luciferase luminescence using a commercial luciferase detection reagent according to the manufacturer's instructions. Luminescence was recorded on a microplate luminometer and reported as relative light units (RLU). Background signal (medium/reagent only) was subtracted, and values were plotted as dose–response curves. Each titration was performed with n = 3 independent biological repeats.

$EC_{50}$ values were calculated by nonlinear regression in GraphPad Prism using a sigmoidal dose–response model (variable slope; four-parameter logistic), following the reporting convention used for antibody titration–based functional assays in *Science Translational Medicine* (including presentation as mean ± SD and $EC_{50}$ estimation from fitted curves).

### *In Vivo* Arthritis Models and Interventions

### Collagen-induced arthritis (CIA) model and TR-Ab19 treatment

CIA was induced in C57BL/6J mice using chicken type II collagen (CII; Chondrex, Cat. No. 20012) following standard procedures. Briefly, CII was dissolved at 2 mg/mL in 0.05 M acetic acid overnight at 4°C and emulsified 1:1 (v/v) with complete Freund's adjuvant (CFA; Chondrex, Cat. No. 7001). Mice received a primary intradermal injection of 100 μg CII at the base of the tail (day 0). On day 21, mice received a booster injection of 100 µg CII emulsified in incomplete Freund's adjuvant (IFA; Chondrex, Cat. No. 7002). TR-Ab19 dosing schedules, treatment groups, monitoring windows, and predefined endpoints are described in the main manuscript. For CIA induction, chicken type II collagen (CII; Chondrex, Cat. No. 20012) was dissolved at 2 mg/mL in 0.05 M acetic acid overnight at 4°C and emulsified 1:1 (v/v) with CFA (Chondrex, Cat. No. 7001). Mice received a primary intradermal injection of 100 µg CII at the base of the tail (day 0) and a booster of 100 µg CII emulsified in IFA (Chondrex, Cat. No. 7002) on day 21. Injection volume per mouse and emulsification quality were kept consistent across cohorts.

**Serum-transfer arthritis (STA) model and intra-articular antibody dosing**

Serum-transfer arthritis (STA) was induced in C57BL/6J mice by intraperitoneal injection of arthritogenic serum collected from arthritic K/BxN donor mice. STA induction, dosing schedules, and endpoints are described in the main manuscript. Arthritogenic serum was collected from clinically arthritic K/BxN donors, pooled from multiple mice, aliquoted and stored at −80°C; a single aliquot was thawed per experiment to minimize freeze–thaw cycles. Recipient C57BL/6J mice received intraperitoneal injection of pooled serum at the indicated volume. K/BxN mice were generated by crossing KRN TCR-transgenic C57BL/6 mice with NOD mice as previously described; arthritogenic serum was collected from clinically arthritic K/BxN donors.

**Clinical assessment of arthritis**

Arthritis incidence, clinical scoring, body weight and paw swelling were assessed as described in the main manuscript. Clinical scoring and histological quantification were performed blinded to treatment when feasible, and paw thickness was measured using a Vernier caliper at the indicated time points.

**Tamoxifen induction for $CX_3CR1$ lineage labeling**

To label $CX_3CR1$ lineage–positive cells, $CX_3CR1^{cre}$R26-tdTomatomice received tamoxifen (Sigma-Aldrich, Cat. No. T5648) prepared at 40 mg/mL in corn oil and administered intraperitoneally at 50 µL per mouse, twice within 72 h. For baseline lineage-labeling analyses, mice were euthanized 7 days after the final injection for tissue processing and imaging.

**Histology, Immunofluorescence, and Imaging**

**Joint tissue processing and histology**

Hind limbs were dissected and fixed in 4% paraformaldehyde (PFA) at 4°C for 48 h. Samples were decalcified in 10% (w/v) EDTA (pH 7.4) at 4°C for 2–3 weeks with frequent solution changes, processed, and embedded in paraffin. Serial sagittal sections (5 µm) were cut for hematoxylin and eosin (H&E) staining to assess synovial inflammation and pannus formation, Safranin O–Fast Green staining to evaluate cartilage integrity, and tartrate-resistant acid phosphatase (TRAP) staining to identify osteoclasts. Histopathological scoring was performed using established criteria for synovitis, cartilage degradation, and bone erosion, as described in the corresponding figure legends.

**Immunofluorescence staining of synovium and joint sections**

For immunofluorescence, paraffin sections were baked at 60°C for 2 h, deparaffinized in xylene, and rehydrated through graded ethanol. Sections were permeabilized with 0.5% Triton X-100 for 30 min and subjected to heat-mediated antigen retrieval in citrate buffer. After blocking with 5% BSA for 1 h at room temperature, sections were incubated with primary antibodies overnight at 4°C. After washing, sections were incubated with species-appropriate fluorophore-conjugated secondary antibodies for 1 h at room temperature, counterstained with DAPI, and mounted for imaging. Where indicated, TO-PRO-3 iodide (Invitrogen) was used for nuclear counterstaining in multichannel imaging.

**Microscopy and image analysis**

Imaging was performed on an Olympus SpinSR system or a Zeiss LSM900 confocal microscope (20× or 63× objectives), as specified for each experiment. Images were acquired using identical exposure/laser settings for all samples within a given experiment. Fluorescence quantification was performed in ImageJ (NIH) or vendor software (e.g., Zeiss ZEN or Leica LAS X) using consistent thresholds and regions of interest across groups.

**Micro–computed tomography (micro-CT)**

Ankle joints were scanned using a SkyScan 1172 micro-CT system (Bruker). Reconstructed images were analyzed in CTAn software (Bruker) to quantify bone structural parameters including cortical bone surface/total volume (Cr.BS/TV), cortial bone surface (Cr.BS), cortical bone volume (Cr.BV), cortical tissue volume (Cr.TV), trabecular bone volume (Tb.BV), trabecular tissue volume (Tb.TV), trabecular number (Tb.N), trabecular thickness (Tb.Th), and trabecular separation (Tb.Sp). Regions of interest and segmentation thresholds were applied consistently across groups within each experiment.

**Serological and Soluble Mediator Measurements**

**Multiplex cytokine analysis**

Serum cytokine and chemokine concentrations were quantified using a bead-based multiplex immunoassay (Bio-Plex Pro Mouse Cytokine 17-plex; Bio-Rad) according to the manufacturer’s instructions. Standard curves were generated for each analyte, and samples were analyzed on a compatible Luminex platform. Analytes included TNF-α, IL-1β, IL-6, IFN-γ, MCP-1, and other cytokines/chemokines as provided in the kit panel.

**TREM2 ELISA**

Soluble TREM2 (sTREM2) concentrations in human synovial fluid were measured using a commercial ELISA kit (Abcam, Cat. No. ab224881) following the manufacturer’s protocol. Samples were thawed on ice, diluted as needed to fall within the linear range of the assay, and assayed in duplicate.

**Single-Cell Preparation, Flow Cytometry, Cell Sorting, and qRT-PCR**

**Synovial tissue digestion and single-cell preparation**

Murine synovial tissues were dissected and minced into small fragments, then digested in serum-free RPMI 1640 containing collagenase IV (2 mg/mL; Sigma-Aldrich, Cat. No. C5138), Dispase I (5 mg/mL; Sigma-Aldrich, Cat. No. D4818), and DNase I (0.2 mg/mL; Roche, Cat. No. 10104159001) at 37°C with gentle agitation for 30 min. Digested suspensions were filtered through 70-µm cell strainers, washed with staining buffer (PBS with 2% FBS), and counted. For bone marrow and spleen, single-cell suspensions were prepared by mechanical dissociation and passage through 70-µm strainers; red blood cells were lysed using an ammonium chloride-based lysis buffer when required.

**Flow cytometry staining and acquisition**

Cells were blocked with anti-CD16/32 (Fc block) and stained with fluorophore-conjugated antibodies in staining buffer for 30 min at 4°C. Synovial macrophage phenotyping used combinations of markers including CD45, CD11b, F4/80, Ly6C, Ly6G, TREM2, $CX_3CR1$, Aqp1, Clec4d, Arg2, Ifit3, Ki67, and others as required for subset definition. Bone marrow B-cell development was analyzed using antibodies against CD19, B220, CD93, IgM, CD43, CD25, and CD117 as indicated for Hardy fraction gating. After staining, cells were washed and resuspended in staining buffer for acquisition. Data were acquired on a BD FACSymphony A5/FACSDiscover S8 flow cytometer (BD Biosciences) using BD FACSDiva software (v8.0.2) and analyzed in FlowJo v10.8.1 (BD Biosciences). Gating strategies are provided in the corresponding supplementary figures.

**Magnetic-Activated Cell Sorting （MACS）**

Single cell suspensions from synovium were generated as described above. The macrophages were enriched via MACS sorting using EasyStep mouse F4/80 positive selection kit (STEMCELL, 100-0671) diluted 1:10 in FACS buffer for further sorting following the manufacturer's instructions.

**Fluorescence-activated cell sorting (FACS)**

For downstream transcriptional assays, synovial macrophages were sorted by FACS as live $CD45^+CD11b^+Ly6G^-F4/80^+$ cells (and further subset gates where indicated). Sorted cells were collected into chilled collection buffer containing RNase inhibitors and immediately processed for single-cell capture as described below.

**Single-cell RNA Sequencing and Computational Analysis**

**Single-cell capture and library preparation**

Single-cell suspensions from synovial tissues were prepared as described above and adjusted to ~1000 cells/µL in ice-cold PBMC buffer. Where indicated, cells from different conditions were barcoded using the BD Mouse Single-Cell Multiplexing Kit (BD Biosciences, Cat. No. 633781), washed, and combined at equal ratios. Single-cell capture and library construction were performed on a BD single-cell platform following the manufacturer's protocol. Libraries were sequenced on an Illumina platform to a depth sufficient for downstream analyses.

**Preprocessing and quality control**

Raw sequencing reads were processed using DropSeqPipe to generate gene-by-cell UMI count matrices, with alignment to the mouse reference genome (mm10). Downstream analyses were performed in R using Seurat (v5.2.1). Cells were retained if they contained 200–3000 detected genes and <15% mitochondrial transcript fraction. Potential doublets were detected and removed using DoubletFinder. Additional filtering steps removed low-quality cells enriched for red blood cell and platelet transcripts and cells lacking macrophage lineage markers.

#### Normalization, integration, clustering, and annotation

Filtered count matrices were normalized and scaled in Seurat; highly variable genes were identified for dimensionality reduction. Principal component analysis (PCA) was performed on variable features, and batch effects across experiments and/or multiplexed samples were corrected using Harmony. Uniform Manifold Approximation and Projection (UMAP) was used for visualization. Graph-based clustering was performed using shared nearest-neighbor (SNN) modularity optimization, and clusters were annotated using canonical marker genes (e.g., Cx3cr1, Aqp1, Clec4d, Arg2, Ifit3, Mki67) together with published references.

#### Differential expression and pathway enrichment

Differential expression between groups or clusters was assessed using Seurat's FindMarkers/FindAllMarkers functions (Wilcoxon rank-sum test) with multiple-testing correction. For selected comparisons, pseudo-bulk differential expression analyses were performed using DESeq2. Functional enrichment analyses were conducted using clusterProfiler with Gene Ontology (GO) and Kyoto Encyclopedia of Genes and Genomes (KEGG) annotations.

#### Cell–cell communication and trajectory analysis

Ligand–receptor interaction networks were inferred using CellChat, with analysis restricted to expressed genes in the curated CellChat database and default filtering thresholds. Comparative communication strengths across conditions were visualized by aggregate network plots and signaling pathway-level summaries. Cell-state trajectories were reconstructed using Monocle 2. Macrophage subsets were embedded in reduced-dimensional space, ordered in pseudotime, and gene expression dynamics along trajectories were summarized using generalized additive models and/or Monocle's graph-learning framework.

### *In Vitro* Signaling Analyses

#### Western blotting

Cells were lysed in ice-cold RIPA buffer supplemented with protease and phosphatase inhibitors. Protein concentrations were determined using a BCA assay. Equal amounts of protein were resolved by SDS-PAGE and transferred to PVDF membranes. Membranes were blocked with 5% non-fat milk in TBST for 1 h at room temperature and incubated with primary antibodies overnight at 4°C. After washing with TBST, membranes were incubated with HRP-conjugated secondary antibodies for 1 h at room temperature. Signals were developed using enhanced chemiluminescence (ECL) reagents and imaged on a chemiluminescence detection system. Band intensities were quantified using ImageJ and normalized to loading controls.

#### Immunofluorescence staining of cultured cells

For signaling readouts by immunofluorescence, macrophages were cultured on glass coverslips, treated as indicated, fixed in 4% PFA, and permeabilized with Triton X-100. After blocking with 5% BSA, cells were incubated with primary antibodies against targets such as PI3K pathway components or macrophage markers, followed by fluorophore-conjugated secondary antibodies. Nuclei were counterstained with DAPI and samples were imaged by confocal microscopy. Quantification was performed using ImageJ with identical settings across conditions.

**Quantification and statistical analysis**

Statistical analyses were performed in GraphPad Prism (v10.2.0) and R (v4.3.1). Data are presented as mean ± SEM unless otherwise stated. Normality of distributions was assessed using the Kolmogorov–Smirnov and/or Shapiro–Wilk tests. For comparisons between two groups, two-tailed unpaired Student's t tests were used for normally distributed data; otherwise, the Mann–Whitney U test was applied. For comparisons among three or more groups, one-way ANOVA with Tukey's multiple-comparisons test (parametric) or Kruskal–Wallis with Dunn's multiple-comparisons test (nonparametric) was used. For experiments involving two independent variables (e.g., treatment × time), two-way ANOVA with appropriate multiple-comparisons correction (Sidak or Tukey) was used; repeated-measures ANOVA was used for longitudinal measures when applicable. For scRNA-seq differential expression and pathway analyses, multiple-testing correction was applied as described (e.g., Benjamini–Hochberg false discovery rate control). P values < 0.05 were considered statistically significant. Significance levels are denoted as *P < 0.05, **P < 0.01, and ***P < 0.001.

**Lymphoid Tissue Analysis and Follicle Imaging**

**Flow cytometry of spleen, lymph node, and bone marrow**

To evaluate systemic immune remodeling, spleen and draining lymph node cells were analyzed by multicolor flow cytometry. After Fc blocking, cells were stained with antibody panels to identify T cells (CD3, CD4), T follicular helper (Tfh) cells ($CD4^+CXCR5^+PD\text{-}1^+$), B cells ($CD19^+B220^+$), germinal center (GC) B cells ($B220^+GL7^+Fas^+$), plasmablasts/plasma cells ($B220^{lo/-}CD138^+$), and memory B-cell subsets ($CD38^+$ and/or IgD/IgM-based gates). For bone marrow B-cell development, Hardy fractions were defined from CD19/B220-gated cells using sequential CD93, IgM, CD43, CD25, and CD117 gates. Absolute cell counts were calculated from total viable cell numbers and subset frequencies.

**Sample collection and processing**

For terminal analyses, mice were deeply anesthetized and blood was collected by cardiac puncture. Serum was isolated by clotting at room temperature, centrifugation at 2000g for 10 min, and storage at −80°C until use. Joints were dissected for either histology (fixed immediately in 4% PFA) or single-cell preparation (kept in ice-cold RPMI with 2% FBS and processed promptly). Spleens and lymph nodes were weighed, photographed when appropriate, and processed into single-cell suspensions on the day of harvest. All samples within a given experiment were processed using identical protocols and in parallel to minimize batch effects.

Raw projections were reconstructed into axial image stacks using manufacturer-provided reconstruction software with beam-hardening correction. Analyses were performed by an investigator blinded to treatment group, and identical reconstruction and segmentation parameters were applied across samples within each cohort.

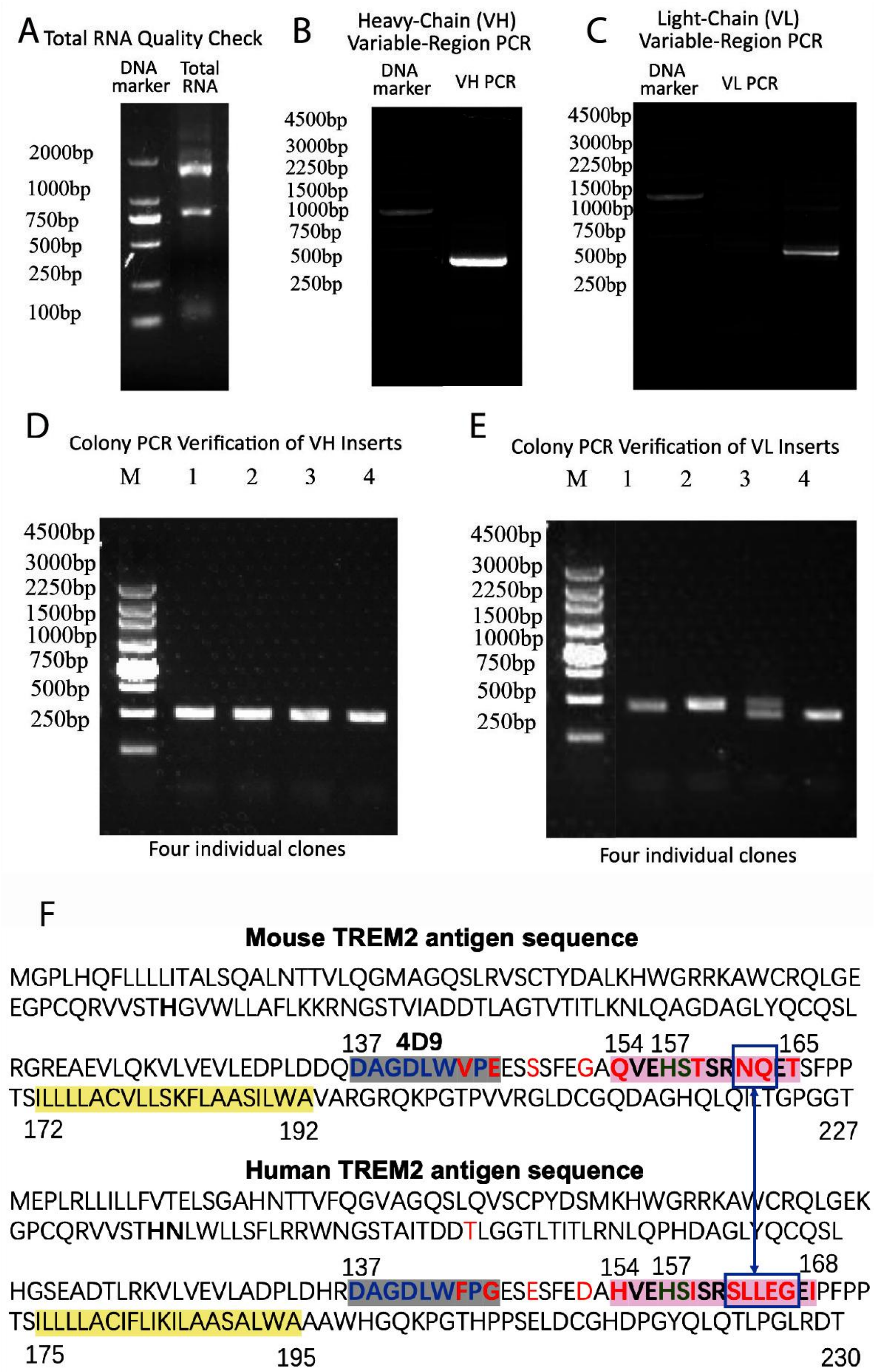


**Figure S1**. **Amplification and cloning of TR-Ab19 heavy- and light-chain variable regions and**

**sequence-level epitope inspection scheme of TR-Ab19**. A, Agarose gel quality check of total RNA isolated from the TR-Ab19–secreting hybridoma, showing intact ribosomal RNA bands; lane 1, DNA size marker; lane 2, total RNA. B–C, RT-PCR amplification of immunoglobulin variable regions from cDNA. B, Heavy-chain (VH) variable region PCR yields a single product of the expected size. c, Light-chain (VL) variable region PCR similarly produces a single specific band. D–E, Colony PCR verification of cloned VH (d) and VL (e) inserts in expression vectors. Lanes 1–4 represent four independent bacterial colonies; all positive colonies display bands corresponding to the expected VH or VL insert size. M, DNA size marker. F, Sequence-level epitope inspection suggested that the commercial anti-TREM2 antibody targets a segment encompassing the reported ADAM10/17 cleavage region (aa157-158, H and S), whereas the previously reported 4D9 epitope does not appear to cover this site. Notably, our TR-Ab19 epitope-mapping/structural prediction places its binding footprint across the 4D9-adjacent segment and into the cleavage-proximal region, raising the possibility that TR-Ab19 may reduce ectodomain shedding via steric hindrance while engaging TREM2. This proposed anti-shedding contribution remains a working hypothesis and was not directly tested in the current study.

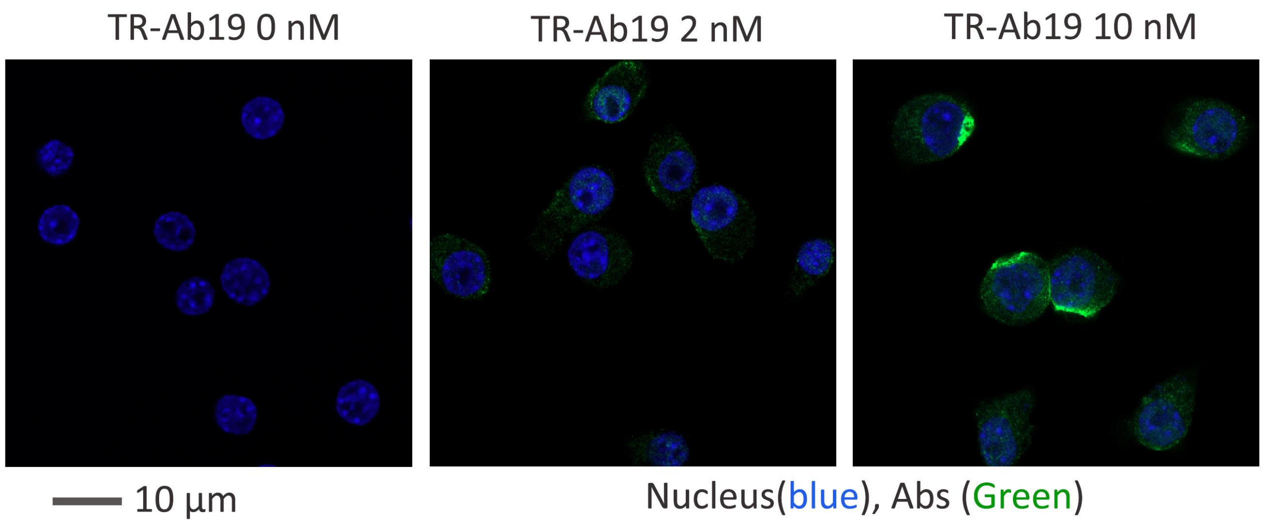


**Figure S2**. Confocal images of RAW264.7 macrophages incubated with TR-Ab19 reveal dose-dependent, saturable binding of TR-Ab19 to endogenous TREM2.

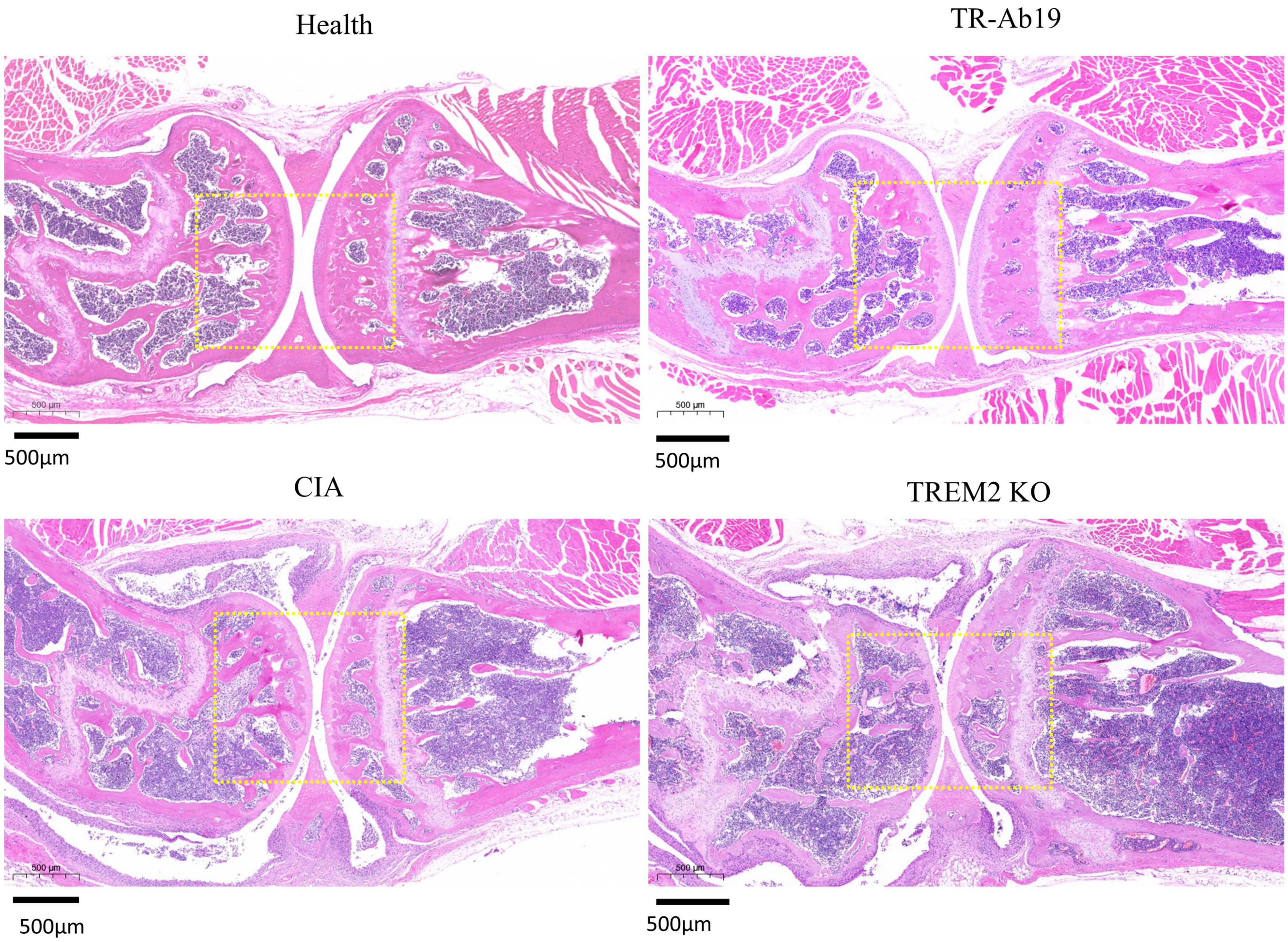


**Figure S3**. **Representative low-magnification H&E staining of knee joints from CIA cohorts.** H&E-stained sections of knee joints from healthy, TR-Ab19–treated CIA, CIA, and TREM2 KO CIA mice. CIA and especially TREM2 KO joints show marked synovial hyperplasia and inflammatory infiltration compared with healthy and TR-Ab19–treated mice, which preserve joint architecture: scale bars, 500 µm.

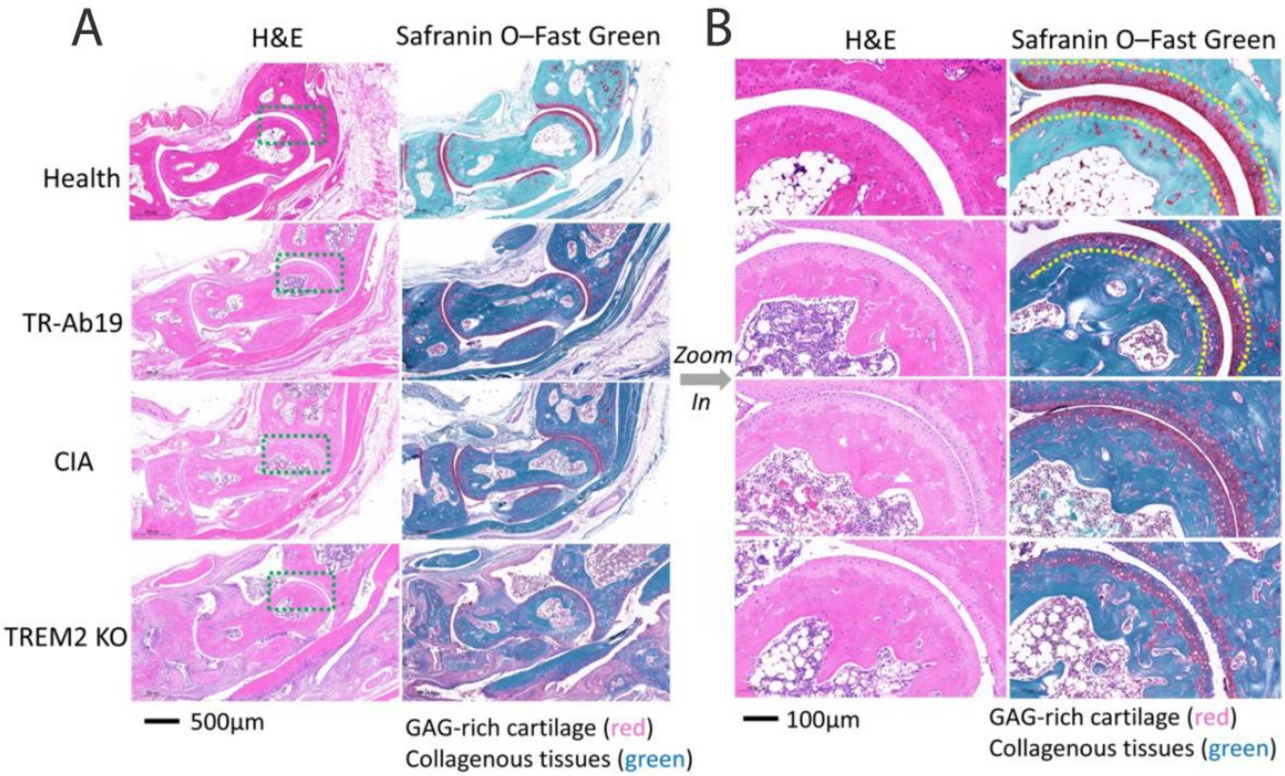


**Figure S4**. **TR-Ab19 preserves articular cartilage integrity in CIA joints. A,** Low-magnification H&E (left) and Safranin O–Fast Green (right) staining of ankle joints from healthy, TR-Ab19–treated CIA, CIA, and TREM2 KO CIA mice. Dashed boxes indicate regions used for higher-magnification views. **B,** Zoomed-in images of the boxed areas in **A**. Healthy and TR-Ab19–treated joints show smooth articular surfaces and intense Safranin O staining of glycosaminoglycan (GAG)-rich cartilage (red, yellow dotted line), with preserved cartilage thickness. CIA and TREM2 KO joints display pannus invasion, cartilage surface irregularity, and marked loss of GAG staining. Collagenous tissues are counterstained green. Scale bars, 500 μm (**A**) and 100 μm (**B**).

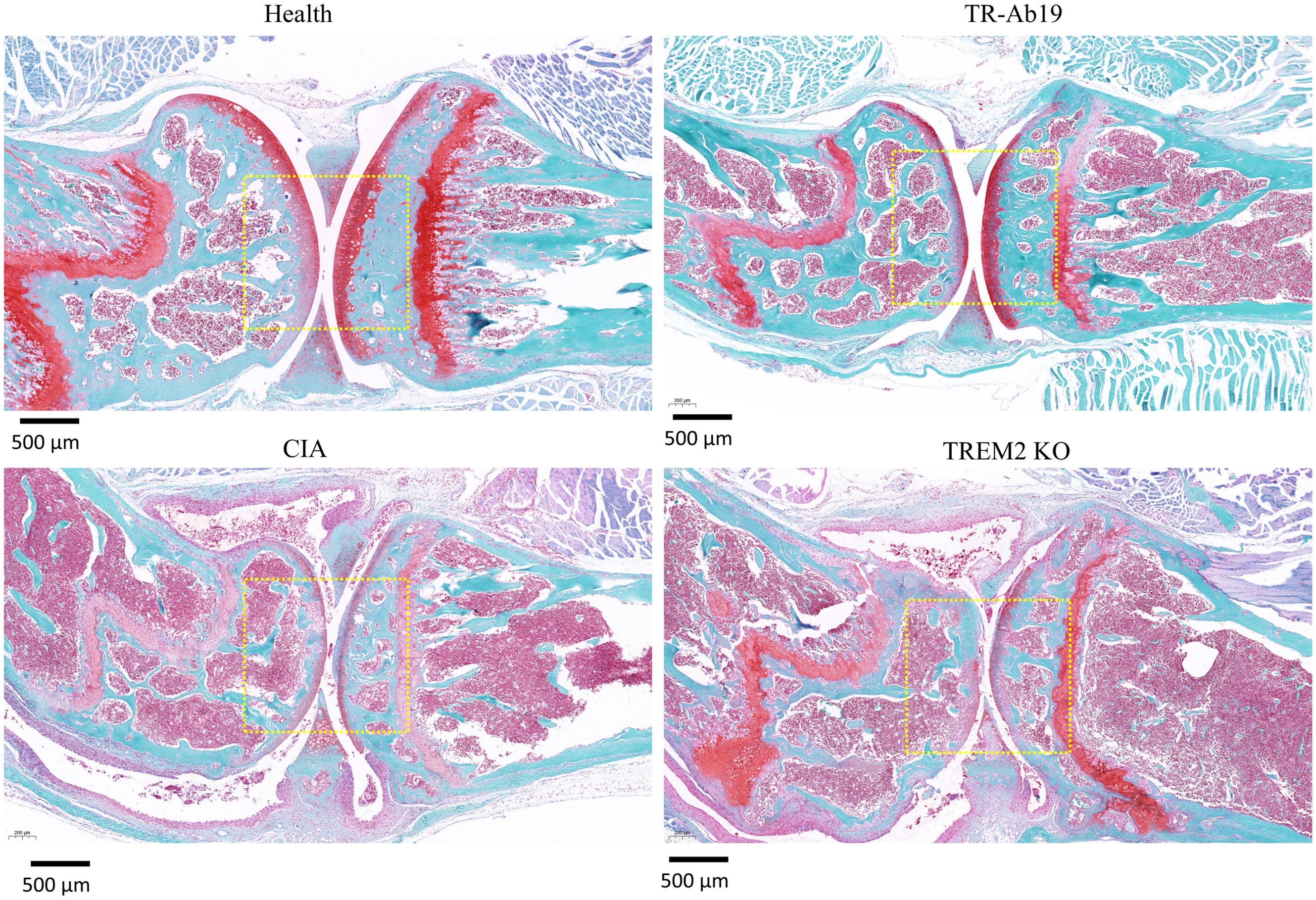


**Figure S5**. **Safranin O–Fast Green staining of knee joints from CIA cohorts.** Representative Safranin O–Fast Green–stained sections of knee joints from healthy, TR-Ab19–treated CIA, CIA, and TREM2 KO CIA mice. In healthy and TR-Ab19–treated knees, GAG-rich articular cartilage (red) is thick and continuous, with well-preserved joint surfaces, whereas CIA and especially TREM2 KO joints show marked loss of Safranin O staining, cartilage thinning, and surface irregularity, accompanied by pannus invasion. Collagenous tissues are counterstained green. Scale bars, 500 µm.

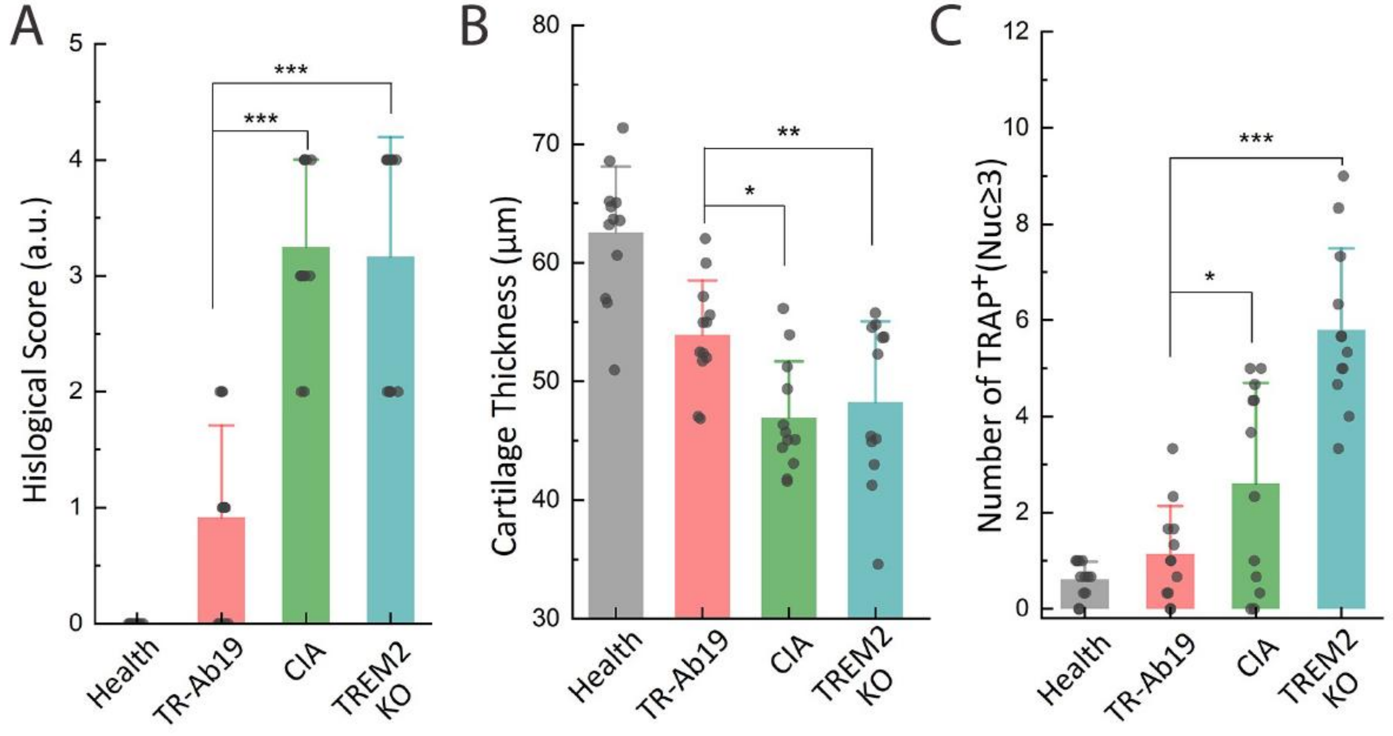


**Figure S6**. **Quantitative histomorphometric analysis of joint protection by TR-Ab19 in CIA mice.**

**A,** Histological arthritis scores of ankle joints from healthy, TR-Ab19–treated CIA, CIA, and TREM2 KO CIA mice, showing marked reduction of joint pathology after TR-Ab19 treatment. **B,** Articular cartilage thickness measured in Safranin O–Fast Green–stained sections, indicating partial preservation of cartilage in TR-Ab19–treated joints compared with CIA and TREM2 KO mice. **C,** Numbers of $TRAP^+$ multinucleated (nuc ≥ 3) osteoclasts per joint section, demonstrating decreased osteoclast burden with TR-Ab19 and increased osteoclastogenesis in CIA and TREM2 KO groups. *P < 0.05, **P < 0.01, ***P < 0.001.

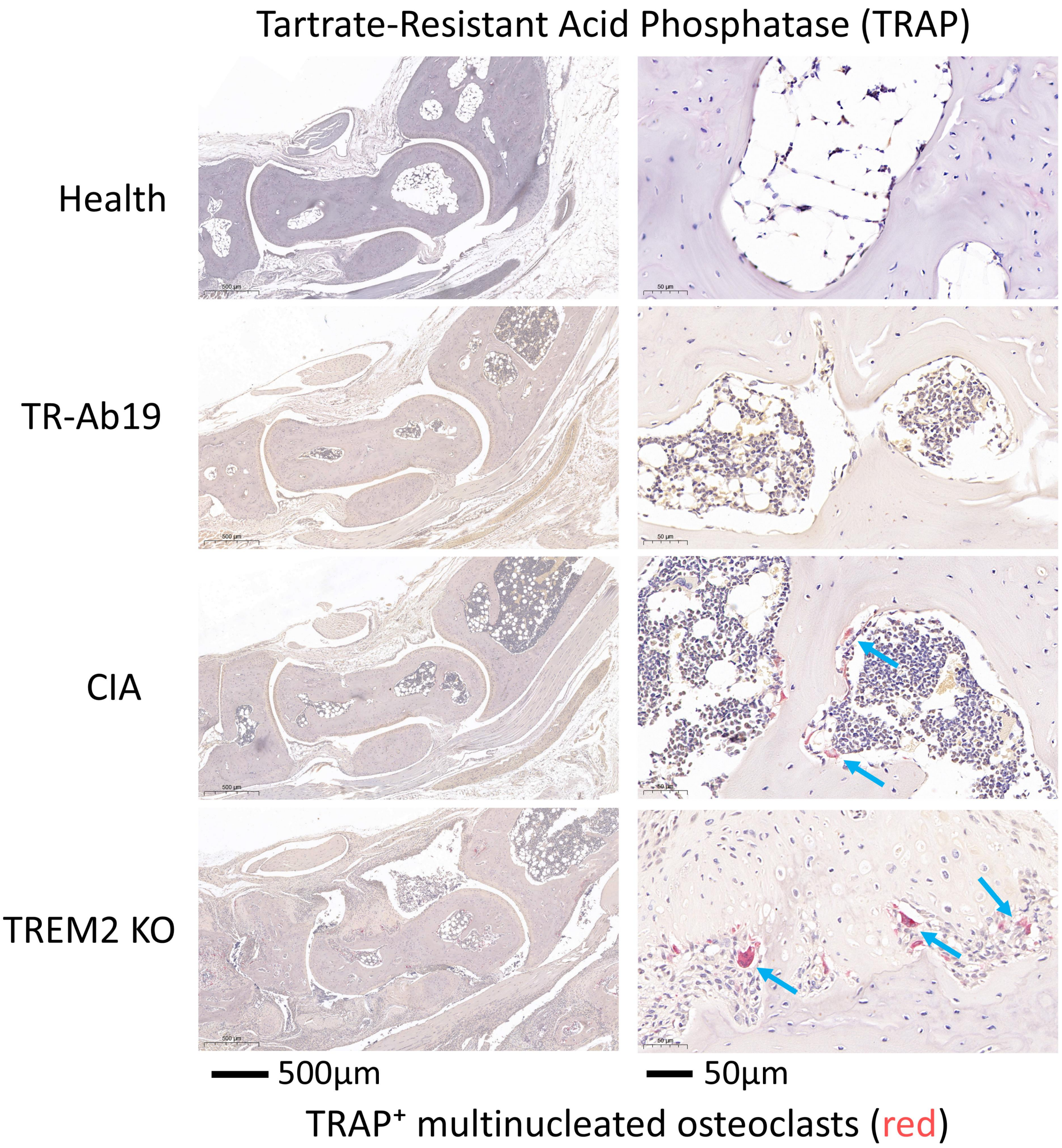


**Figure S7**. **TR-Ab19 reduces osteoclast accumulation in CIA joints.** Representative tartrate-resistant acid phosphatase (TRAP) staining of ankle joints from healthy, TR-Ab19–treated CIA, CIA, and TREM2 KO CIA mice. Left column, low-magnification views; right column, zoomed-in images of subchondral bone surfaces. $TRAP^+$ multinucleated osteoclasts (red, blue arrows) are rare in healthy joints, increased in CIA mice, and further enriched in TREM2 KO mice, whereas TR-Ab19 treatment markedly reduces $TRAP^+$ osteoclasts. Scale bars, 500 μm (left) and 50 μm (right).

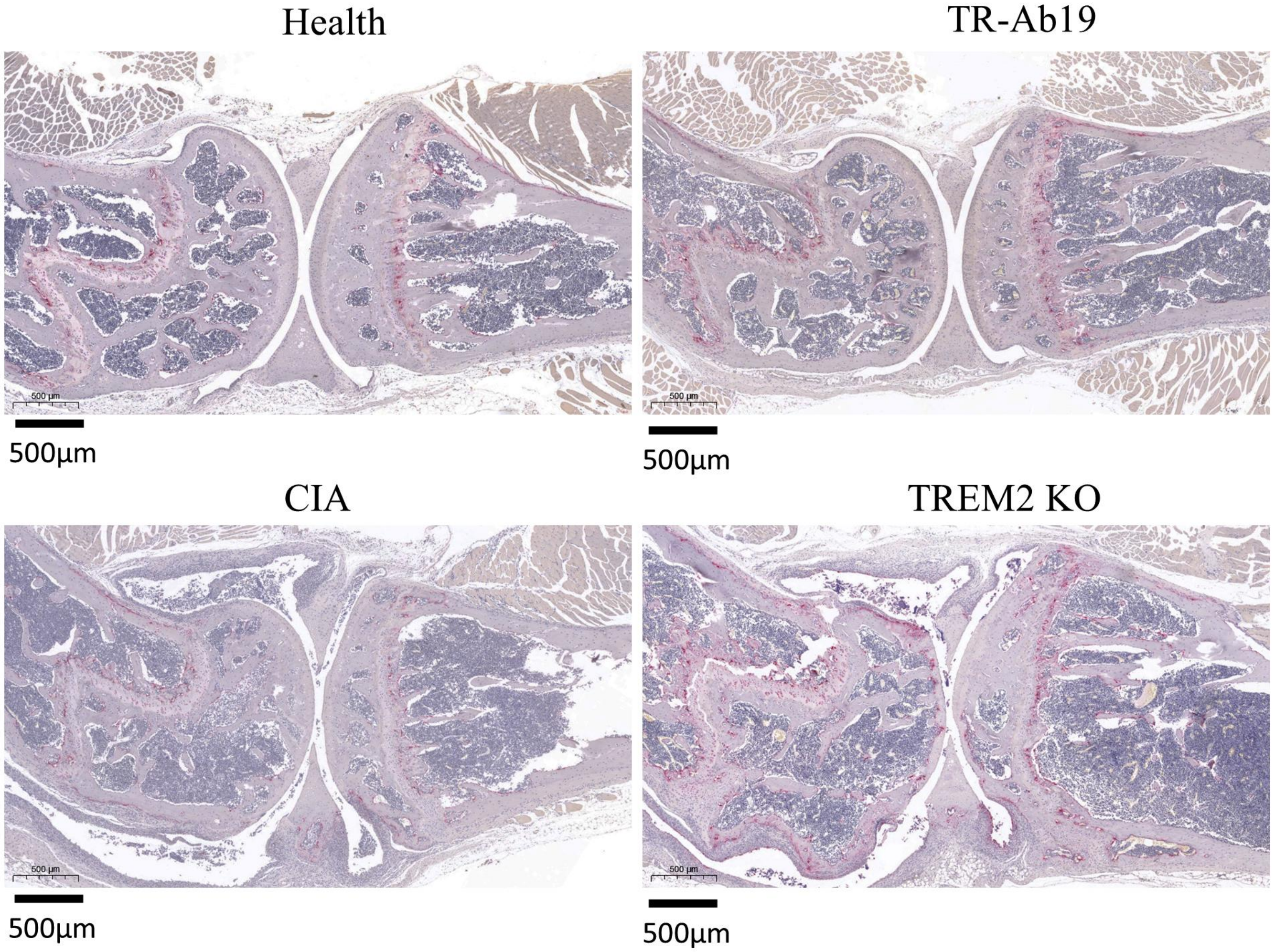


**Figure S8**. **Representative TRAP staining of ankle joints from CIA cohorts.** Tartrate-resistant acid phosphatase (TRAP) staining of ankle joints from healthy, TR-Ab19–treated CIA, CIA, and TREM2 KO CIA mice. Red signal marks $TRAP^+$ multinucleated osteoclasts along subchondral and trabecular bone surfaces. CIA and especially TREM2 KO joints show increased and expanded $TRAP^+$ osteoclasts compared with healthy and TR-Ab19–treated mice, consistent with the quantitative analysis in Fig. 2i. Scale bars, 500 µm.

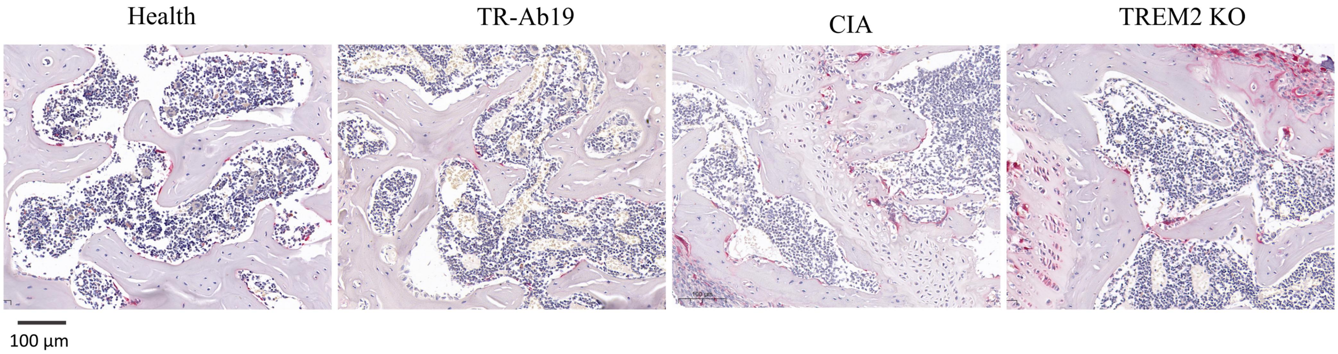


**Figure S9**. **TR-Ab19 reduces osteoclast accumulation in knee joints of CIA mice.** Representative tartrate-resistant acid phosphatase (TRAP)–stained sections of knee joints from healthy, TR-Ab19–treated CIA, CIA, and TREM2 KO CIA mice. TRAP$^+$ multinucleated osteoclasts (red) along subchondral and trabecular bone surfaces are sparse in healthy and TR-Ab19–treated joints but increased in CIA and further enriched in TREM2 KO knees: scale bar, 100 µm.

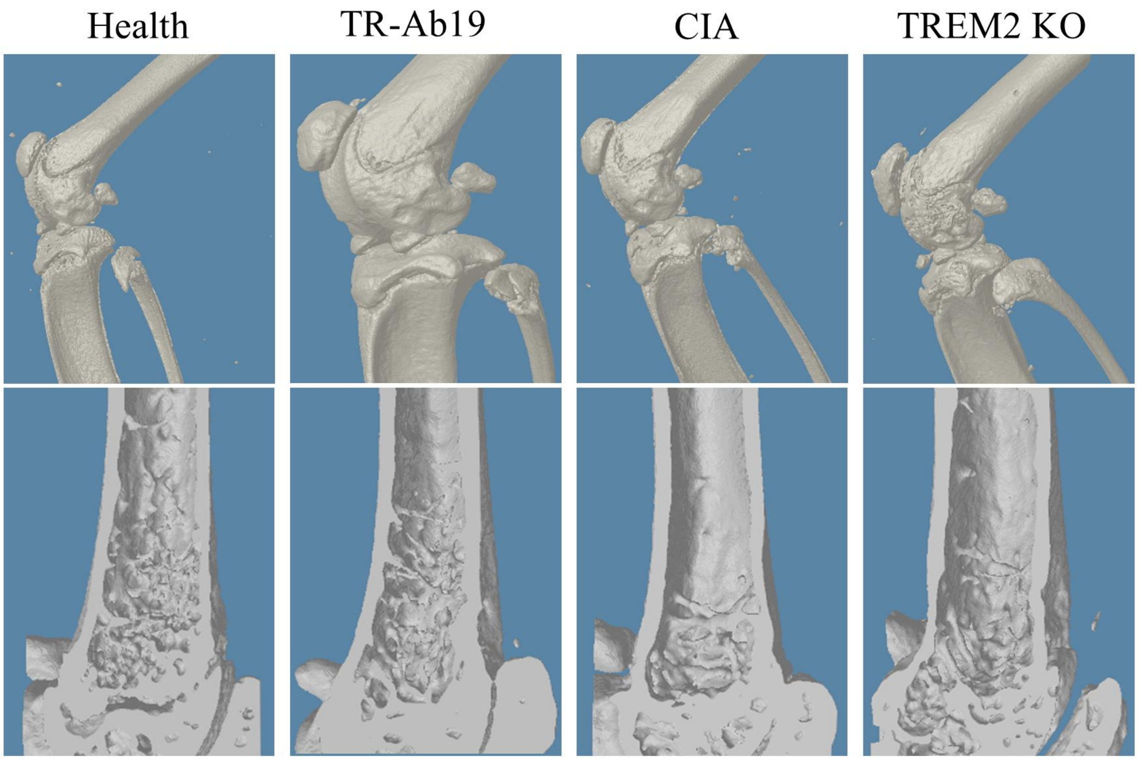


**Figure S10**. **Micro-CT assessment of joint bone destruction in CIA cohorts.** Representative 3D micro-CT reconstructions of murine knee joints (top row) and distal tibial metaphyses (bottom row) from healthy, TR-Ab19–treated CIA, CIA, and TREM2 KO CIA mice. Compared with healthy and TR-Ab19–treated mice, CIA and Trem2$^{-}$/$^{-}$ CIA joints show osteophyte formation, cortical irregularities, and reduced trabecular bone, consistent with more severe structural pathology under these conditions.

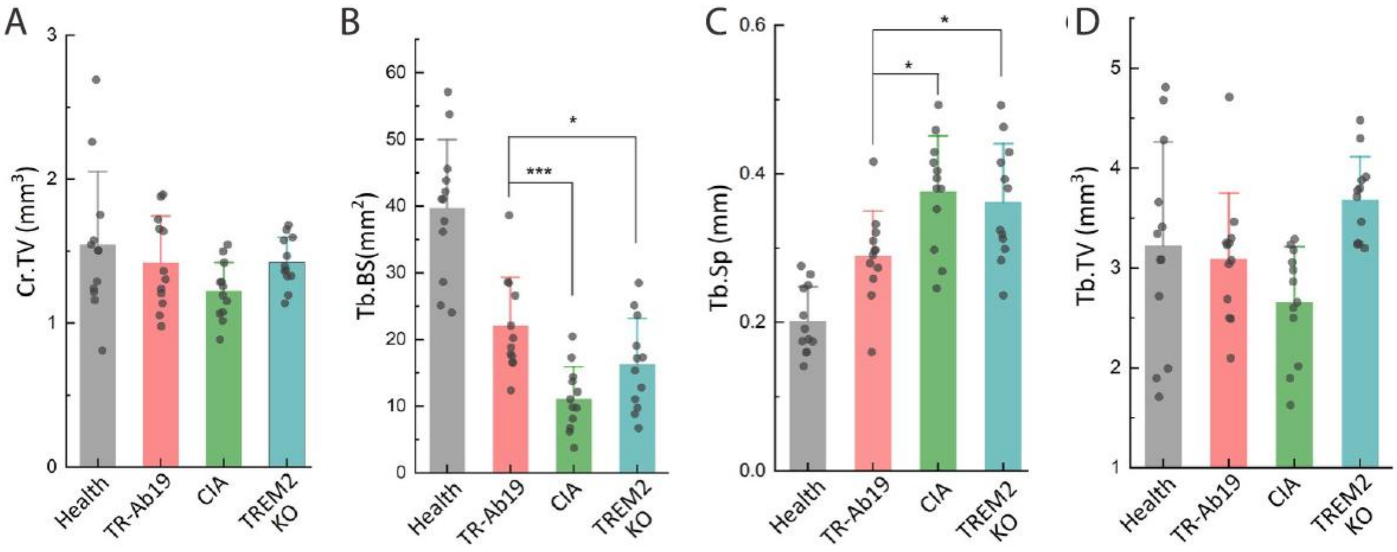


**Figure S11**. **Quantitative micro-CT analysis of knee joint bone architecture in CIA cohorts.** Micro-CT–derived structural parameters of distal tibial metaphyses from healthy, TR-Ab19–treated CIA, CIA, and TREM2 KO CIA mice corresponding to the 3D images in Fig. S9. **A,** Cortical tissue volume (Cr.TV). **B,** Trabecular bone surface (Tb.BS). **C,** Trabecular separation (Tb.Sp). **D,** Trabecular tissue volume (Tb.TV). CIA and TREM2 KO mice show reduced Tb.BS and increased Tb.Sp, indicative of trabecular loss and rarefaction, whereas TR-Ab19 treatment partially preserves trabecular structure. *P < 0.05, ***P < 0.001.

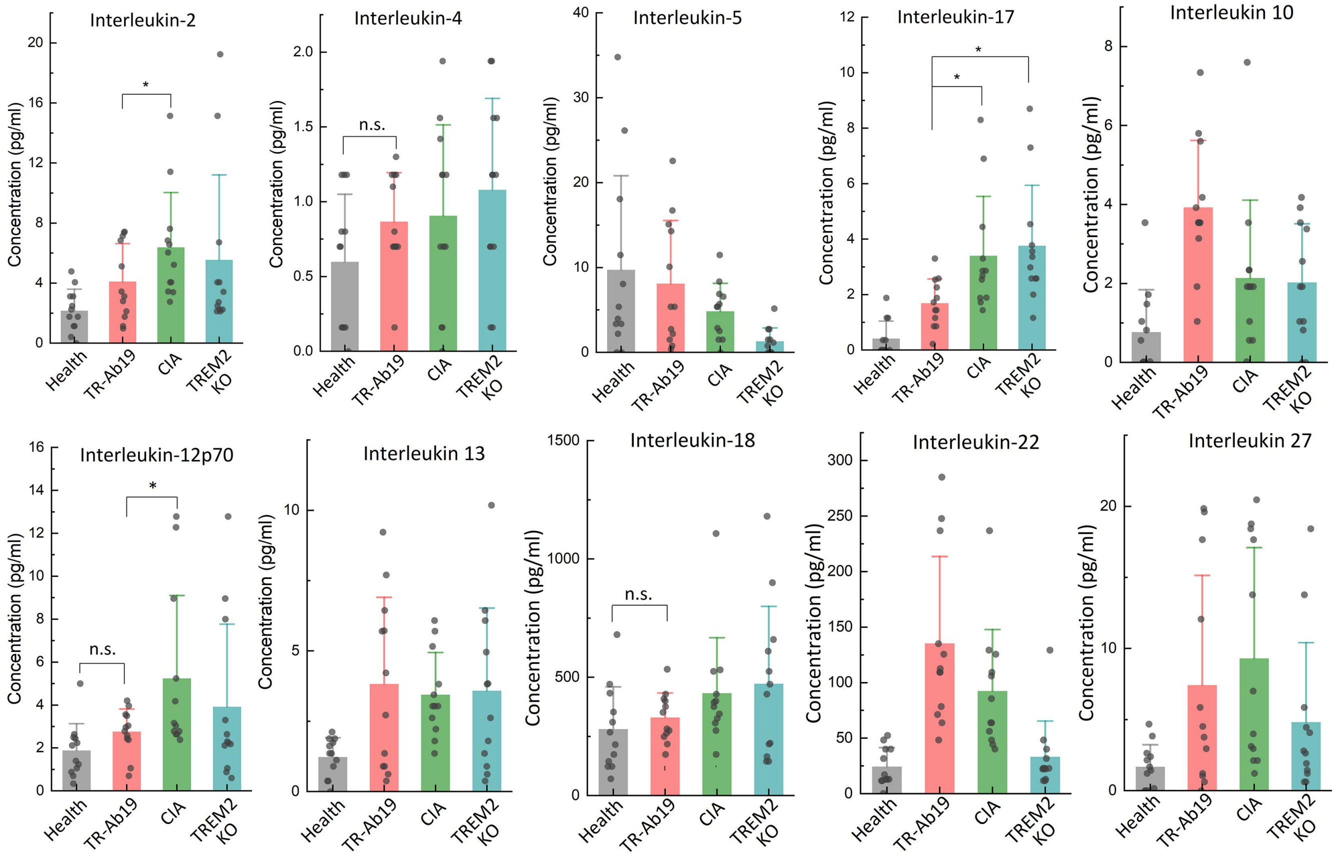


**Figure S12**. **Extended serum cytokine profiling in CIA cohorts.** Multiplex bead–based immunoassay of additional serum interleukins in healthy, TR-Ab19–treated CIA, CIA, and TREM2 KO CIA mice corresponding to Fig. 3c. Shown are concentrations of IL-2, IL-4, IL-5, IL-17, IL-10, IL-12p70, IL-13, IL-18, IL-22 and IL-27. CIA and TREM2 KO mice display elevated levels of several pro-inflammatory cytokines (for example IL-2, IL-17 and IL-12p70), whereas TR-Ab19 treatment tends to reduce or normalize these mediators relative to CIA. n.s., not significant; *$P < 0.05$.

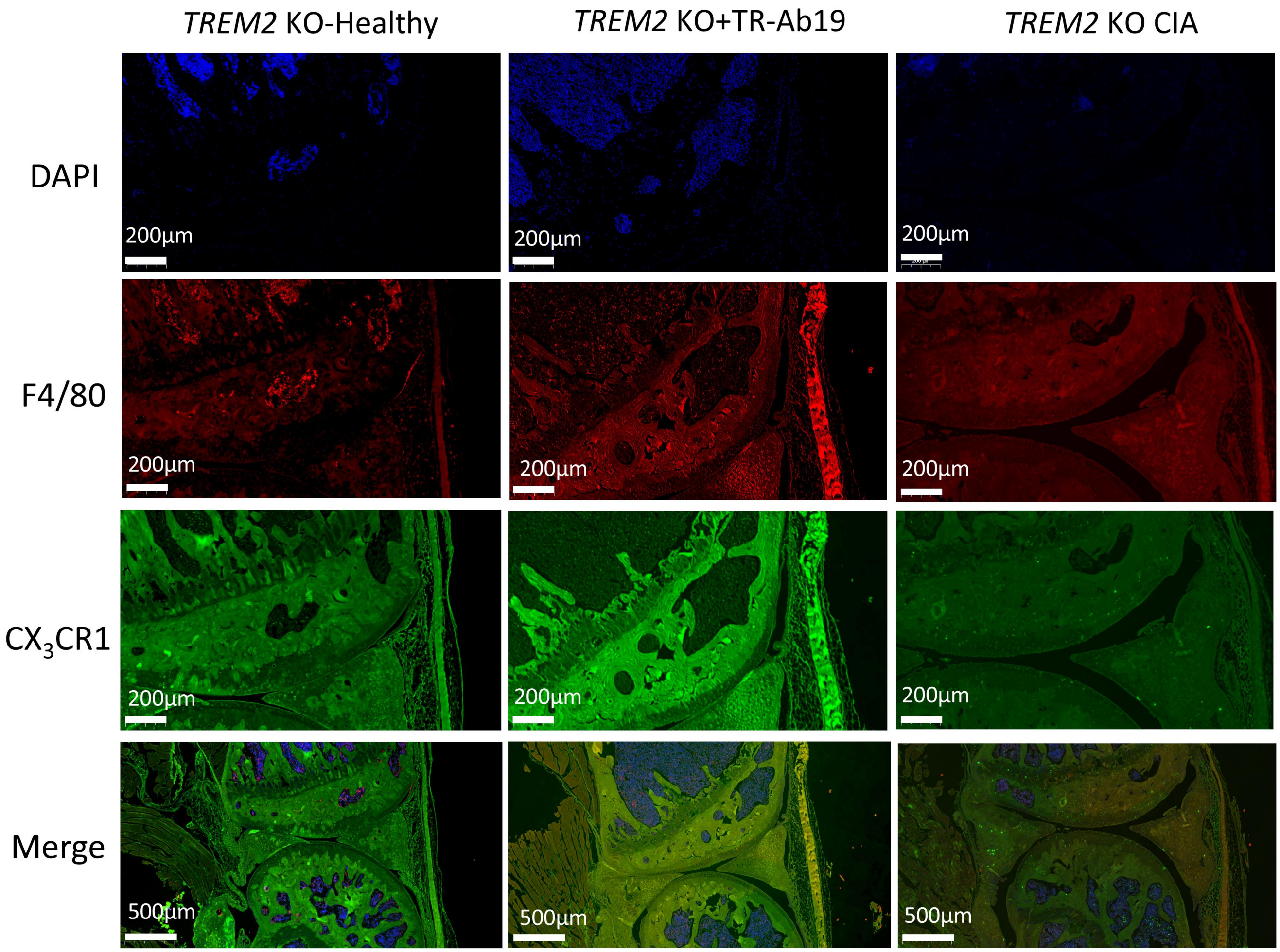


**Figure S13**. **Severe disruption of the synovial CX₃CR1$^{High}$ macrophage barrier in TREM2-deficient mice.** Representative immunofluorescence images of ankle joints from TREM2 KO–healthy, TREM2 KO + TR-Ab19–treated CIA, and TREM2 KO CIA mice stained for nuclei (DAPI, blue), F4/80 (red), and $CX_3CR1$ (green). In TREM2 KO–healthy joints, F4/80$^+$$CX_3CR1^+$ macrophages already fail to organize into the continuous synovial lining layer described in wild-type mice, and this $CX_3CR1^+$ barrier is further fragmented and disorganized in TREM2 KO CIA. TR-Ab19 administration does not restore the lining in the absence of TREM2, consistent with a TREM2-dependent mechanism. Scale bars, 200 µm (single channels) and 500 µm (merged images).

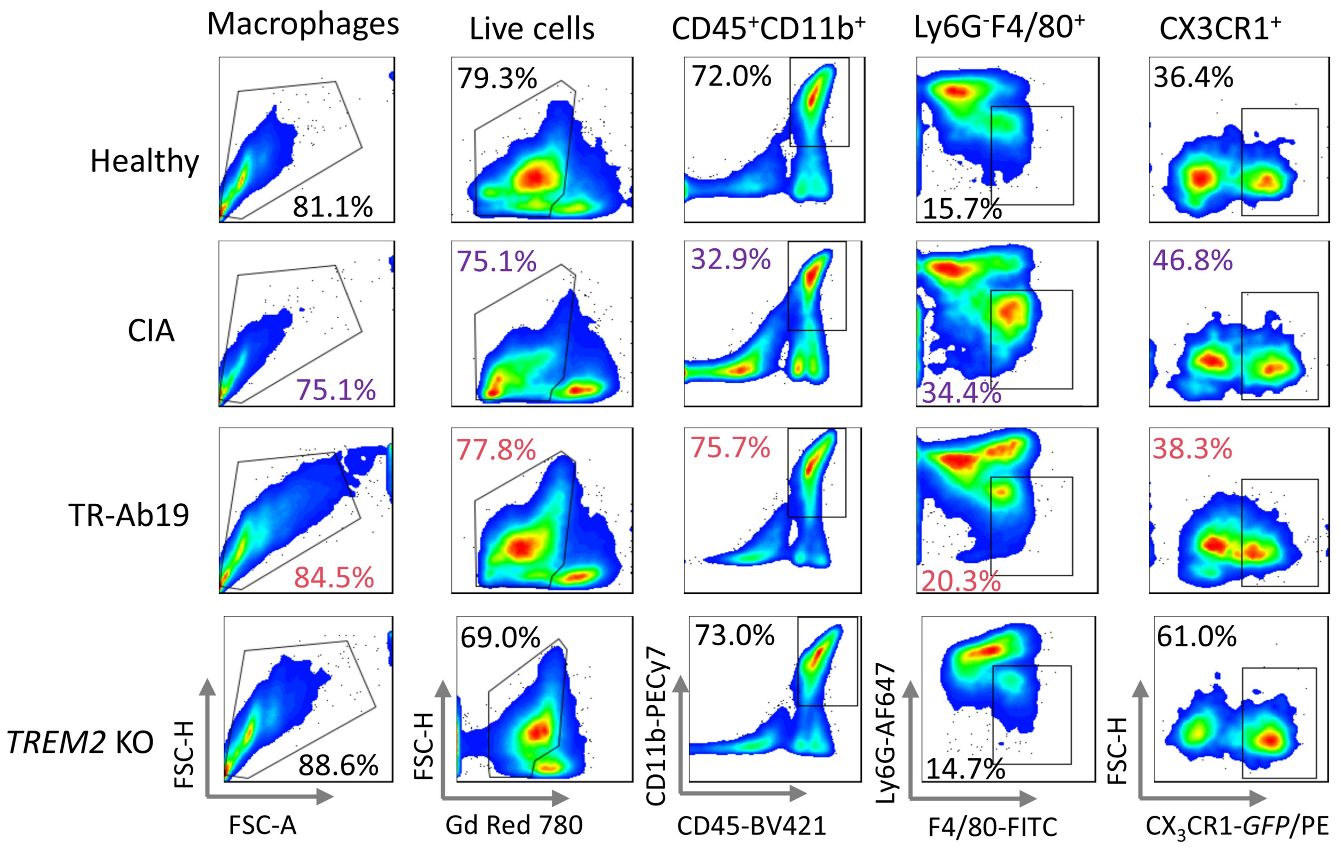


**Figure S14**. **Multicolor flow-cytometry analysis of synovial macrophages in CIA mice.** Representative flow-cytometry plots of synovial cell suspensions from ankle joints of Healthy, CIA, CIA + TR-Ab19, and TREM2 KO CIA mice. From left to right, panels show gating of total macrophage-enriched cells (FSC/SSC), live cells (Gd Red 780$^-$), CD45$^+$CD11b$^+$ myeloid cells, Ly6G$^-$F4/80$^+$ macrophages, and $CX_3CR1^+$ macrophages. Percentages indicate the fraction of cells within each gate. Compared with untreated CIA mice, TR-Ab19 treatment increases the proportion of live CD45$^+$CD11b$^+$ and F4/80$^+$ macrophages and maintains $CX_3CR1^+$ macrophages, whereas TREM2 KO joints display altered myeloid composition and expansion of $CX_3CR1^+$ cells.

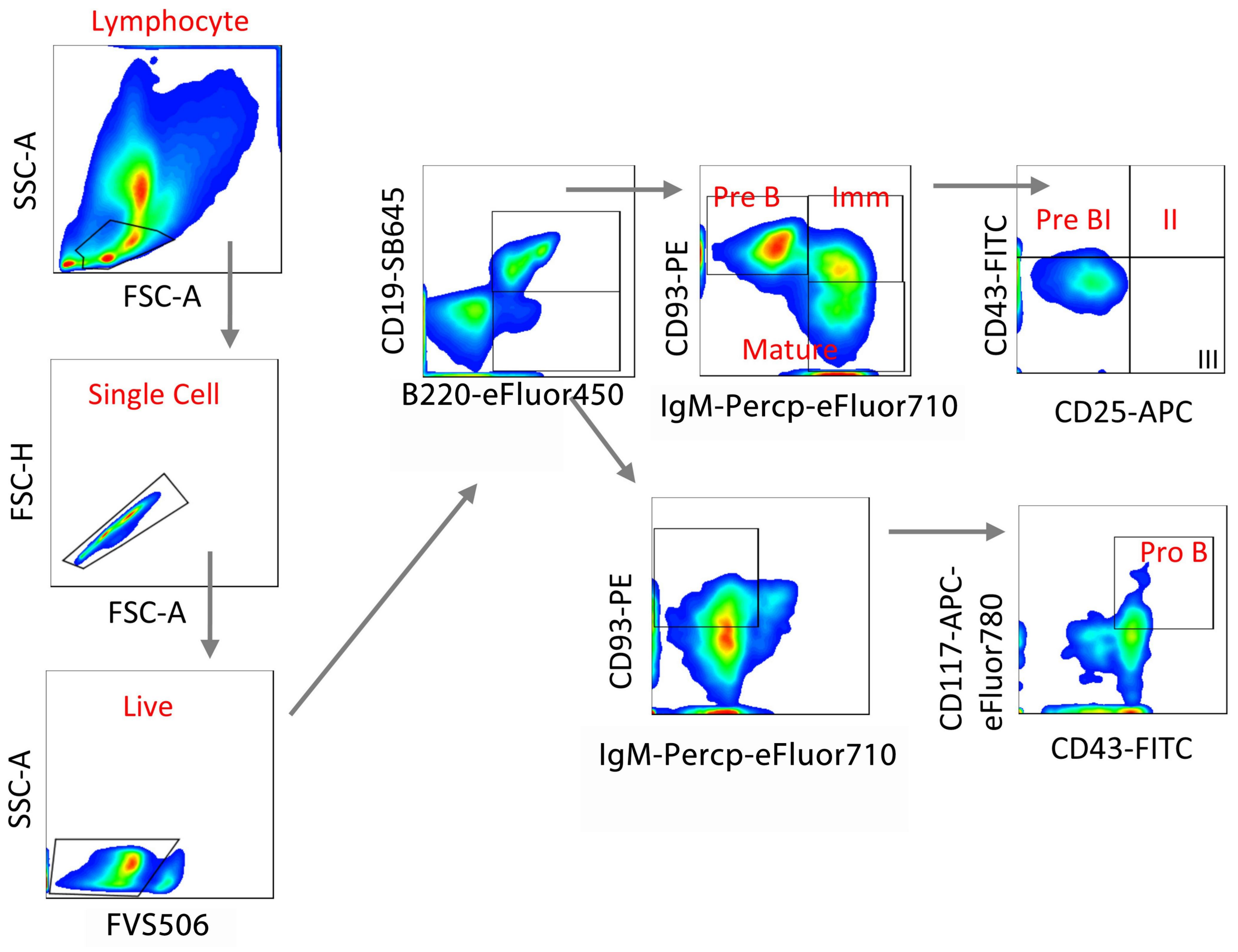


**Figure S15**. **Flow-cytometry gating strategy for bone marrow B-cell development.** Representative gating of bone marrow cells used to quantify B-cell subsets in Fig. 5h–j. Lymphocytes were first identified by FSC-A/SSC-A, followed by singlet discrimination (FSC-H vs FSC-A) and exclusion of dead cells ($BV510^+$). Live $CD19^+B220^+$ B-lineage cells were then subdivided by CD93 and surface IgM into pre-B, immature and mature B cells. Pre-B cells were further separated into Pre-BI, Pre-BII, and Pre-BIII fractions by CD43 and CD25 expression, while Pro-B cells were defined as $CD93^+IgM^-CD117^+CD43^+$ cells.

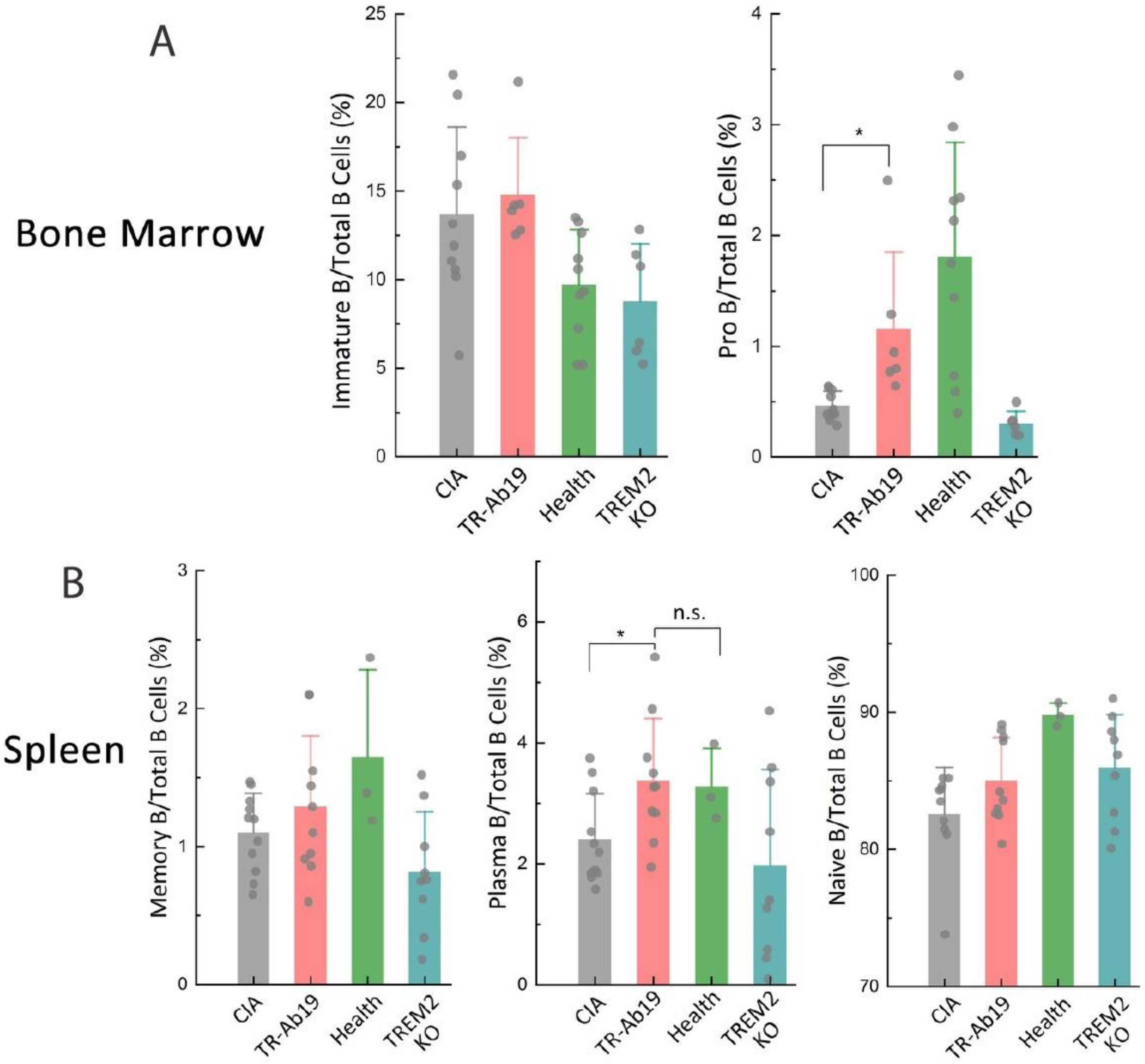


**Figure S16**. **Additional analysis of B-cell subsets in bone marrow and spleen. A, Bone marrow.**

Frequencies of immature B cells (left) and pro-B cells (right) among total CD19⁺B220⁺ bone marrow B cells in CIA, CIA + TR-Ab19, healthy and TREM2 KO mice. TR-Ab19 partially normalizes the excess pro-B compartment seen in CIA, approaching healthy levels. **B, Spleen.** Frequencies of memory B cells (left), plasma B cells (middle) and naïve B cells (right) among total splenic B cells in the same treatment groups. TR-Ab19 tends to increase plasma and naïve B cells relative to CIA, while TREM2 KO mice show reduced plasma B cells. n.s., not significant; *P < 0.05.

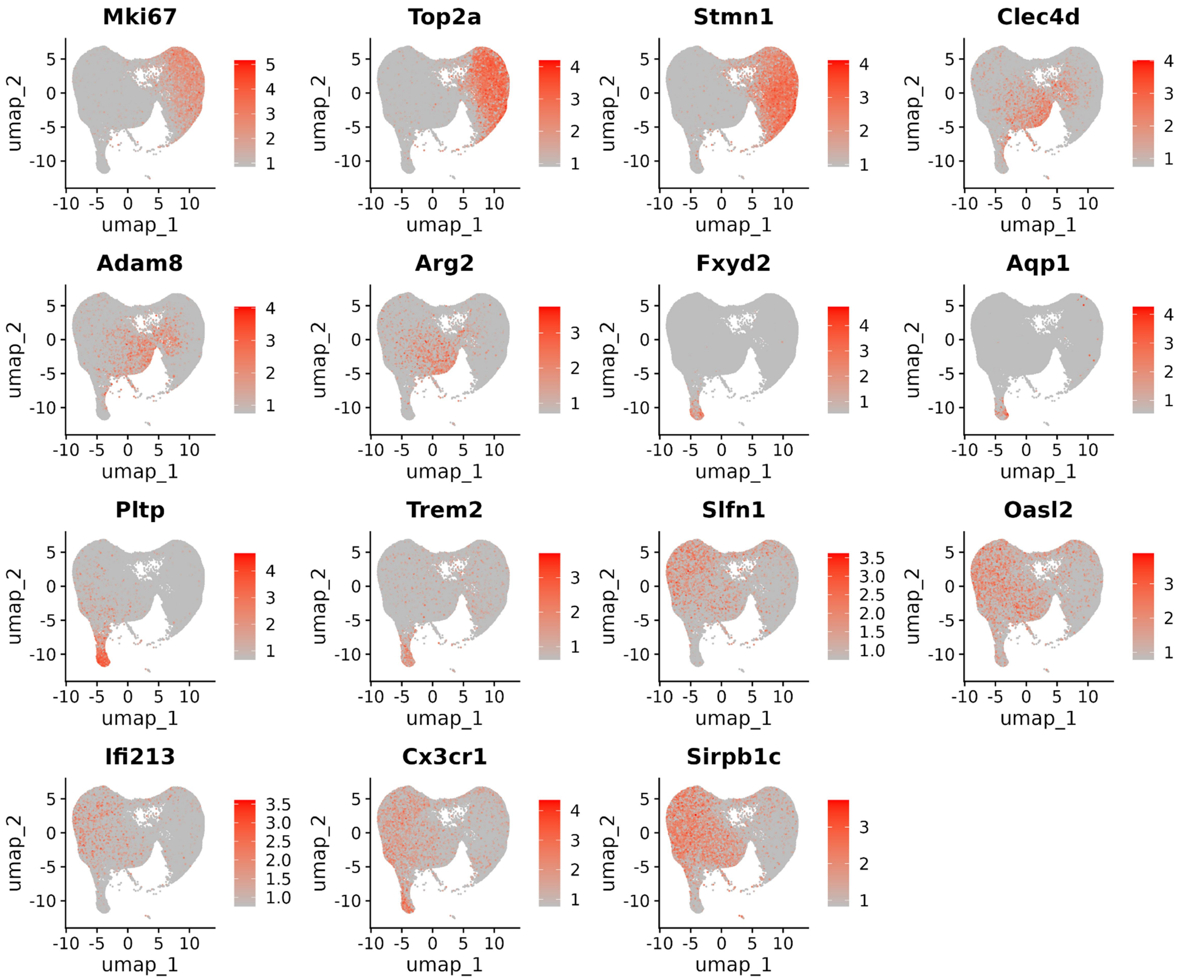


**Figure S17. Marker-gene expression supporting synovial macrophage-state annotations in Fig. 6.**

Integrated UMAP of synovial macrophages from Healthy, CIA, CIA + TR-Ab19, and Trem2$^{-}$/$^{-}$ CIA joints after quality control and batch correction (Supplementary Methods). Feature plots show log-normalized expression of representative markers used to annotate major macrophage states in Fig. 6, including proliferation/cell-cycle markers (e.g., Top2a, Mki67, Stmn1), inflammatory markers (e.g., Clec4d, Adam8, Arg2), reparative markers (e.g., Aqp1), and barrier-associated markers (e.g., CX$_3$CR1 and interferon-associated genes as indicated in the plot). Related to Fig. 6A–B. Abbreviations: CIA, collagen-induced arthritis; UMAP, uniform manifold approximation and projection.

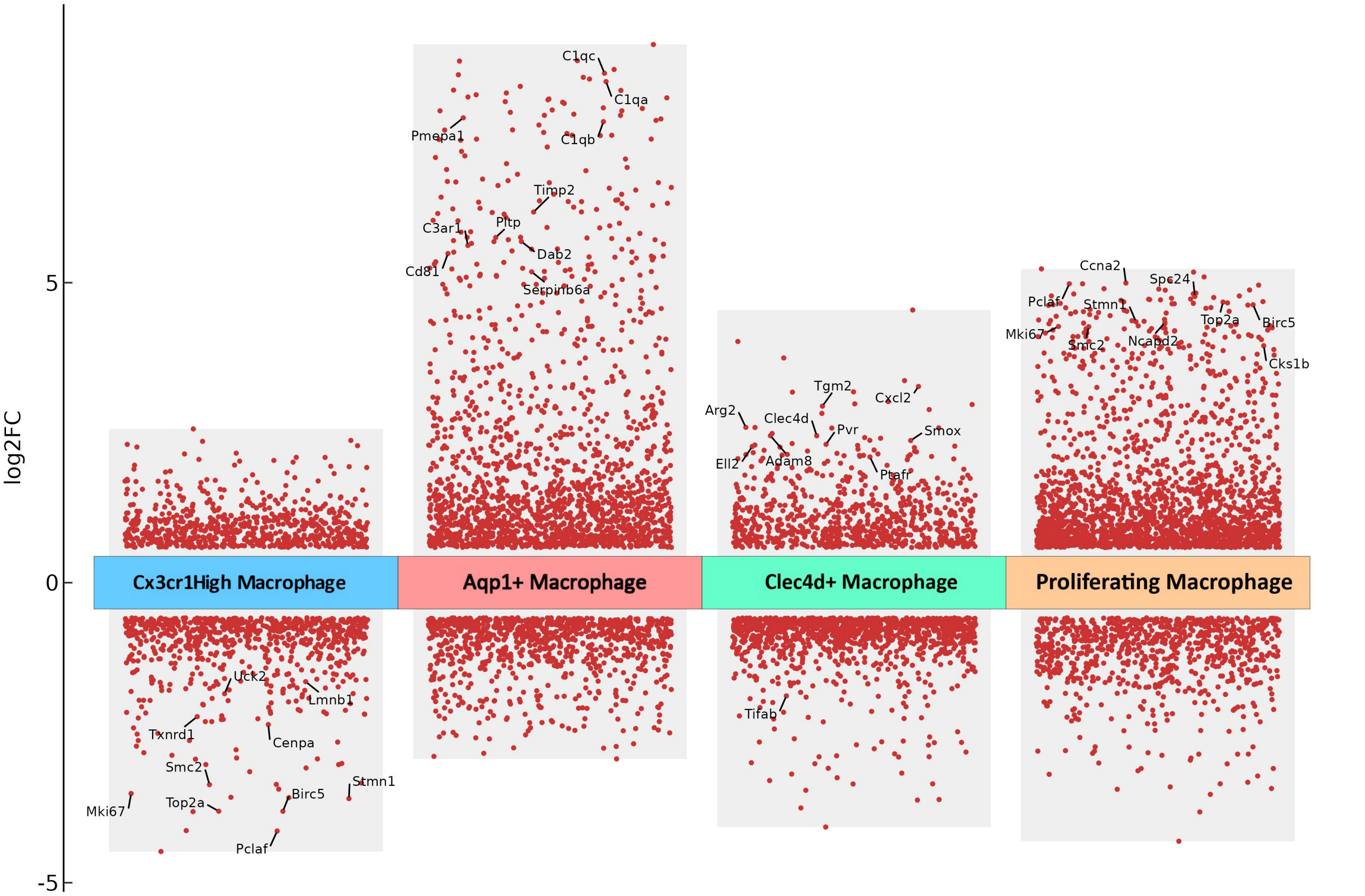


**Figure S18. Cluster-specific differentially expressed genes defining synovial macrophage subsets in Fig. 6.** Dot plot summarizing differentially expressed genes (DEGs) for the four macrophage subsets shown in Fig. 6 ($CX_3CR1^{high}$ barrier-like, $Aqp1^{+}$ reparative, $Clec4d^{+}$ inflammatory, and proliferating macrophages). DEGs were computed using Seurat FindAllMarkers/FindMarkers with Wilcoxon rank-sum testing and multiple-testing correction as described (Supplementary Methods). Genes are ranked by log2 fold change (cluster vs. all other synovial macrophages). Dot size indicates the fraction of cells expressing each gene and dot color indicates average log-normalized expression. "Significant" DEGs denote adjusted P < 0.05 after correction (as indicated in the plot). Related to Fig. 6A and Fig. S17. Abbreviations: DEG, differentially expressed gene; CIA, collagen-induced arthritis.

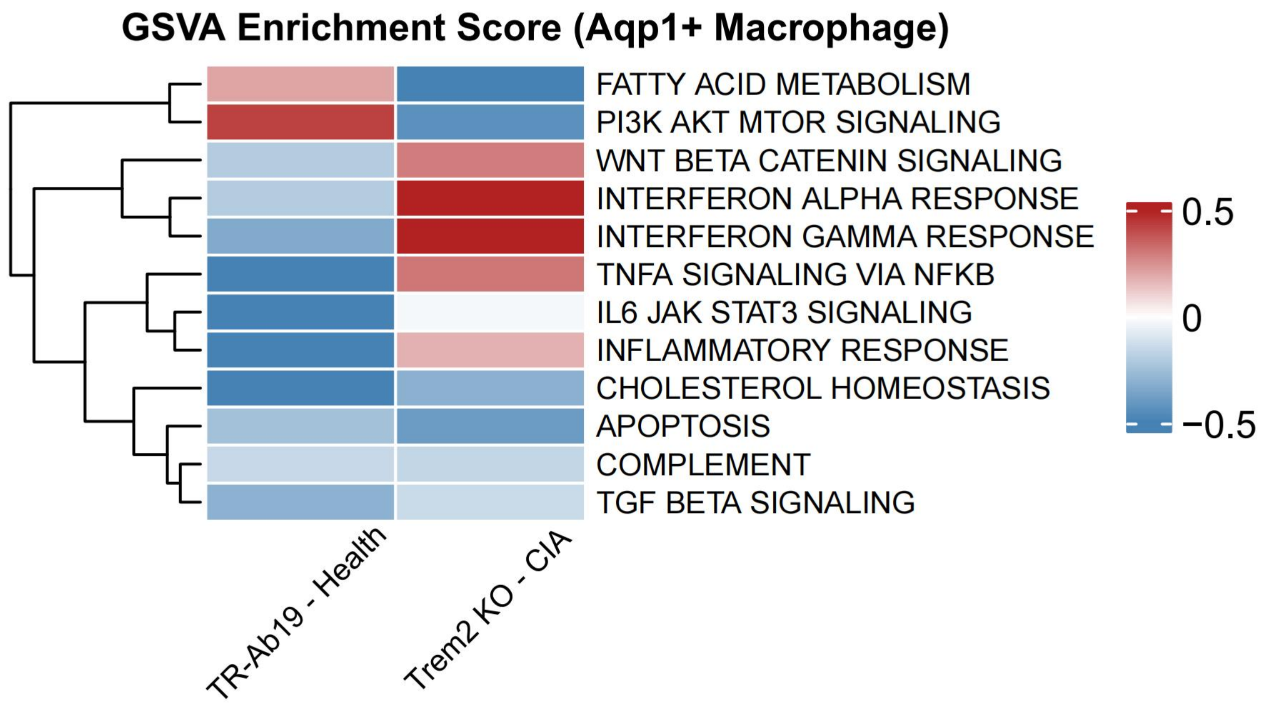


**Figure S19. Hallmark pathway activity in Aqp1⁺ reparative synovial macrophages across conditions.** GSVA heat map of MSigDB Hallmark gene sets calculated in Aqp1⁺ macrophages from Healthy, CIA, CIA + TR-Ab19, and Trem2⁻/⁻ CIA joints. Colors indicate scaled GSVA scores (per-row normalization as shown in the heat map). Gene-set definitions and computational parameters are described in Supplementary Methods. Related to Fig. 6G and Fig. S17–S18. Abbreviations: GSVA, gene set variation analysis; MSigDB, Molecular Signatures Database; CIA, collagen-induced arthritis.

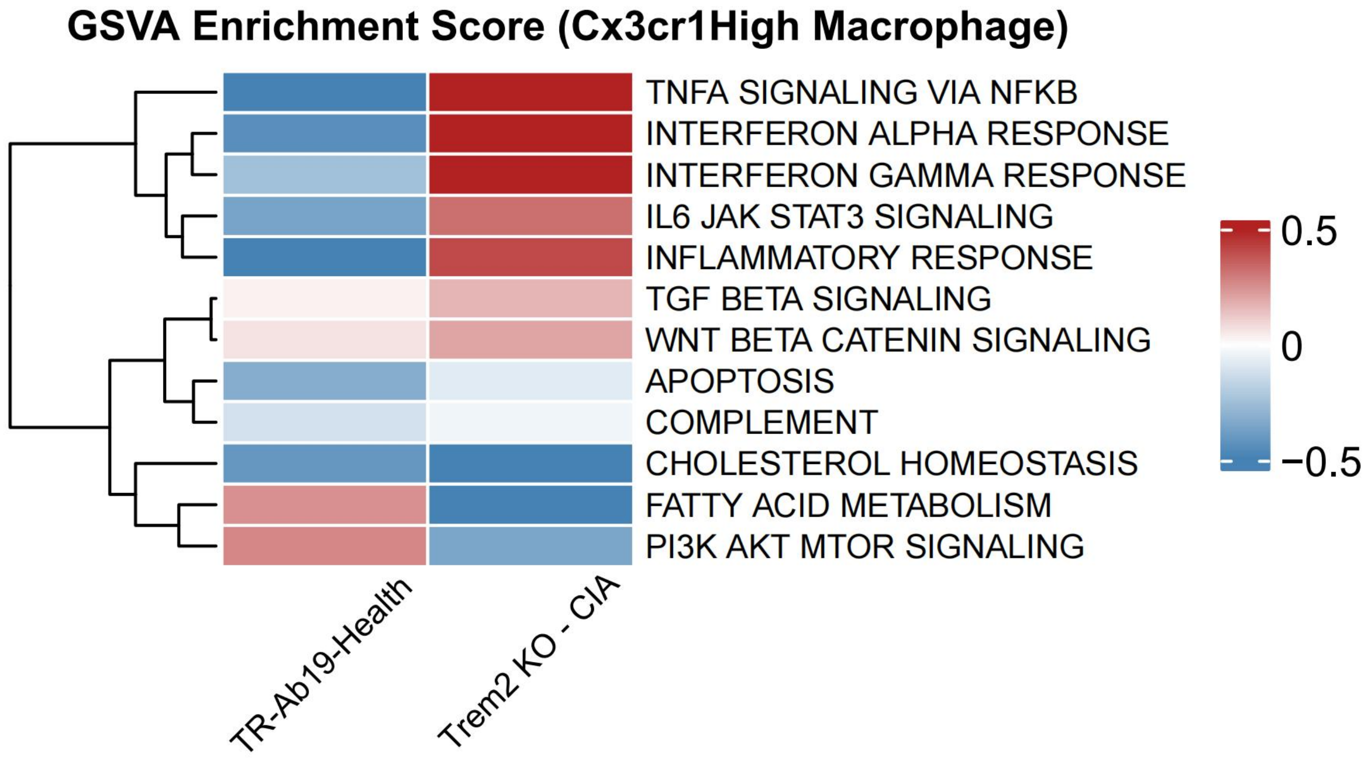


**Figure S20. Hallmark pathway activity in $CX_3CR1^{high}$ barrier-like synovial macrophages across conditions.** GSVA heat map of MSigDB Hallmark gene sets calculated in $CX_3CR1^{high}$ ("barrier-like") macrophages from Healthy, CIA, CIA + TR-Ab19, and $Trem2^{-}/^{-}$ CIA joints. Colors indicate scaled GSVA scores (per-row normalization as shown). Gene-set definitions and computational parameters are described in Supplementary Methods. Related to Fig. 6G and Fig. S17–S19. Abbreviations: GSVA, gene set variation analysis; CIA, collagen-induced arthritis.

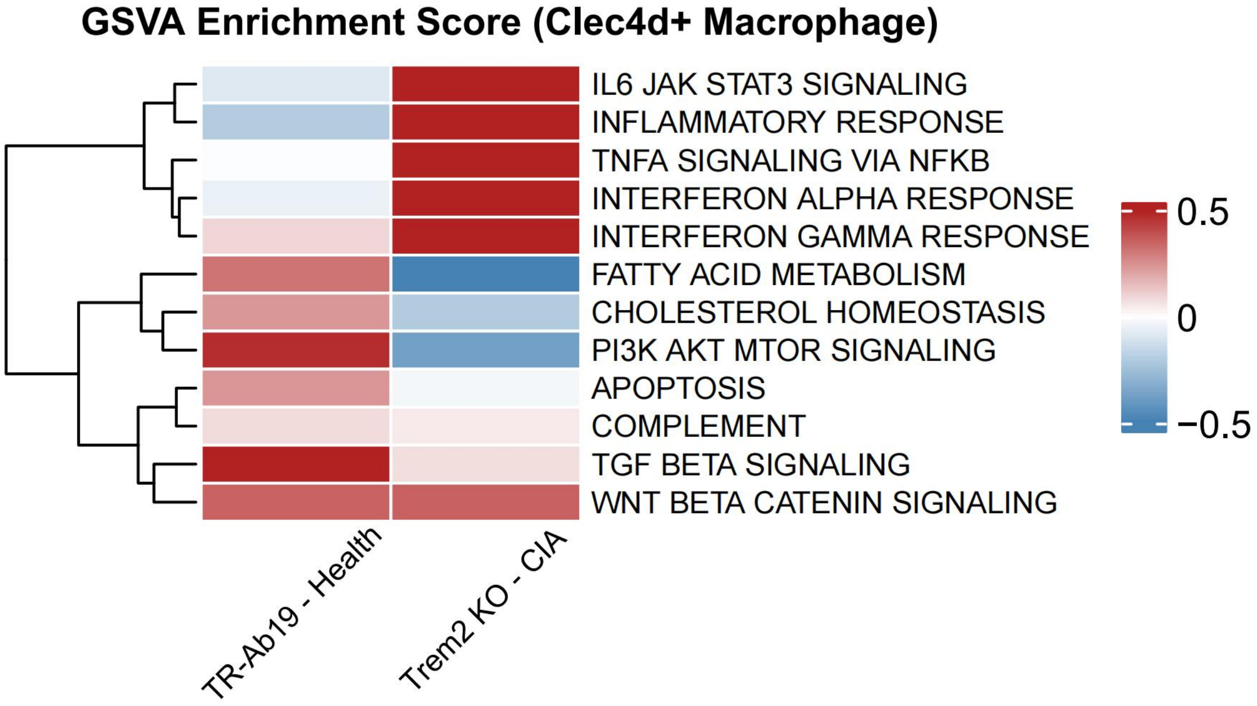


**Figure S21. Hallmark pathway activity in Clec4d⁺ inflammatory synovial macrophages across conditions.** GSVA heat map of MSigDB Hallmark gene sets calculated in Clec4d⁺ macrophages from Healthy, CIA, CIA + TR-Ab19, and Trem2⁻/⁻ CIA joints. Colors indicate scaled GSVA scores (per-row normalization as shown). Gene-set definitions and computational parameters are described in Supplementary Methods. Related to Fig. 6G and Fig. S17–S20. Abbreviations: GSVA, gene set variation analysis; CIA, collagen-induced arthritis.

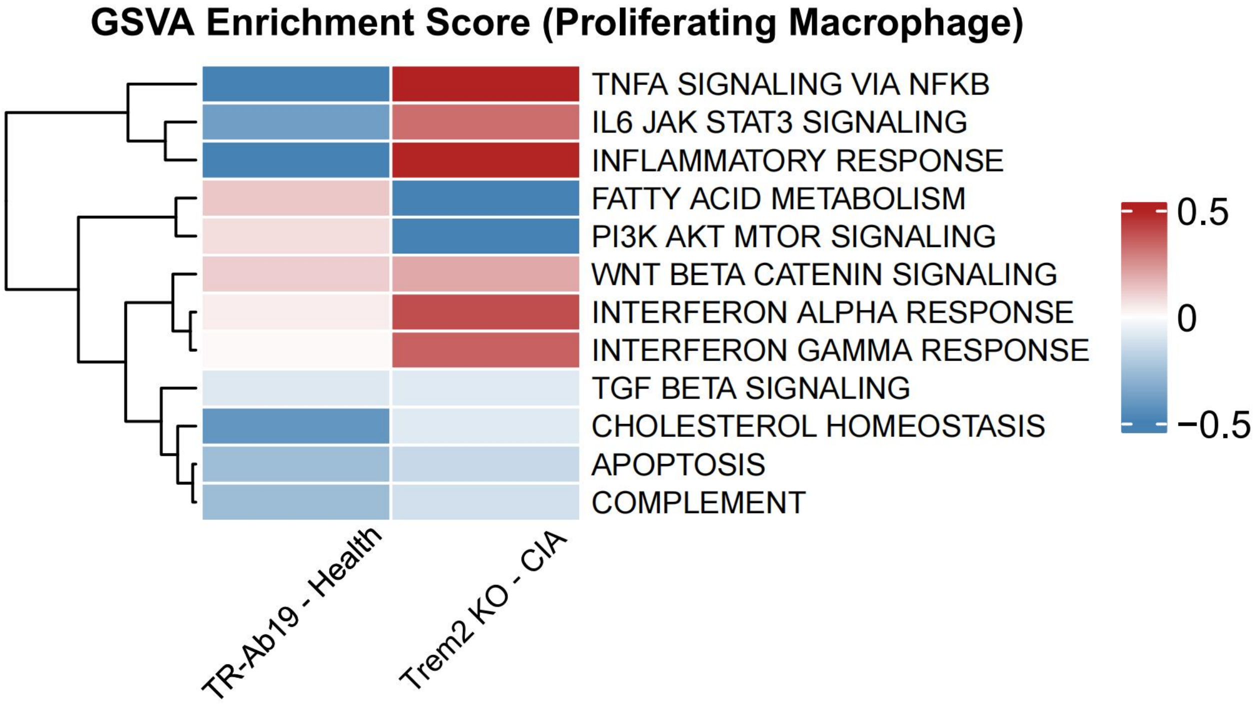


**Figure S22. Hallmark pathway activity in proliferating synovial macrophages across conditions.**

GSVA heat map of MSigDB Hallmark gene sets calculated in proliferating macrophages from Healthy, CIA, CIA + TR-Ab19, and Trem2⁻/⁻ CIA joints. Colors indicate scaled GSVA scores (per-row normalization as shown). Gene-set definitions and computational parameters are described in Supplementary Methods. Related to Fig. 6A–G and Fig. S17–S21. Abbreviations: GSVA, gene set variation analysis; CIA, collagen-induced arthritis.

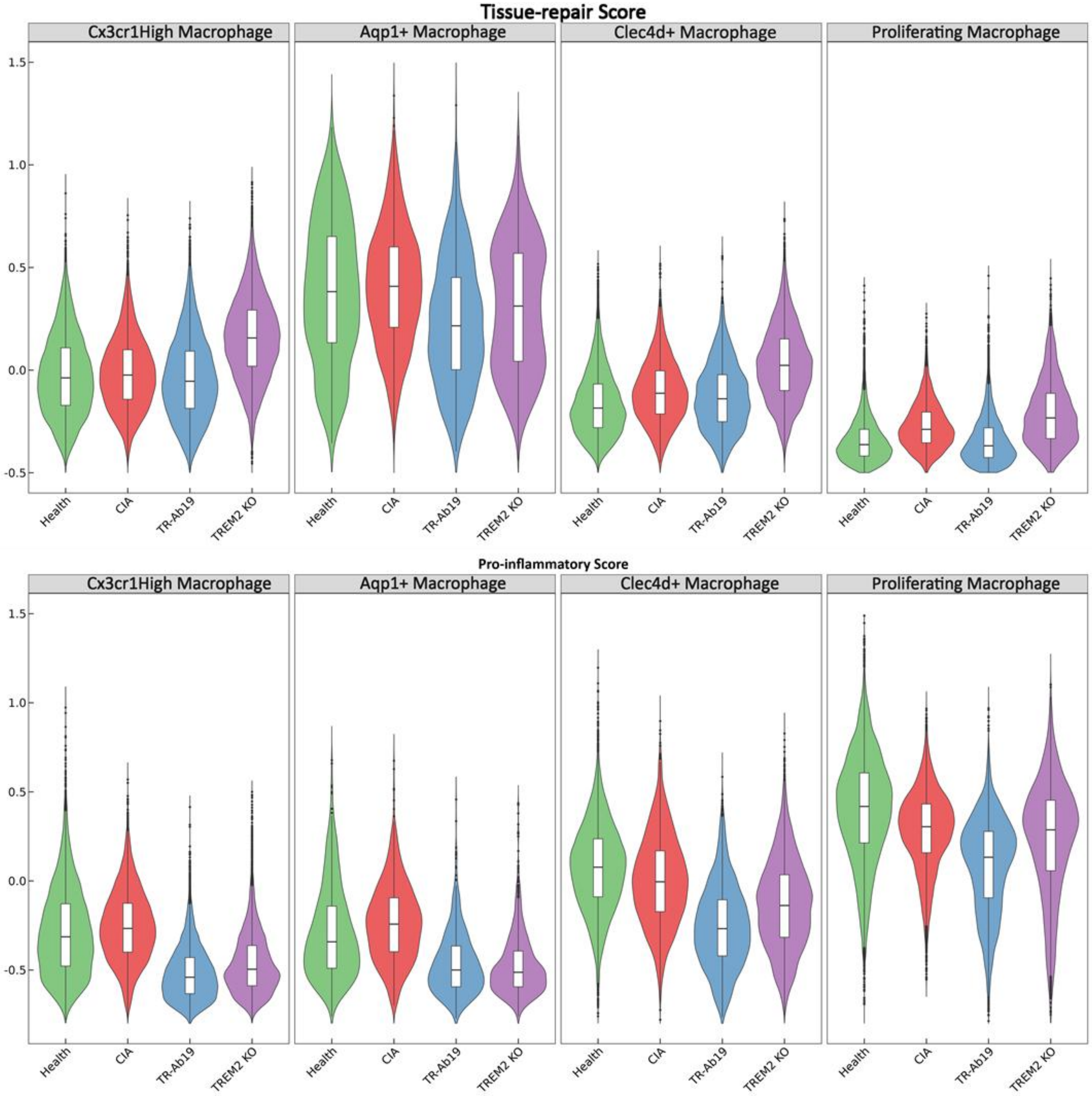


**Figure S23. Module-score analyses support TR-Ab19–driven rebalancing from inflammatory to tissue-repair programs in synovial macrophages.** Top: dot plot of representative marker genes across the four macrophage subsets defined in Fig. 6 ($CX_3CR1^{high}$, $Aqp1^+$, $Clec4d^+$, and proliferating). Dot size indicates the fraction of cells expressing each gene, and dot color indicates average log-normalized expression. Middle: violin plots of the barrier/tissue-repair module score across macrophage subsets and conditions (Healthy, CIA, CIA + TR-Ab19, and $Trem2^{-}/^{-}$ CIA). Bottom: violin plots of the pro-inflammatory module score across subsets and conditions. Related to Fig. 6H–I. Abbreviations: CIA, collagen-induced arthritis.

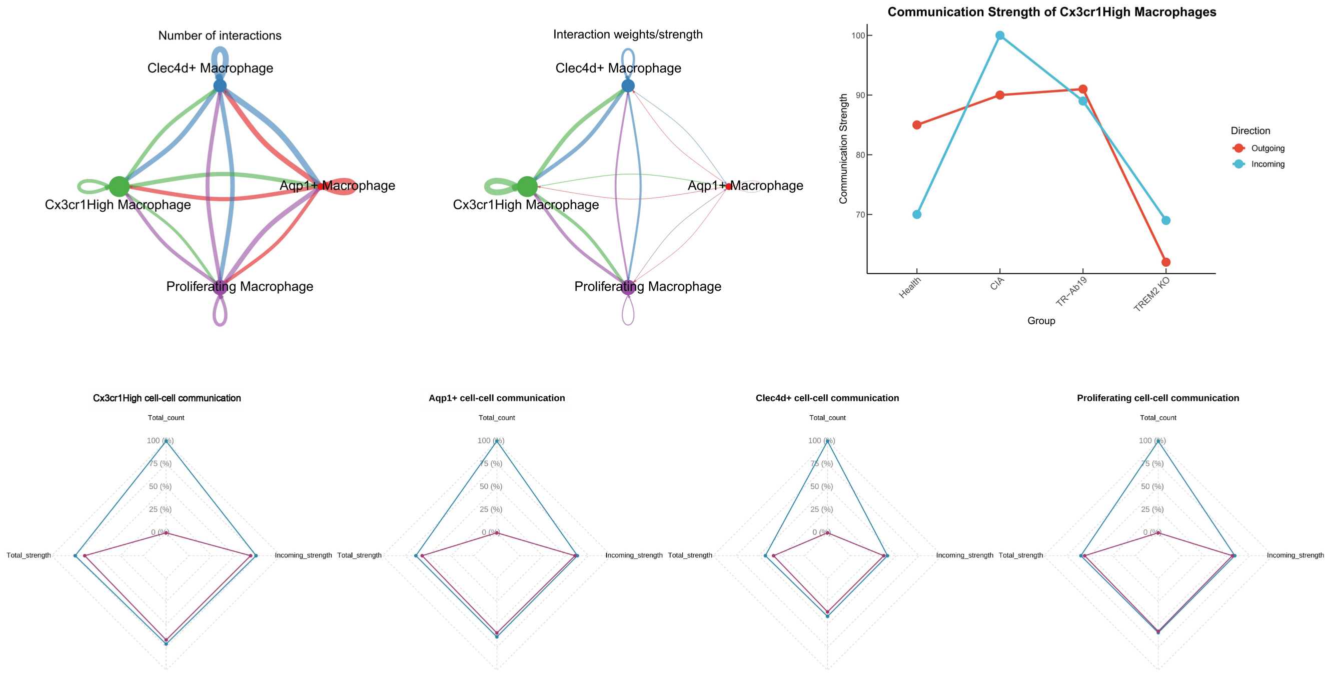


**Figure S24. CellChat inference of inter-subset communication among synovial macrophage states across conditions.** Cell–cell communication networks were inferred using CellChat with default filtering and the curated ligand–receptor database as described in Supplementary Methods. Circle plots summarize (i) the number of inferred ligand–receptor interactions ("interaction counts") and (ii) the summed interaction strength ("interaction strength") among the four macrophage subsets ($CX_3CR1^{high}$, $Aqp1^+$, $Clec4d^+$, and proliferating) under indicated conditions. Line/radar plots summarize outgoing and incoming communication strengths (and derived ratios/summary metrics as labeled) across Healthy, CIA, CIA + TR-Ab19, and $Trem2^{-}/^{-}$ CIA joints. Related to Fig. 6J. Abbreviations: CIA, collagen-induced arthritis.

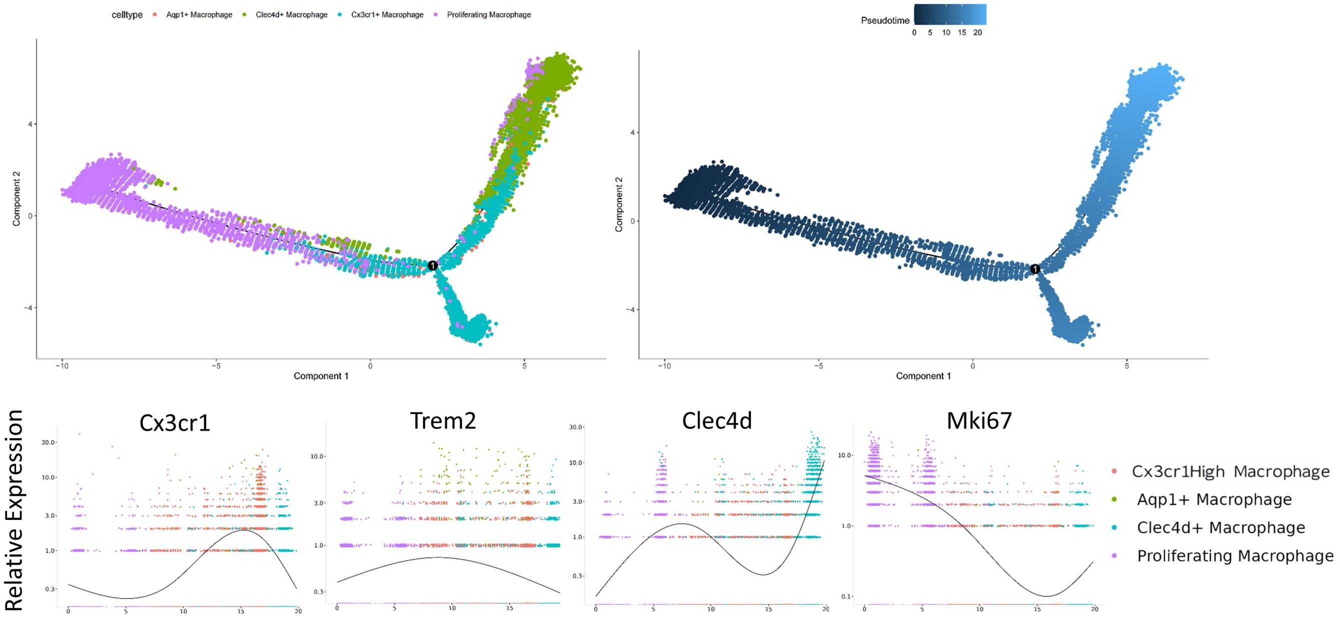


**Figure S25. Pseudotime trajectories of synovial macrophage states in the integrated dataset.**

Monocle 2 trajectory reconstruction (DDRTree) of integrated synovial macrophages from Healthy, CIA, CIA + TR-Ab19, and $Trem2^{-/-}$ CIA joints, colored by macrophage subset and by inferred pseudotime (early to late as shown). Smoothed gene-expression trends along pseudotime for representative markers (e.g., $CX_3CR1$, Trem2, Clec4d, Mki67) are shown as indicated. Trajectory inference parameters and preprocessing are described in Supplementary Methods. Related to Fig. 6 and Fig. S26. Abbreviations: CIA, collagen-induced arthritis.

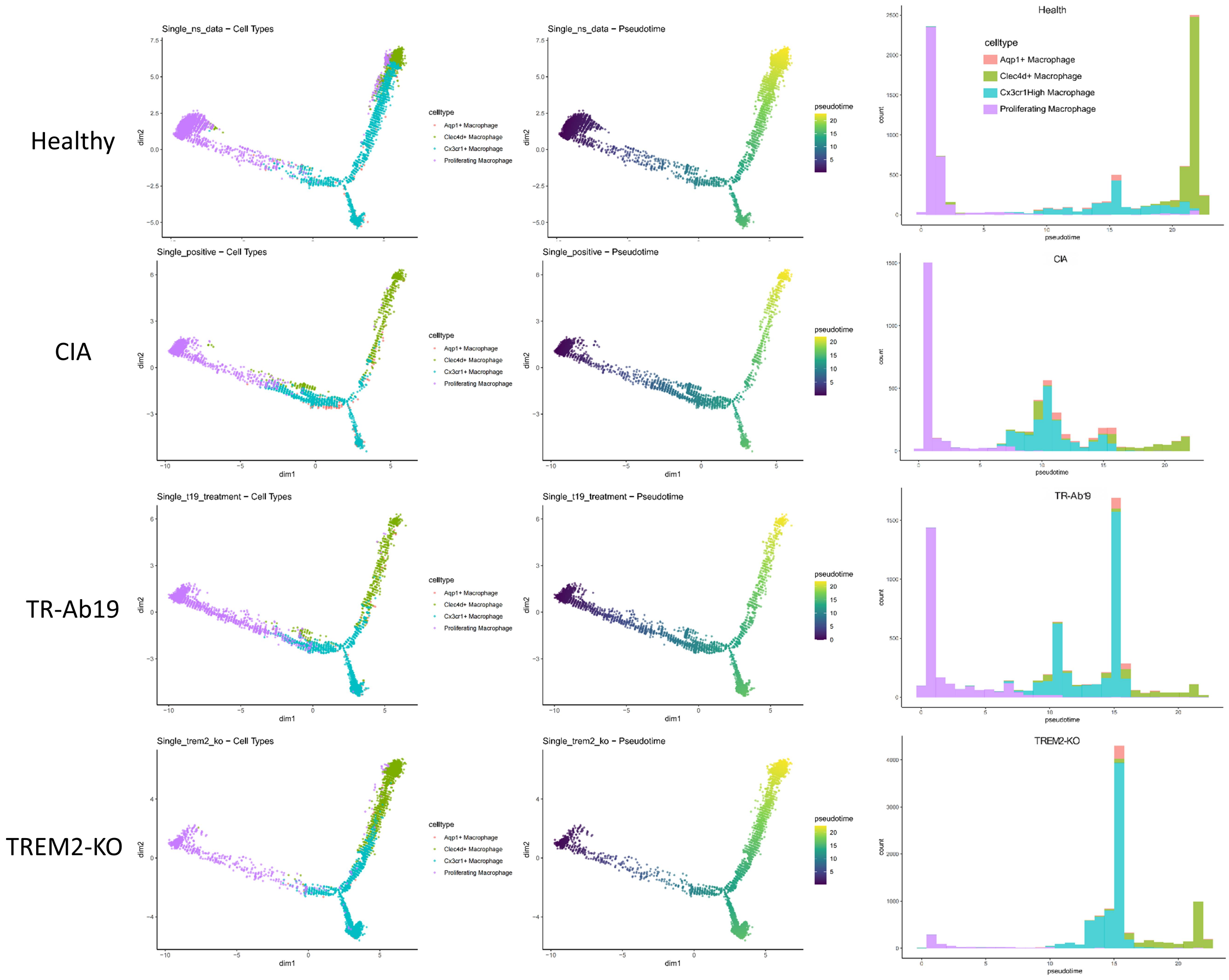


**Figure S26. Condition-specific pseudotime distributions of synovial macrophage states.** Monocle 2 trajectory reconstructions of synovial macrophages stratified by condition (Healthy, CIA, CIA + TR-Ab19, and Trem2$^{-}$/$^{-}$ CIA). For each condition, DDRTree embeddings are shown colored by macrophage subset and by inferred pseudotime. Stacked distributions summarize pseudotime for each subset as indicated. Trajectory inference parameters and preprocessing are described in Supplementary Methods. Related to Fig. 6 and Fig. S25. Abbreviations: CIA, collagen-induced arthritis.

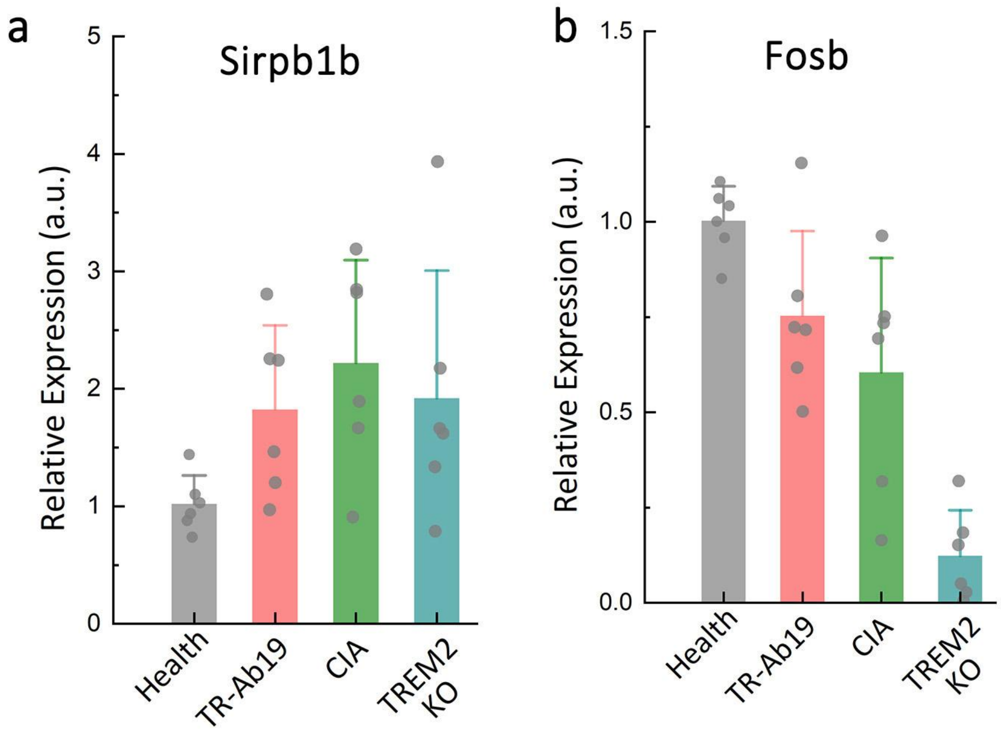


**Figure S27. qRT–PCR validation of selected TREM2-related genes in synovial tissue.** Relative mRNA expression of (A) Sirpb1b and (B) Fosb in synovial samples from Healthy, CIA, CIA + TR-Ab19, and Trem2⁻/⁻ CIA mice. RNA isolation, reverse transcription, cycling conditions, and primer sequences are described in Supplementary Methods and Table S1. Expression was quantified by qRT–PCR and reported as relative expression (arbitrary units) normalized to housekeeping gene(s) used in the experiment (Supplementary Methods). Each dot represents one mouse; bars indicate mean ± SEM; n values are indicated in the plots. Statistical analysis was performed in GraphPad Prism as described (one-way ANOVA with Tukey's multiple-comparisons test for parametric data, or Kruskal–Wallis with Dunn's test for nonparametric data; $*P < 0.05$, $**P < 0.01$, $***P < 0.001$; n.s., not significant). Related to Fig. 7. Abbreviations: qRT–PCR, quantitative reverse transcription PCR; CIA, collagen-induced arthritis.

**Table S1 Primers used for qPCR characterizing $CX_3CR1$ macrophages.**

| | Forward primer | Reverse primer |
|---|---|---|
| fosB | GACCCCTTCCCCGTTGTTAG | CCTTGTTCCTTGCGGGTTTG |
| STAT2 | GTCCTTGAACCGCTTGGAGA | TTGGCAGGATGCTCTGTGAG |
| Sirpb1a | CTGTCCCTCAGCAGACAGTG | AGAGCGGACATCCCTAGGTT |
| Sirpb1b | CAGCCAGAGCTGTCCCTAAG | CCCTAGGTTCCAACACCACC |
| Sirpb1c | CCTGGACCCACATTCCTCAC | ACCCTCCAGCACCAACAAAA |
| SOCS3 | TGTCGGAAGACTGTCAACGG | CCGTTGGGGCTGGATTTTTG |
| Trem2 | CTGGAACCGTCACCATCACTC | CGAAACTCGATGACTCCTCGG |
| Itpr3 | GGGCGCAGAACAACGAGAT | GAAGTTTTGCAGGTCACGGTT |
